# The genome and population genomics of allopolyploid *Coffea arabica* reveal the diversification history of modern coffee cultivars

**DOI:** 10.1101/2023.09.06.556570

**Authors:** Jarkko Salojärvi, Aditi Rambani, Zhe Yu, Romain Guyot, Susan Strickler, Maud Lepelley, Cui Wang, Sitaram Rajaraman, Pasi Rastas, Chunfang Zheng, Daniella Santos Muñoz, João Meidanis, Alexandre Rossi Paschoal, Yves Bawin, Trevor Krabbenhoft, Zhen Qin Wang, Steven Fleck, Rudy Aussel, Laurence Bellanger, Aline Charpagne, Coralie Fournier, Mohamed Kassam, Gregory Lefebvre, Sylviane Métairon, Déborah Moine, Michel Rigoreau, Jens Stolte, Perla Hamon, Emmanuel Couturon, Christine Tranchant-Dubreuil, Minakshi Mukherjee, Tianying Lan, Jan Engelhardt, Peter Stadler, Samara Mireza Correia De Lemos, Suzana Ivamoto Suzuki, Ucu Sumirat, Wai Ching Man, Nicolas Dauchot, Simon Orozco-Arias, Andrea Garavito, Catherine Kiwuka, Pascal Musoli, Anne Nalukenge, Erwan Guichoux, Havinga Reinout, Martin Smit, Lorenzo Carretero-Paulet, Oliveiro Guerreiro Filho, Masako Toma Braghini, Lilian Padilha, Gustavo Hiroshi Sera, Tom Ruttink, Robert Henry, Pierre Marraccini, Yves Van de Peer, Alan Andrade, Douglas Domingues, Giovanni Giuliano, Lukas Mueller, Luiz Filipe Pereira, Stephane Plaisance, Valerie Poncet, Stephane Rombauts, David Sankoff, Victor A. Albert, Dominique Crouzillat, Alexandre de Kochko, Patrick Descombes

## Abstract

*Coffea arabica*, an allotetraploid hybrid of *C. eugenioides* and *C. canephora*, is the source of approximately 60% of coffee products worldwide, and its cultivated accessions have undergone several population bottlenecks. We present chromosome-level assemblies of a di-haploid *C. arabica* accession and modern representatives of its diploid progenitors, *C. eugenioides* and *C. canephora*. The three species exhibit largely conserved genome structures between diploid parents and descendant subgenomes, with no obvious global subgenome dominance. We find evidence for a founding polyploidy event 350,000-610,000 years ago, followed by several pre-domestication bottlenecks, resulting in narrow genetic variation. A split between wild accessions and cultivar progenitors occurred ∼30.5 kya, followed by a period of migration between the two populations. Analysis of modern varieties, including lines historically introgressed with *C. canephora*, highlights their breeding histories and loci that may contribute to pathogen resistance, laying the groundwork for future genomics-based breeding of *C. arabica*.

## Introduction

Polyploidy is a powerful evolutionary force that has shaped genome evolution across many eukaryotic lineages, possibly offering adaptive advantages in times of global change^1,2^. Such whole genome duplications (WGDs) are particularly characteristic of plants^3^, and a great proportion of crop species are polyploid^4-11^. Our understanding of genome evolution following WGD is still incomplete, but outcomes can result in genomic shock, in terms of activation of cryptic transposable elements, subgenome-partitioned gene regulation or fractionation, homoeologous exchange, meiotic instability, and even karyotype variation ^8, 12-16^. Alternatively, few or none of the above phenomena can materialize, and the two genomes can coexist harmonically, gradually adaptating to new ploidy levels^17^. Regardless, the most common fate of polyploids appears to be fractionation and eventual reversion to the diploid state^18^.

With an estimated production of 10 million metric tons per year, coffee is one of the most traded commodities in the world. The most broadly appreciated coffee is produced from the allotetraploid species *Coffea arabica*, especially from cultivars belonging to the Bourbon or Typica lineages and their hybrids^19^. *C. arabica* (2n = 4x = 44 chromosomes) resulted from a natural hybridization event between the ancestors of present-day *C. canephora* (Robusta coffee, subgenome CC) and *C. eugenioides* (subgenome EE; each with 2n = 2x = 22). The founding WGD has been dated to between 10,000 to one million years ago^20-23^, with the Robusta-derived subgenome of *C. arabica* closest related to *C. canephora* accessions from northern Uganda^24^. Arabica cultivation was initiated in 15^th^ -16^th^ century Yemen. (**Ext. data Fig. 1**). Around 1600, “seven seeds” were smuggled out of Yemen^25^, establishing Indian *C. arabica* cultivar lineages. A century later, the Dutch began cultivating Arabica in Southeast Asia – thus setting up the founders of the contemporary Typica group. One plant, shipped to Amsterdam in 1706, was used to establish Arabica cultivation in the Caribbean in 1723. Independently, the French cultivated Arabica on the island of Bourbon (presently Réunion)^26^, and the descendants of a single plant that survived by 1720 form the contemporary Bourbon group. Contemporary Arabica cultivars descend from these Typica or Bourbon lineages, except for a few wild ecotypes with origins in natural forests in Ethiopia. Due to its recent allotetraploid origin and strong bottlenecks during its history, cultivated *C. arabica* harbors a particularly low genetic diversity^20^ and is susceptible to many plant pests and diseases, such as coffee leaf rust (*Hemileia vastatrix*). As a result, the classic Bourbon-Typica lineages can only be cultivated successfully in a few regions around the world. Fortunately, a spontaneous *C. canephora* x *C. arabica* hybrid resistant to *H. vastatrix* was identified on the island of Timor^27^ in 1927. Many modern Arabicas contain *C. canephora* introgressions derived from this hybrid, ensuring rust resistance, but having also unwanted side effects, such as decreased beverage quality^28^.

**Figure 1.**
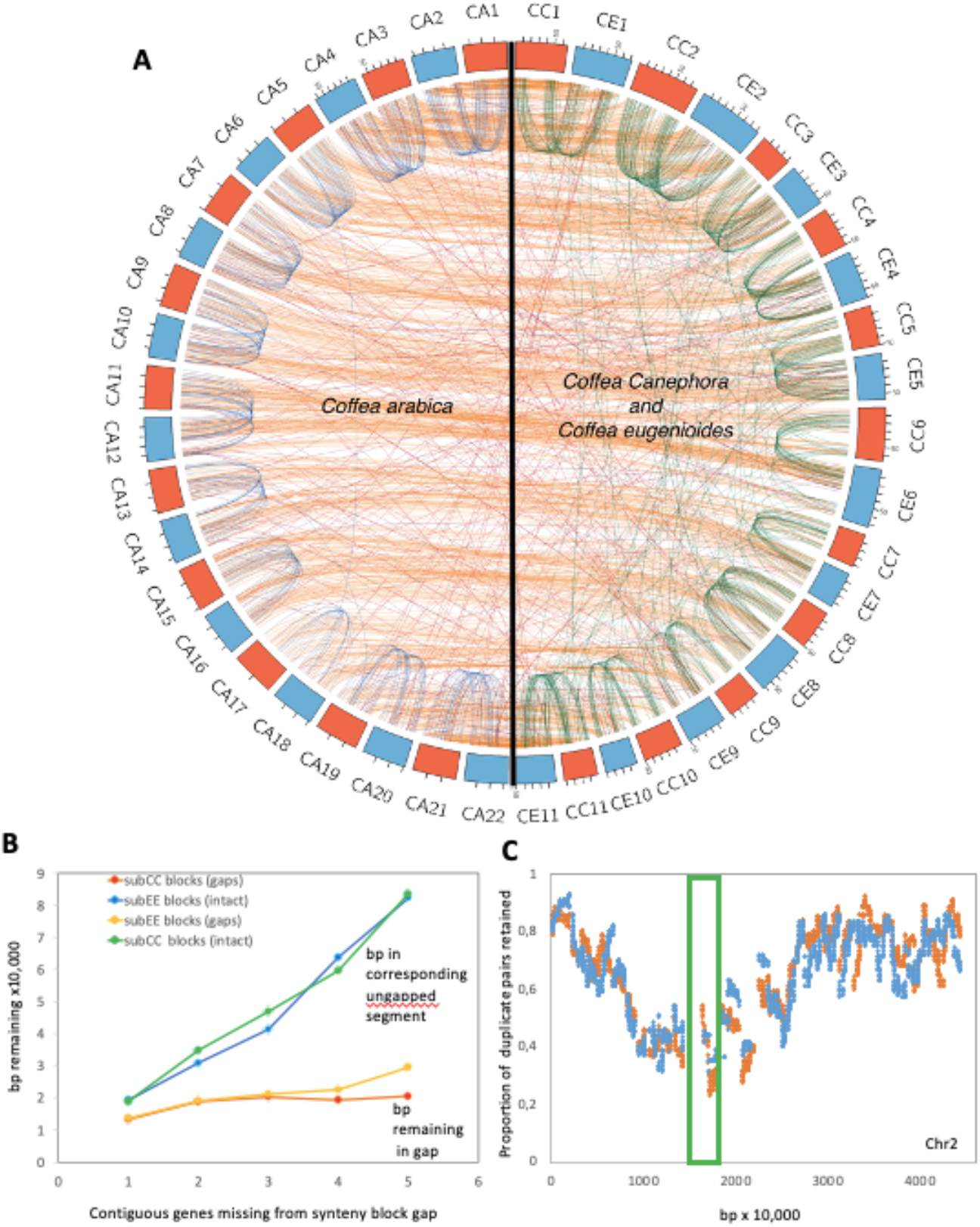
Patterns of synteny, fractionation and gene loss in Coffea arabica (CA) and its progenitor species C. canephora (CC) and C. eugenioides (CE). **A**. Corresponding syntenic blocks between CA subgenomes subCC (orange) and subEE (blue), and with the CC (orange) and CE (blue) genomes. **B**. bp in intergenic DNA in synteny block gaps caused by fractionation in a subCC-subEE comparison, compared to numbers of bp in homoeologous unfractionated regions, as a function of numbers of consecutive genes deleted. **C**. Gene retention rates in synteny blocks plotted along subCC chromosome 2; subCC is plotted in orange and subEE in blue. The green box indicates the pericentromeric region.

Modern genomic tools and a detailed understanding of the origin and breeding history of contemporary varieties are vital to developing new Arabica cultivars, better adapted to climate change and agricultural practices^29-31^. Here, we present chromosome-level assemblies of *C. arabica* and representatives of its progenitor species, *C. canephora* (Robusta) and *C. eugenioides* (hereafter Eugenioides). Whole-genome resequencing data of 41 wild and cultivated accessions facilitated in-depth analysis of Arabica history and dissemination routes, as well as the identification of candidate genomic regions associated with pathogen resistance.

## Results and Discussion

### Chromosome-level assemblies and annotations, genome fractionation, and subgenome dominance

As reference individuals, we chose the di-haploid Arabica line ET-39^32^, a previously sequenced doubled haploid Robusta^33^, and the wild Eugenioides accession Bu-A, respectively. Long and short-read-based hybrid assemblies were obtained (Online methods and **Supplementary sections 2.1-2.2**), spanning 672 Mb (Robusta), 645 Mb (Eugenioides) and 1,088 Mb (Arabica), respectively. Upon HiC scaffolding, the Robusta and Arabica assemblies consisted of 11 and 22 pseudochromosomes, and spanned 82.7% and 62.5%, respectively, of the projected genome sizes (**Table 1**). To improve the Arabica assembly, we generated a second assembly using PacBio HiFi technology followed by Hi-C scaffolding (Online methods and **Supplementary section 2.3**). This assembly was 1,198 Mb long, of which 1,192 Mb (93.1% of the predicted genome size based on cytological evidence^34^) was anchored to pseudochromosomes (**Table 1**). Gene space completeness, assessed using Benchmarking Universal Single-Copy Orthologs (BUSCO)^35^, was >96% for all assemblies. Importantly, 93.2% of the BUSCO genes were duplicated in the HiFi assembly (**Table 1**), indicating that most of the gene duplicates from the allopolyploidy event were retained.

**Table 1.**
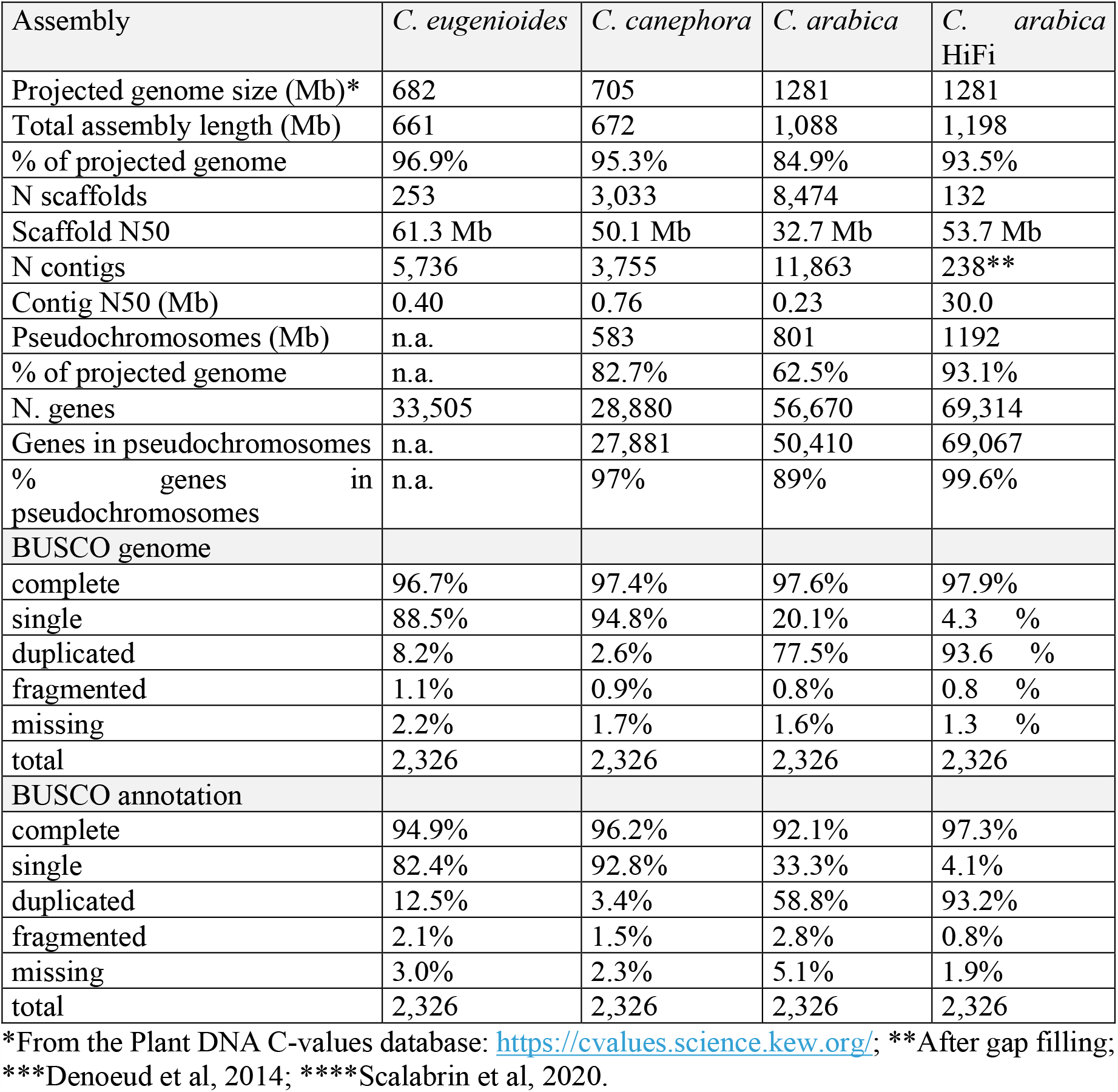
Statistics of the Coffea assemblies presented in this paper.

The Robusta and Eugenioides genomes contained, respectively, 67.5% and 59.7% transposable elements (TEs) (**Supplementary section 3.2**), with Gypsy LTR retrotransposons accounting for most of the difference between the two species. This difference was greatly reduced (63.1% and 63.8%) in the two Arabica subgenomes (subCC and subEE, stemming from Robusta and Eugenioides ancestors, respectively), possibly indicating TE transfer via homoeologous exchange. Robusta contained considerably more recent LTR TE insertion elements than Eugenioides. Again, the two Arabica subgenomes showed greater similarity to each other in recent LTR TE insertions than the two progenitor genomes. No major evidence was found for LTR TE mobilization following Arabica allopolyploidization, in contrast to what has been observed in tobacco^36^, but similar to *Brassica* synthetic allotetraploids^37^. Arabica genome evolution observed instead more closely follows the “harmonious coexistence” pattern^38^ seen in *Arabidopsis* hybrids^17,39^.

High-quality gene annotations, followed by manual curation of specific gene families (**Supplementary Sections 3.1-3.4**), resulted in 28,857, 33,505, 56,670, and 69,314 gene models for the Robusta, Eugenioides, PacBio Arabica, and Arabica HiFi assemblies, respectively (**Table 1**). Altogether ∼97% of Robusta and 99.6% of Arabica HiFi gene models were placed on the pseudochromosomes, with 33,618 and 35,449, respectively, to subgenomes subCC and subEE (**Table 1**). Annotation completeness from BUSCO was ≥95% for Eugenioides and Robusta, and reached 97.3% for Arabica HiFi.

Comparison of Arabica subCC and subEE against their Robusta and Eugenioides counterparts revealed high conservation in terms of chromosome number, centromere position and numbers of genes per chromosome (**Fig. 1, Supplementary section 4**). Patterns of gene loss following the *gamma* paleohexaploidy event displayed high structural conservation between Robusta and Eugenioides during the 4-6 million years (My) since their initial species split^22,23^ (**Supplementary section 4**). Likewise, the structures of the two Arabica subgenomes were highly conserved between each other, with, since the Arabica-founding allotetraploidy event, only ∼5% of BUSCO genes having reverted to the diploid state (**Fig. 1A; Table 1**). Syntenic comparisons revealed that genomic excision events, removing one or several genes at a time in similar proportions across the two subgenomes, have been the main driving force in genome fragmentation both before and after the polyploidy event (**Fig. 1B, Supplementary section 4**). Fractionation occurred mostly in pericentromeric regions, whereas chromosome arms showed more moderate paralogous gene deletion (**Fig 1C, Supplementary section 4**). The Arabica allopolyploidy event seemingly did not affect the rate of genome fractionation, which remained roughly constant when comparing deletions in progenitor species versus Arabica subgenomes after the event. In support of the dosage-balance hypothesis^40^, subgenomic regions with high duplicate retention rates were significantly enriched for genes that originated from the Arabica WGD (Fisher exact test, *p*<2.2e-16). In contrast, low duplicate retention rate regions significantly overlapped with genes originating from small-scale (tandem) duplications (**Table S1**). Genes with high retention rates were enriched in Gene Ontology (GO) categories such as “cellular component organization or biogenesis”, “primary metabolic process”, “developmental process” and “regulation of cellular process”, while low retention rate genes were enriched in categories such as “RNA-dependent DNA biosynthetic process” and “defense response” (in both subgenomes) and “spermidine hydroxycinnamate conjugate biosynthetic process” (involved in plant defense^41^) and “plant-type hypersensitive response” (in subEE) (**Tables S2-S5**).

To study possible expression biases between subgenomes, we identified syntelogous gene pairs and removed the pairs showing homoeologous exchanges in the Arabica subgenomes (see under **Origin and domestication of Arabica coffee**, below)^42^ (**Supplementary section 5**). Overall, no significant global subgenome expression dominance was observed (**Tables S6-S7**). However, gene families regularly displayed mosaic patterns of expression, including several encoding enzymes that contribute to cup quality, such as *N*-methyltransferase (*NMT*), terpene synthase (*TPS*), and fatty acid desaturase 2 (*FAD2*) families, all having some genes being more expressed in one of the two subgenomes (**Ext. data Fig. 2**), as per a recent study^43^. Similar gene family-wise patterns occur in other evolutionarily recent polyploids such as rapeseed^10^ and cotton^44^, which are also at their early stages of transitioning back to a diploid state.

**Figure 2.**
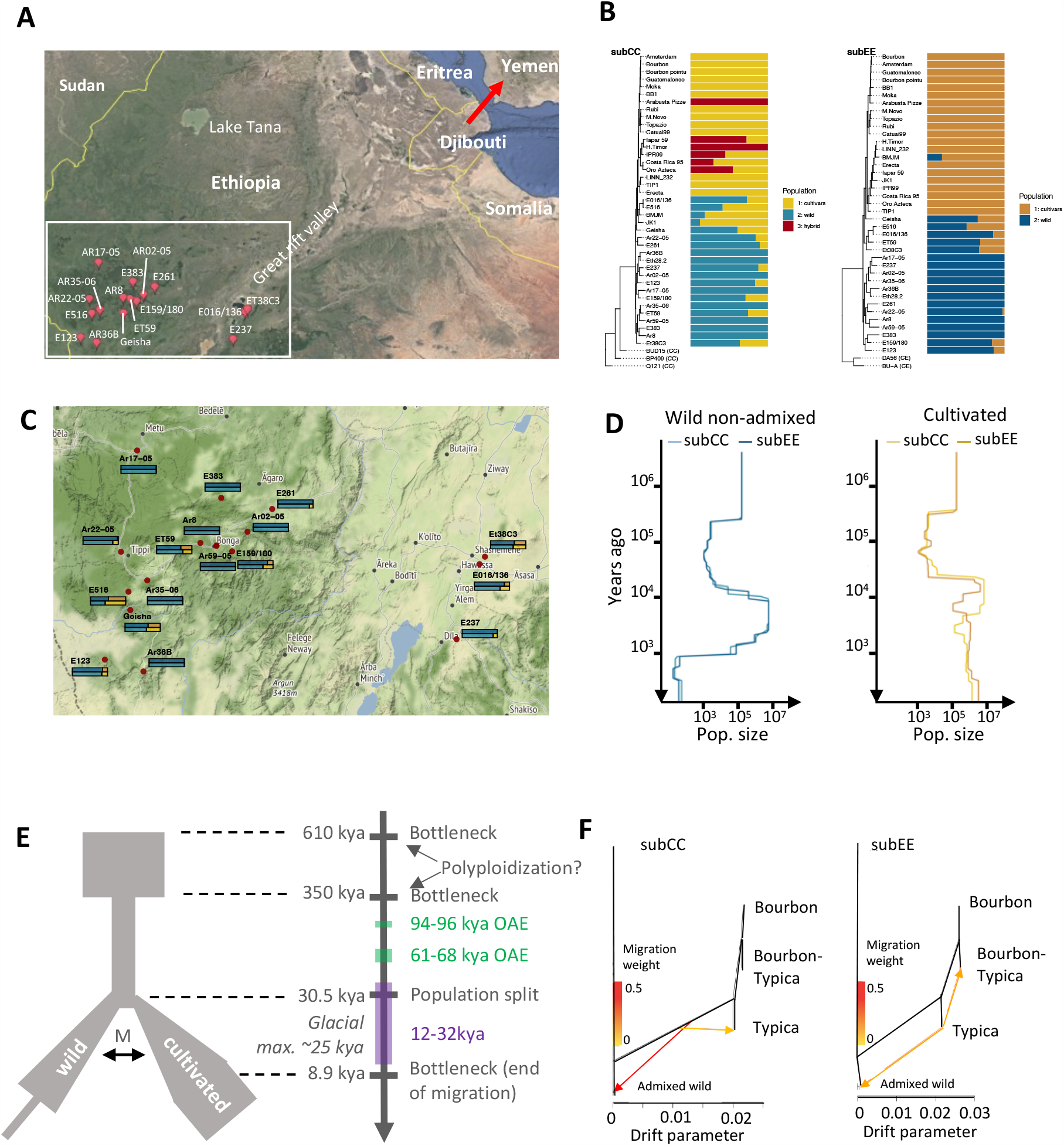
Population history of Coffea arabica. **A**. Geographic origin of resequenced wild C. arabica accessions. The red arrow indicates the probable route of migration to Yemen in historical times. **B**. Ancestral population assignments of C. arabica accessions for subCC (left) and subEE (right). Relationships among individuals are illustrated with phylogenetic trees obtained from independent SNPs. For magnified views of the trees, see **Fig. S45. C**. Magnification of panel A, showing the admixture values for each of the accessions in subCC (top) and subEE (bottom); the colors correspond to the analysis in panel B. **D**. Population sizes of wild and cultivated accessions, inferred using SMC++, suggest genetic bottlenecks at ∼350 and 1 kya (limited to non-admixed wild individuals). **E**. FastSimcoal2 output, suggesting a population split ∼30.5 kya, followed by a period of migration between the populations until ∼8.9 kya. This timing corresponds with increased population diversity in cultivars at a similar time, calculated using SMC++. Green rectangles along the timeline show “windows of opportunity”, times when Yemen was connected with the African continent wherein human migrations to the Arabian Peninsula may have occurred. The purple rectangle shows the last ice age. **F**. Directional gene flow analysis using Orientagraph suggests two hypotheses: gene flow from the shared ancestral population of all cultivars to the Ethiopian wild individuals (subCC), or gene flow from the Typica lineage to Ethiopia (subEE).

### Origin and domestication of Arabica coffee

To obtain a genomic perspective on the evolutionary history of Arabica, we sequenced 46 accessions, including three Robusta, two Eugenioides, and 41 Arabica. The latter included an 18th-century type specimen, kindly provided by the Linnaean Society of London, 12 cultivars with different breeding histories, the Timor hybrid and five of its backcrosses to Arabica, and 17 wild and three wild/cultivated accessions collected from the Eastern and Western sides of the Great Rift Valley^45,46^ (**Table S8, Fig. 2A**).

Homoeologous exchange (HE) between subgenomes has been observed in several recent polyploids^8,42,47^. Arabica generally displays bivalent pairing of homologous chromosomes and disomic inheritance^48^, but since the subgenomes share high similarity, occasional homoeologous pairing and exchange may also occur. We therefore explored the extent of HE among Arabica accessions and its possible contribution to genome evolution. Overall, all accessions shared a fixed allele bias toward subEE at one end of chromosome 7, which contained genes enriched for chloroplast-associated functions (**Ext. data Fig. 3A, Supplementary section 5, Table S9**). Since the Arabica plastid genome is derived from Eugenioides^49^, HE in this region was likely selected for, due to compatibility issues between nuclear and chloroplast genes encoding chloroplast-localized proteins^50^. Surprisingly, all but one accession (BMJM) showed significant (*p*-value <9.8e-37) 3:1 allelic biases towards subCC. The highly concordant HE patterns, present in both wild and cultivated Arabicas (**Ext. data Fig. 4)**, suggested that i) the allelic bias is an adaptive trait not associated with breeding, and ii) it originated in a common ancestor of all sampled accessions, possibly immediately after the founding allopolyploidy event. Some exchanges, shared by only a few accessions, probably originated more recently (**Ext. data Fig. 3B**). More recent HE events were also found in some cultivars and also showed a bias towards subCC, except for BMJM, which showed bias towards subEE due to a single large crossover in chromosome 1 (**Ext. data Fig. 3A**). An interesting hypothesis for future investigation is that in a low-diversity polyploid species such as Arabica, HE could be a major contributor to phenotypic variation observed among closely related accessions^51^.

**Figure 3.**
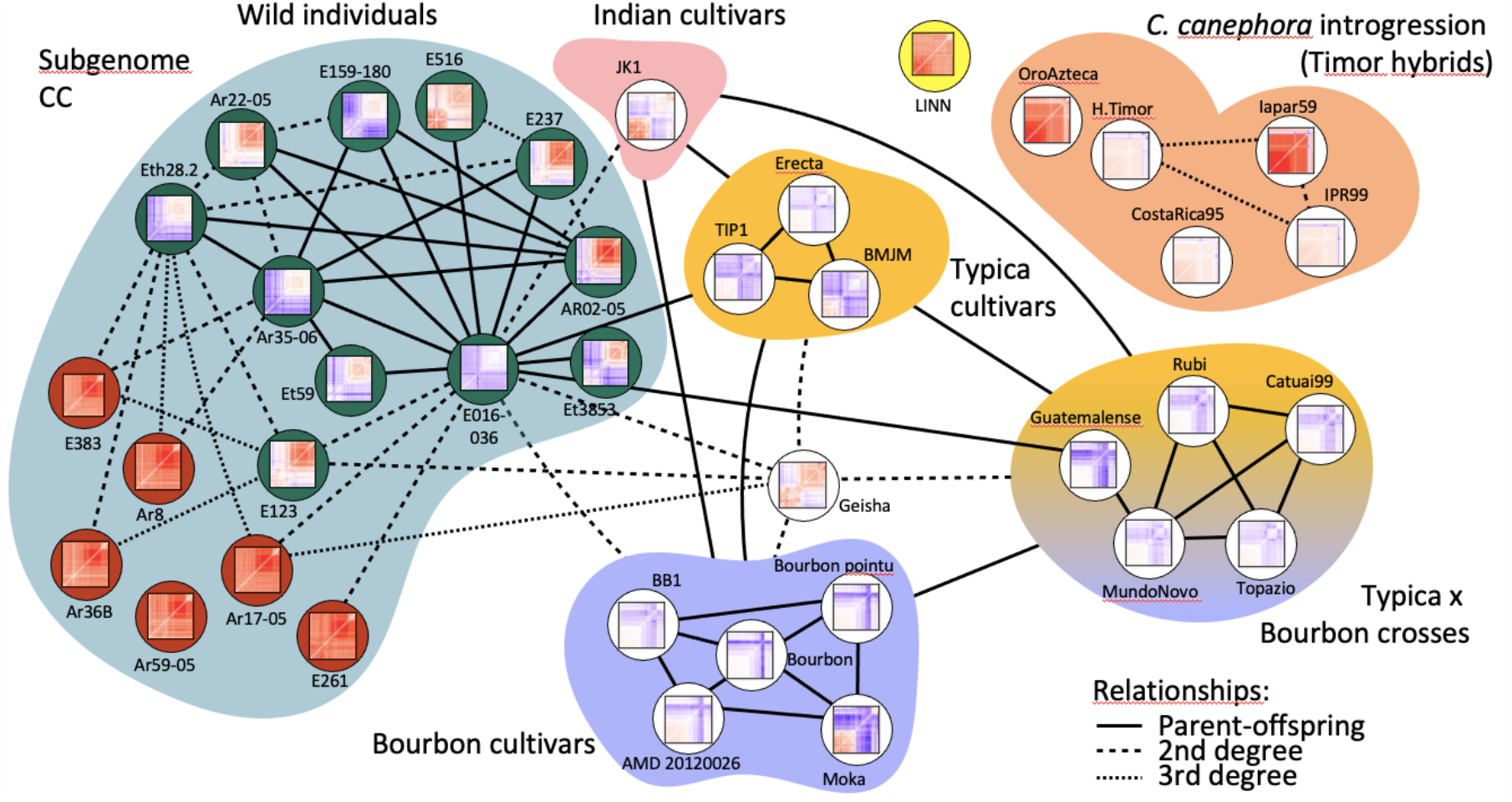
Kinship estimation of C. arabica accessions, inferred from SNPs in subCC. The degree of relatedness was estimated using Kinship-based INference for GWAS (KING) and describes the number of generations between the related accessions. Thumbnail images show false discovery rate corrected F3 tests of introgression Z-statistics for each of the target individuals. Each cell in the matrix illustrates an F3 test result for the target accession containing introgression from two different sources (x- and y-axis); blue color illustrates significant gene flow (or allele sharing via identity by descent^61^; IBD) from the two source accessions to the target, while red color illustrates lack of gene flow. See **Ext. data Fig. 7** for corresponding analyses in subEE. In the wild accessions, the dark green background highlights the admixed individuals (**Fig. 3B**), while the non-admixed individuals are highlighted with red background. Relationships follow standard nomenclature (e.g., 2^nd^ degree refers to an individual’s grandparents, grandchildren etc., whereas 3rd degree refers to great-grandparents, great-grandchildren, etc.)

**Figure 4.**
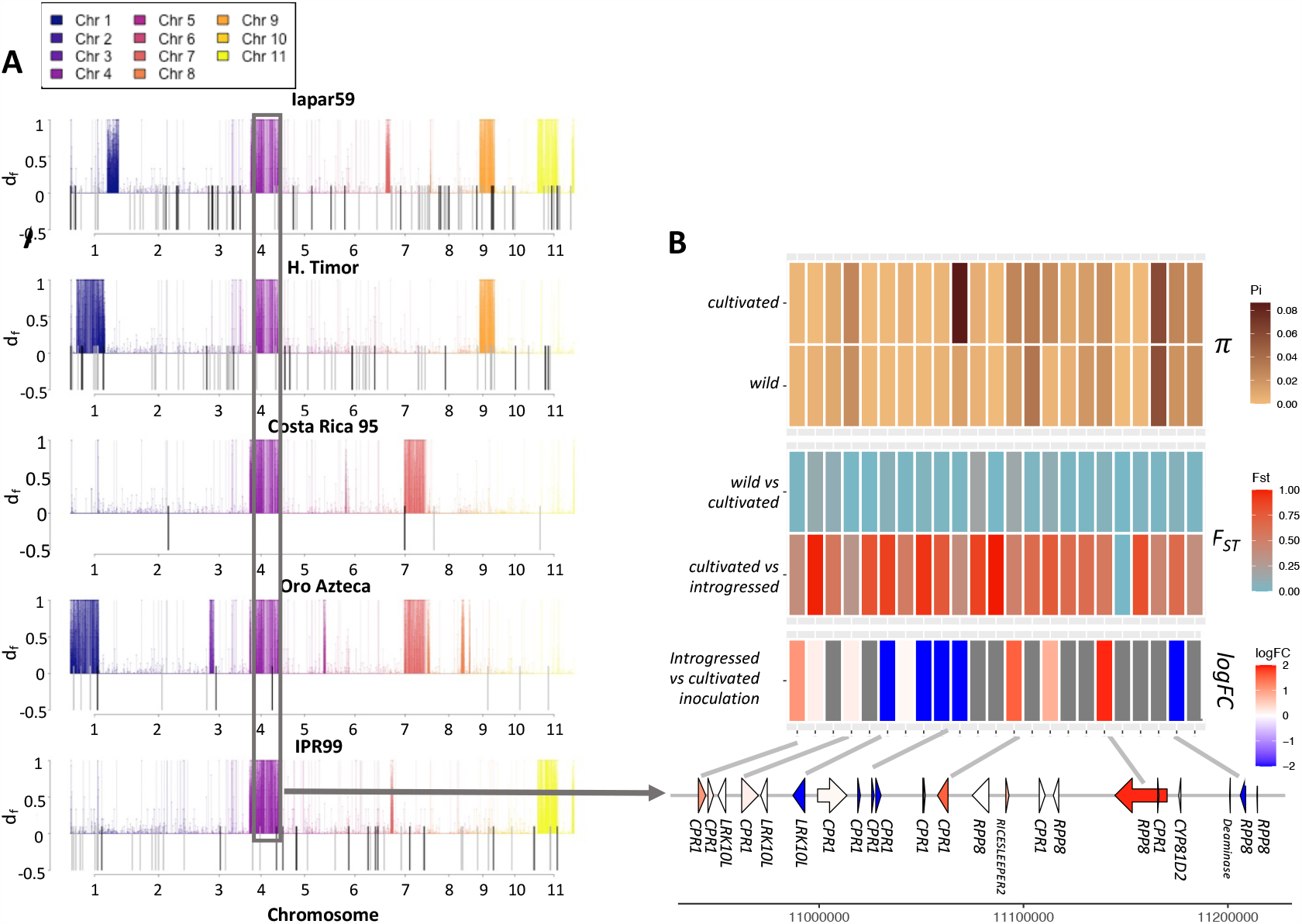
Introgression of Coffea canephora into H. vastatrix-resistant C. arabica lineages. **A**. Introgression d_f_ statistic estimated for different Timor hybrid derivatives. Colored lines above the axis mark regions of significant introgression in the line under inspection, and are colored by chromosome. The shared introgressed region on Chr 4 is colored in purple and boxed. Transposon Insertion Polymorphisms are represented as lines below the X axis and exhibit overlap with introgressed regions. **B**. The shared introgressed genomic region on subCC chromosome 4 contains a cluster of R genes (RPP8), a cluster of homologs of a negative regulator of R genes (CPR1), and a cluster of homologs of leaf rust resistance 10 kinases (LRK10L) (bottom). The heatmap shows, from the bottom up, (i) log fold change of gene expression after H. vastatrix inoculation, when comparing resistant Timor hybrid lineage against a susceptible cultivar; red color means elevated expression in the hybrid, and blue decreased expression. (ii) Fixation index (F_ST_) values for the introgessed lines vs cultivars and between cultivars and wild accessions. (iii) Nucleotide diversity for the wild and cultivated accessions for each gene coding region, plus the flanking 2kb upstream and downstream of the region.

We next studied population genetic statistics for each of the subgenomes (**Table S10**). The 17 wild samples demonstrated low genomic diversities, indicative of small effective population sizes, while negative Tajima’s D suggested an expanding population, possibly following one or more population bottlenecks. The cultivars and wild population samples had similar genetic diversities, as demonstrated by low fixation index (F_ST_) values. In cultivars, nucleotide diversities were only slightly lower than in wild populations and Tajima’s D scores were less negative, suggesting that only minor bottlenecks and subsequent population expansions occurred during domestication.

SNP tree estimation and ADMIXTURE analyses (**Fig. 2B**) identified a three-population solution for subCC: Typica-Bourbon cultivars (Population 1), wild accessions (Population 2), and Timor hybrid-derived cultivars (Population 3). The old BMJM and the recently established Geisha cultivars showed admixed states on both subgenomes, similar to about half of the wild accessions. Indian varieties encompassed both Typica and Bourbon variation, in agreement with previous studies^20^. The Linnaean sample grouped with the cultivars, supporting its hypothesized origin from the Dutch East Indies^25^. A complementary analysis using PCA (**Ext. data Fig. 5)** was in agreement with ADMIXTURE analysis.

In wild accessions, both subgenomes concordantly showed two population bottlenecks (**Fig. 2D**) in the SMC++^52^ modeling. Assuming a 21-year generation time^53^, the oldest bottleneck initiated abruptly around 350 thousand years ago (kya) and ended around 15 kya, at the start of the African humid period (AHP)^54^, when climatic conditions were more favourable for Arabica growth. The more recent bottleneck initiated more gradually around 5 kya and lasts to this day. Cultivated accessions, however, exhibited the oldest, but not the more recent bottleneck. In part due to these differences, we also modeled Arabica population history using FastSimcoal2^55^, modeling the wild population and cultivars as two separate lineages. In the best-fitting model (**Fig. 2E**) the wild population was predicted to split from the cultivar founding population 1,450 generations ago (∼30 kya), i.e., before the last glacial maximum. The original founding event was analyzed using the non-admixed wild individuals, revealing an ancestral population bottleneck at 350 kya (**Ext. data Fig. 6A**). Divergence estimates based on gene fractionation, the distribution of nonsynonymous mutations (**Ext. data Fig. 6B**), and calibrated SNP trees (**Fig. 2B**) suggested the allopolyploid founding event occurred at 610 kya, which is close to previous estimates^22,23^. The 350 kya bottleneck, on the other hand, corresponds to that found in the SMC++ analyses (**Fig. 2D**). We therefore consider 610-350 kya a likely time range for the polyploidization event (**Fig. 2E**). The wild and pre-cultivar lineages maintained some gene flow (in terms of migration) until ∼8-9 kya, which may have contributed to the modeled increase in effective population size (**Fig. 2D-E**).

While these data were not able to identify the precise place of origin of the modern cultivated population (see also the following section), the extended period of migration between wild and cultivated accessions suggests that they were separated only by a relatively small geographic distance, such as along the two sides of the African Great Rift Valley (**Fig. 2A-C**). It is also possible that the cultivated lineage could have extended as far as Yemen, and that the end of migration between the two populations could have been caused by the widening of the Bab al-Mandab strait (separating Yemen and Africa) due to rising sea levels^56^ at the end of the AHP. A native Arabica population exists in Yemen^57^, which could support this hypothesis. The Linnaean sample, together with the Typica and Bourbon cultivars, originate from this second population that was also used to establish cultivation in Yemen, as suggested by the SNP, ADMIXTURE, and PCA analyses (**Fig. 2B, Ext. data Fig. 5**).

In conclusion, our analyses suggest that the Arabica allopolyploidy event occurred between 610 and 350 kya, when considering that inbreeding present in *Coffea* populations would accelerate coalescence estimation^58,59^. Earlier work proposing more recent timings, such as 20 kya^20^, could be underestimates stemming from confounding effects of population bottlenecks in cultivated and wild lineages.

### Origin of modern cultivars

The known breeding history of several of our Arabica cultivars provided us with a gold standard set for deducing the Arabica pedigree using Kinship-based INference for Gwas (KING)^60^ (**Fig. 3**). The method correctly identified the relationships between Bourbon and Typica group cultivars and the Bourbon-Typica crosses in subCC. In contrast, the subEE pedigree showed lower (2^nd^) order relationships, possibly due to HE in that subgenome (**Ext. data Fig. 7**). Timor hybrid-derived accessions did not show significant relationships to mainline cultivars in subCC (likely due to Robusta introgressions in this subgenome that broke the haplotype blocks, see below), while subEE showed 2^nd^ degree relationships to both the Typica and Bourbon groups (**Fig. 3; Ext. data Fig. 7**), confirming that subEE has not received substantial introgression.

Interestingly, Typica, Bourbon and JK1 individuals were also 1^st^ degree related, suggesting direct parent-offspring relationships. Besides confirming their shared Yemeni origins, this finding also underscores the Yemeni germplasm’s limited genetic diversity. Further, the old cultivar lines JK1 (Indian), Erecta (Indonesian Typica), BMJM (Caribbean Typica), TIP1 (Brazilian Typica), and BB1 (Brazilian Bourbon) showed 2^nd^ or higher degree relationships with a cluster of closely related wild admixed accessions, centered on E016/136 (**Fig. 2B**). The recently established Geisha cultivar showed similar relationships to the wild admixed individuals and the Bourbon and Typica groups, suggesting common origins. Interestingly, admixed wild accession E016/136 was closely related to both wild and cultivated populations.

In a comparison of geographic origins, wild individuals from the Eastern side of the Great Rift Valley had some levels of admixture and were closely interrelated, while on the Western side, the admixed, related individuals were mostly concentrated around the Gesha region (**Fig. 2C, Fig. 3**). The E016/136 admixed accession, closest to cultivars, demonstrated a first-degree relationship with several wild accessions, of which only Ar35-06 and Eth28.2 were pure representatives of the wild population (**Fig. 2B**). Therefore, these two accessions are genetically closest, in our sample, to the hypothetical true wild parent of cultivated Arabica, with E016/136 representing an intermediate form. Ar35-06 was collected near Gesha mountain, close to the origin of the modern Geisha cultivar. Altogether these data point to the Gesha region as a hotspot of wild accessions amenable to domestication.

Admixed wild samples may have originated from a recent hybridization event that occurred before or after their collection from the wild. A third alternative is that the Yemeni population (and hence the cultivars) originated from an admixed population from the Eastern side of the Great Rift Valley or the Gesha region. Analysis of admixture patterns with Orientagraph^62^ (**Fig. 2F**) suggested hybridization with the common ancestor of the Bourbon and Typica lineages in subCC, and of Typica in subEE. In the case of recent hybridization, introduced haplotypes would exist as long contiguous blocks (as in the Timor hybridization, which occurred 100 years ago), while for older events the blocks would be more fragmented due to crossing over. Analysis using the distance fraction (d_f_) statistic^63^ showed the latter to be the case (**Ext. data Fig. 8**, indicating that admixture events among wild accessions were not very recent, supporting our third hypothesis.

Domestication and cultivation usually involve strong population bottlenecks based on high wild diversity, resulting in reduced genetic diversity in cultivars^64^. However, Arabica nucleotide diversity was already very low in the wild, probably as a result of earlier bottlenecks (**Figs. 2D-E**), but only marginally reduced in the pre-cultivated lineage (**Ext. data Fig. 9A**). Bourbon had lower diversity than Typica, probably resulting from the known single-individual bottleneck in this group. Also, the inbreeding coefficients in the wild and cultivated accessions were similar (**Ext. data Fig. 9B**), differing from general expectations for a domesticated species^64^.

To look for pathways under purifying selection in cultivars, we identified genes with high *F*_*ST*_ (95 % quantile) between cultivars and wild accessions. This resulted in a set of 1,908 genes that were enriched for the GO categories “cellular response to nitrogen starvation”, “regulation of innate immune response” and “regulation of defense response” (**Table S11**), and contained homologs of ammonium transporters *AMT1* and *AMT2*, important for nitrogen uptake in *Coffea*^*65*^, a homolog of the salicylic acid receptor *NONEXPRESSER OF PR GENES 1* (*NPR1*), required in SA signaling and systemic acquired resistance^66^, as well as a homolog of the *Arabidopsis LSU2* gene, previously identified as a hub convergently targeted by effectors of pathogens from different kingdoms^67^. A second screen, focused on genes with a large number of high-impact nonsynonymous mutations shared among cultivars (>40% individuals having the mutation), generated a list of 556 genes that were significantly enriched for only one GO category, “defense response” (**Table S12**). From the 22 genes in this category, 16 were NB-ARC domain-containing resistance (R) genes, and two were members of the leucine-rich repeat (LRR) defense gene family. High diversity in immune related responses is one possible pathogen resistance mechanism in plant communities^68^, and therefore reduced diversity may have compromised modern Arabica cultivar immunity.

The high level of conservation between the Arabica subgenomes and their diploid progenitors may have facilitated spontaneous interspecific hybridization events. This was the case for the Timor hybrid, a spontaneous Robusta x Arabica hybrid resistant to *Hemileia vastatrix*^27^. Our sample set included five descendants of the original Timor hybrid, obtained by backcrossing to Arabica. As expected, the hybridization affected subCC more profoundly, with much higher levels of nucleotide divergence apparent (*F*_*ST*_=0.185) than in subEE (*F*_*ST*_=0.0897), when comparing cultivars and hybrids. The divergence from wild populations was even greater, with *F*_*ST*_=0.254 for subCC and *F*_*ST*_=0.138 for subEE, illustrating that introgression occurred almost exclusively within subCC.

In the Timor hybrids, the regions found with d_f_ statistics^63^ largely overlapped the introgressed loci identified using *F*_*ST*_ scans (**Fig. 4A**) and were found in large blocks, reflecting recent hybridization, and covering 7-11% of the genome (**Fig. 4A, Ext. data Fig. 8**). Transposon Insertion Polymorphisms (TIPs) also overlapped with introgressed regions (Gypsy p=0.0002; Copia p=0.035; Fisher exact test), confirming their recent origin from Robusta (**Fig. 4B**). The introgressed regions overlapped with regions of higher subgenome fractionation (p=0.001873; **Table S13**), possibly due to heterologous recombination between subCC and Robusta, resulting in unequal crossing-over.

An introgressed region shared by all Timor hybrid lines was evident on chromosome 4 (**Fig. 4A**). We identified a set of 233 genes shared by all hybrids (**Table S14**). The set contained members of three colocalized tandemly duplicated blocks of resistance-related genes on chromosome 4, subCC, and showed high *F*_*ST*_ values between cultivars and introgressed lines. A tandem array of five genes were homologs of *Arabidopsis RPP8*, an NLR resistance locus conferring pleiotropic resistance to several pathogens^69,70^. *RPP8* shows a great amount of variation in *Arabidopsis* alone, where intrachromosomal gene conversion combined with balancing selection contributes to its exceptional diversity^71^. The same subCC region also included a tandem array of ten homologs of *CONSTITUTIVE EXPRESSER OF PR GENES 1* (*CPR1*), a negative regulator of defense response that targets resistance proteins^72,73^. Finally, we identified three duplicates encoding leaf rust 10 disease-resistance locus receptor-like protein kinases (LRK10L). The LRK10L are a gene family that is widespread across plants. First identified as a protein kinase in a locus contributing leaf rust resistance in wheat^74^, they found to be upregulated during various biotic and abiotic stresses^75^ and confirmed as positive regulators of wheat hypersensitive resistance response to stripe rust fungus^75^ and powdery mildew^76^.

The high *F*_*ST*_ values between cultivated and introgressed, but not wild individuals (**Fig. 4B**), indicate that the wild population cannot be the source for allelic asymmetries. Nucleotide diversities further illustrate this point; some genes demonstrate lower nucleotide diversity in wild individuals, suggesting these genes to have experienced selective sweeps. To further narrow down candidate genes involved in leaf rust resistance, we reanalyzed comparative gene expression data from susceptible and resistant accessions after *H. vastatrix* inoculation^77^. This analysis identified 723 differentially expressed genes, most of which were associated with defense responses (**Fig. 4B, Tables S14-S14b**). The combination of high *F*_*ST*_ values, nucleotide diversities, and differential expression data highlight several strong candidate genes (one *RPP8*, six *CPR1* and one *LRK10L*) at this locus.

## Conclusions and outlook

Besides providing genomic resources for molecular breeding of one of the most important agricultural commodities, our Arabica, Robusta and Eugenioides genomes provide a unique window into the genome evolution of a recently formed allopolyploid stemming from two closely related species. Our Arabica data did not suggest a genomic shock induced by allopolyploidy, but instead, only higher LTR transposon turnover rate. Genome fractionation rates remained basically unaltered before and after the allopolyploidy event. Likewise, no global subgenome dominance in gene expression was observed, but rather a mosaic-type pattern as in other recent polyploids^44,47^, affecting the expression of individual gene family members. However, similar to octoploid strawberry^8^, we detected genome dominance in terms of biased homoeologous exchanges favoring subCC. Since Robusta has one of the widest geographic ranges in the *Coffea* genus, whereas Eugenioides is more range-limited, this biased HE might be adaptive. This hypothesis was supported by the site frequency spectrum of HE loci, showing signs of directional selection (**Ext. data Fig. 3**). Intriguingly, transposable insertion polymorphisms significantly overlapped with tandem gene duplications and biosynthetic gene clusters, hinting at their possible roles in cluster evolution.

Domestication of perennial species like Arabica coffee differs markedly from that of annual crops, consisting instead of three phases: selection of outstanding genotypes from wild forests, clonal propagation and cultivation, then breeding and diversification^78^. In addition to being a perennial crop, Arabica is also a predominantly autogamous allopolyploid, which puts it in a class of its own. We show here that genetic diversity was already very low among wild accessions, due to multiple pre-domestication bottlenecks, and that the genotypes selected for cultivation by humans (both the ancient cultivated Ethiopian landraces, and the recent Geisha cultivar) already were somewhat admixed between divergent lineages. The resequenced accessions displayed a geographic split along the Eastern versus Western sides of the Great Rift Valley, with cultivated coffee variants all placed with the Eastern population. Such admixture has played a large role in breeding many fruit-bearing crops, the non-polyploid allogamous perennial lychee being one of the most extreme cases^59^.

The prevalent autogamy of Arabica, combined with the multiple genetic bottlenecks it underwent in the wild, may have selectively purged deleterious alleles, explaining the capacity of the species to survive single-plant bottlenecks that occurred during its cultivation. An additional element buffering deleterious alleles was probably Arabica’s allopolyploidy itself, which provided some level of heterosis^79^. However, the narrow genetic basis of both cultivated and wild modern Arabica constitutes a major drawback, as well as an obstacle for its breeding using wild genepool diversity. On the other hand, the extensive collinearity of its CC and EE subgenomes with those of its Robusta and Eugenioides progenitors is likely to facilitate introgression of interesting traits from these species, as already happened historically in the Timor spontaneous hybrid. The high-quality genome sequences of the three species provided in this work, together with the identification of the genomic region conferring resistance to coffee leaf rust, constitute a cornerstone for the breeding of novel Arabica varieties with superior adaptability and pathogen resistance.

## Data availability

Coffee genome assemblies are available at CoGe (https://genomevolution.org/): *C. canephora*: 50947, *C. eugenioides*: 60235, and *C. arabica*: 66663 (Pacbio HiFi) and 53628 (Pacbio). All genome information, including the VCF files with SNP information are available at ftp.solgenomics.net; the genome data is also available at ORCAE (https://bioinformatics.psb.ugent.be/orcae/overview/Coara and https://bioinformatics.psb.ugent.be/gdb/coffea_arabica/).

The sequencing data have been deposited to NCBI under bioproject ID: PRJNA698600.

## ACKNOWLEDGMENTS

The authors acknowledge the Natural History Museum in London for providing a sample of the *C. arabica* lectotype. JS acknowledges funding from Academy of Finland (decisions 318288 and 329441) and a Nanyang Technological University start-up grant. RG and S.O-A acknowledge funding from Ecos-Nord N°C21MA01 and STICAMSUD 21-STIC-13. PR acknowledges Academy of Finland (grant 343656). ARP acknowledges NAPI Bioinformática from Fundação Araucária and TELearning Project 2021-22 (21-STIC-13) from STIC AmSud. YB acknowledges the funding from Research Foundation - Flanders (FWO, No G056517N). YVdP acknowledges funding from the European Research Council (ERC) under the European Union’s Horizon 2020 research and innovation program (No. 833522) and from Ghent University (Methusalem funding, BOF.MET.2021.0005.01). GG acknowledges the Horizon Europe program, PRO-GRACE project (n. 101094738). AA acknowledges funding from INCT-Café-CNPq/Fapemig. DD and SMCL acknowledge São Paulo State Research Foundation (FAPESP), grant numbers #2016/10896-0 and #2017/01455-2. DS acknowledges funding from the NSERC and the Canada Research Chairs programs. VAA acknowledges United States National Science Foundation grants 1442190 and 2030871. PD acknowledges funding from Nestlé Research. JS wishes to acknowledge the High Performance Computation Centre at NTU Singapore and University of Helsinki Linux administrators, as well as the CSC – IT Center for Science, Finland, for computational resources.

## AUTHOR CONTRIBUTIONS

Conceived the study: AdK, DC, PD. Provided genetic resources: AA, AN, CK, EC, GHS, HR, LB, LFP, LP, MS, MTB, OGF, PaM, PM, PH, US. Carried out DNA sequencing: AC, CF, DM, GL, JeS, LB, MK, ND, PD, SM. Sequencing of the Linnaean accession: EG. Carried out genome assembly: SS, CW, JS, SP, LM. Genetic mapping: PR, MR, JS. Genome annotation: AR, SS, LM, JS, SR, VP, ZQW, DD, SIS, MM, RA, SMCL, ML, MP, CT-D, GG. Annotation of non-coding RNA: ARP, JE, PS. Transposable element annotation and analysis: SOA, AG, RG. Telomere identification: VAA, WCM. Analyzed genome evolution: ZY, ZC, DSM, RG, JM, DS, LC-P, TL, TK, VAA, SOA, AG, JS. Gene family analysis: ZQW, VP, DD, GG, SF, VAA, SiR, JS. RNA-seq data analysis: AR, SP, SiR, JS. Provided RNA-seq data: RH. Analyzed population data: JS. Analysed GBS data: YB, RG. Arranged online data access: LM, SR. Wrote the first draft: JS, completed with input from GG, DS, VAA, LFP, RG, SR, AdK, PD, VP, LM, DC, DD, SP, AA, as well as PM, YB, TR, YVdP, and all co-authors.

