## Supplementary material for "The genome and population genomics of allopolyploid *Coffea arabica* reveal the diversification history of modern coffee cultivars"

https://coffeabase.org/

Coffee BioProject ID PRJNA698600

### 1. Coffee history

Arabica cultivation was initiated in 15th -16th century Yemen, which held the world monopoly on coffee production at the time (**Fig. S1**). This domination was broken around 1600 AD by an Indian monk, Baba Budan, who smuggled the fabled "seven seeds" out of Yemen (Wellman, 1961), thus establishing Indian *Coffea arabica* cultivar lineages. In the 17th century, the Dutch obtained Arabica plants either from Sri Lanka or from India – these became the founding population of the contemporary Typica group – from which Arabica cultivation in Java and Southeast Asia was established. One plant from the same stock was shipped to Amsterdam in 1706, where one of its descendants was later donated to Louis XIV of France. Seeds from this cultivar were subsequently used to establish Arabica cultivation in the Caribbean, starting from Martinique in 1723. The French also began cultivating Arabica on the island of Bourbon (presently Réunion) from seeds obtained from Mocha, Yemen (Lécolier et al., 2009). Only one plant from this population survived by 1720, a single parent of the contemporary Bourbon group. Most important Arabica cultivars today are thought to be descendants of these Typica or Bourbon lineages, except for a few wild ecotypes whose origins can be traced back to natural forests in Ethiopia. Due to Arabica's recent allotetraploid origin and strong bottlenecks that occurred during its early cultivation and global spread, cultivated *C. arabica* harbors a particularly low genetic diversity with an effective population size ($N_e$) estimated to range between 10,000-50,000 individuals (Scalabrin et al., 2020).

To illustrate the geographic spread of coffee, we assembled relevant events of its history (Wellman, 1961) and displayed them on the world map (**Fig S1**).

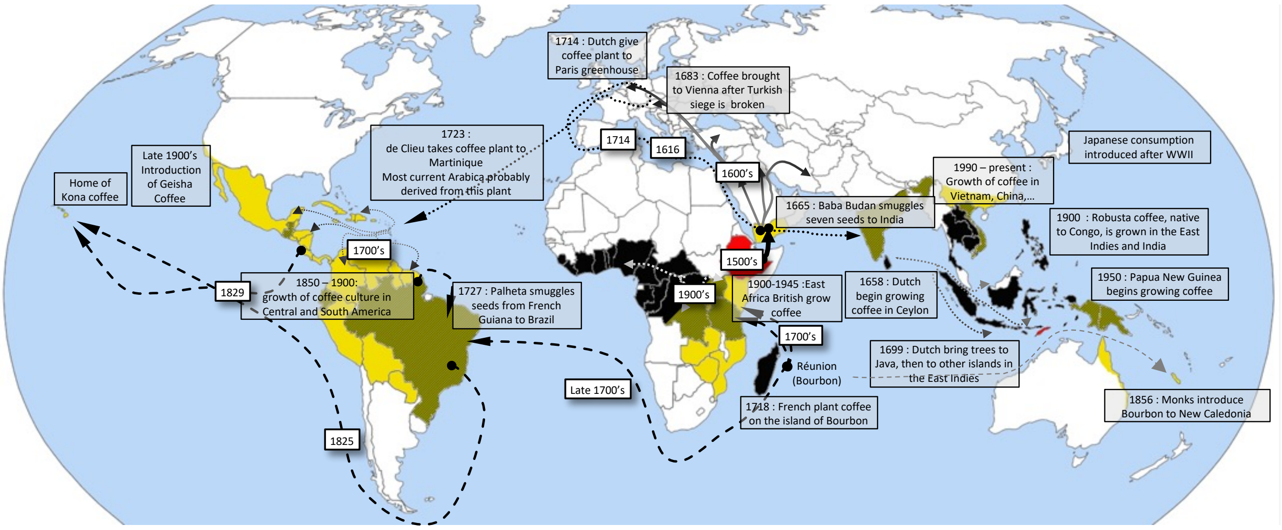


***Fig. S1.*** *Coffee dissemination routes. Yellow: current* Coffea arabica *cultivation; black: current* C. canephora *cultivation; ochre: current cultivation, both species. Solid lines: early spread of coffee consumption; dashed lines: main Bourbon routes; dotted lines: main Typica routes; Ethiopia (the center of origin of Arabica) and Timor Island (the origin of the Timor hybrid) are colored in red.*

### 2. Genome assembly

#### 2.1. Sequencing

For short read data, WGS genomic libraries for *Coffea arabica* were prepared following the manufacturer's standard instructions with an insert size of 400 base pairs. Paired-end sequences were generated on the Illumina HiSeq2000 platform by Weill Cornell Medical School.

For long read genomes of *C. canephora* and *C. eugenioides*, 20 kb genomic libraries were prepared following PacBio protocols including size selection with BluePippin. The same procedures were followed to obtain data for our first PacBio assembly of *C. arabica* (see below for details on our second assembly, which employed PacBio HiFi data). Sequencing was performed on a PacBio RS II instrument using P4/C2 and P5/C3 (*C. arabica*), and P6/C4 (*C. canephora* and *C. eugenioides*) chemistries, respectively, at Nestlé Research (**Table S16**). For HiFi reads at ~10 x coverage for *C. arabica* , SMRTbell libraries were prepared with the SMRTbell kit 3.0 and sequenced on a Sequel IIe using adaptive loading, 2 h of pre extension and 30 h movies, at Nestlé Research.

For population resequencing, the libraries of 39 wild and cultivated *C. arabica* accessions were prepared using the KAPA HyperPrep Kits (Roche) following manufacturer's instructions, and paired-end sequencing (2 x 125) to ~ 40 x coverage was carried out on Illumina HiSeq2500 platforms at Nestlé Research.

#### 2.2 Assembly & Dovetail scaffolding

A contig-level assembly for *Coffea canephora* was obtained using MHAP (Berlin et al., 2015). Scaffolding was performed using BAC-end sequences and 454 paired-end sequences generated previously (Denoeud et al., 2014). Both *C. eugenioides* and *C. arabica* were assembled with Falcon (Chin et al., 2016), and *C. arabica* was subsequently phased using Falcon_unzip. Assemblies were corrected with six rounds of PILON (Walker et al., 2014) before submitting for HiC scaffolding by Dovetail. Gap-filling for the HiC assemblies was carried out with PBJelly (English et al., 2012) followed by polishing with 6 rounds of correction using PILON with DNA-seq/RNA-Seq data.

Dovetail HiC libraries were prepared in a similar manner as described previously (Lieberman-Aiden et al., 2009). Briefly, for each library, chromatin was fixed in place with formaldehyde in the nucleus and then extracted. Fixed chromatin was digested with *Dpn*II, the 5' overhangs filled in with biotinylated nucleotides, and then free blunt ends were ligated. After ligation, crosslinks were reversed, and the DNA purified from protein. Purified DNA was treated to remove biotin that was not internal to ligated fragments. The DNA was then sheared to ~350 bp mean fragment size and sequencing libraries were generated using NEBNext Ultra enzymes and Illumina-compatible adapters. Biotin-containing fragments were isolated using streptavidin beads before PCR enrichment of each library. For *C. arabica*, the libraries were sequenced on an Illumina HiSeqX to produce 222 million 2x151 bp paired-end reads, which provided 50x physical coverage of the genome (1-50kb pairs). For *C. canephora*, the libraries were also sequenced on an Illumina HiSeqX platform, and produced 273 million 2x151 bp paired-end reads, which provided 100x physical coverage of the genome (1-50kb pairs).

The input *de novo* assembly, shotgun reads, and Dovetail HiC library reads were used as input data for HiRise, a software pipeline designed specifically for using proximity ligation data to scaffold genome assemblies (Putnam et al., 2016). Shotgun and Dovetail HiC library sequences were aligned to the draft input assembly using a modified SNAP read mapper (http://snap.cs.berkeley.edu). The separations of Dovetail HiC read pairs mapped within draft scaffolds were analyzed by HiRise to produce a likelihood model for genomic distance between read pairs, and the model was used to identify and break putative misjoins, to score prospective joins, and make joins above a threshold (**Fig. S2**). After scaffolding, shotgun sequences were used to close gaps between contigs.

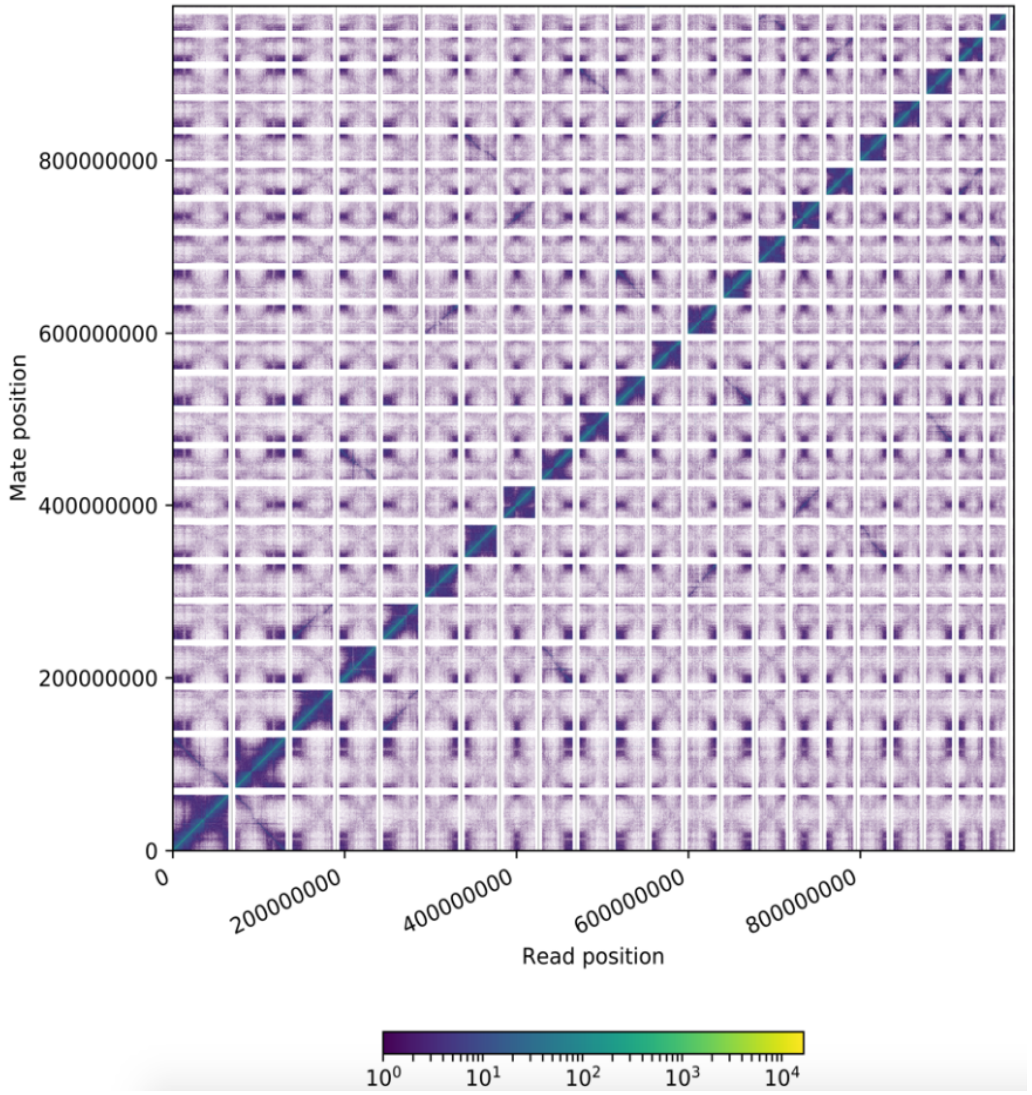


***Fig. S2.*** *A HiC contact heatmap depicting the linkages between different scaffolds. The x and y axes give the mapping positions of the first and second read in the read pair respectively, grouped into bins. The color of each square indicates the number of read pairs within that bin, as described in the color key.*

##### 2.3 PacBio HiFi assembly

Arabica Pacific Biosciences HiFi long reads were assembled into contigs using Hifiasm (Cheng et al., 2021). Next, Dovetail HiC reads were used for scaffolding using a combination of the ALLHiC (Zhang et al., 2019) and 3d-dna (Dudchenko et al., 2017) pipelines. HiC reads were first mapped to the contig-level assembly using BWA-mem (Li & Durbin, 2009) and cleaned to remove redundant, chimeric, and uninformative read pairs. To assist in the scaffolding, the contig assembly was initially annotated with GeMoMa (Keilwagen et al., 2019), and then the predicted coding sequences were aligned using mmseq (Steinegger & Söding, 2017) against the chromosome-level genome assembly of the diploid parental species *Coffea canephora*, in order to anchor contigs to homologous chromosomes based on synteny. The resulting allelic table was used as a reference to partition contigs, and it cleanly mapped reads into 12 homologous chromosome groups using ALLHiC_partition. Following the ALLHiC pipeline, we then pruned the noisy HiC signal, and split the contigs into three groups within each homologous chromosome set (assuming two groups to correspond to two homologous *C. arabica* chromosomes and one group for possible interchromosomal rearrangements).

Following the split of the gene-containing contigs, we then used ALLHiC_rescue to add contigs without genes to the scaffold-level assembly, and then optimized the ordering and orientation of the contigs using ALLHiC optimize, and built scaffold files for each homologous chromosome set using ALLHiC. Chromosome assemblies were then visualized and manually corrected in juicebox (Robinson et al., 2018) and finalized by a post-review run in 3d-dna. Mapped contigs from each chromosome assembly were integrated and used in the global rescue step in ALLHiC, which was then optimized to obtain a whole-genome assembly. This assembly was visualized in Juicebox and corrected manually, and 3d-dna was run from the post-review step to obtain a highly accurate final genome assembly (**Fig. S3**). Next, gap-closing was carried out first by using TGS-gapCloser (M. Xu et al., 2020) and HiFi reads longer than 15k, followed by gap-filling with Racon (Vaser et al., 2017). The contigs that were not assigned to pseudomolecules were found to contain many organellar fragments. Therefore, all reads were aligned against the gap-closed pseudomolecules and organellar genomes. The reads that did not align were then collected and a new assembly was made. Finally, the assembled contigs with 100% match to pseudomolecule assemblies were removed. The completeness of the genome assembly was assessed using BUSCO v. 5.2.2 (Manni et al., 2021).

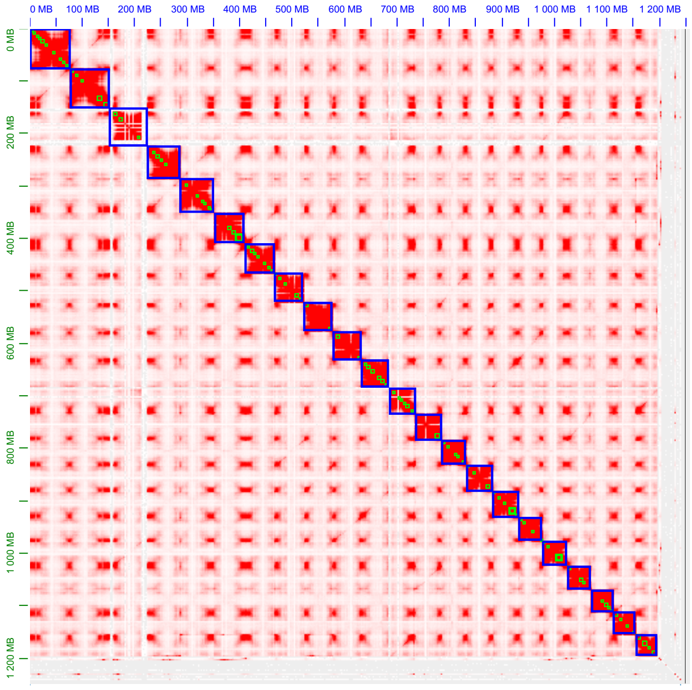


***Fig. S3.*** *Hi-C contact map of the* C. arabica *PacBio HiFi assembly. The x and y axes give the mapping positions of the first and second read in the read pair respectively, grouped into bins. The color intensity of each square is proportional to the number of read pairs within that bin.*

#### 2.4 Linkage map

A consensus international reference genetic map of *C. canephora* has been developed by Nestlé Research, Tours and ICCRI based on a F1 cross between two highly heterozygous genotypes, a Congolese group genotype (BP409) and a Congolese x Guinean hybrid parent (Q121). The segregating population was composed of 93 F1 individuals (Lefebvre-Pautigny et al., 2010).

In order to estimate an ultra-high density linkage map, the parents were sequenced to 60x and progeny to 20x coverage using the Illumina HiSeq2000 platform at Nestlé Research. Following quality control with FastQC and trimming with Trimmomatic v0.36 (Bolger et al., 2014), the reads were mapped against the *C. canephora* reference assembly using BWA-mem v0.7.15 (Li & Durbin, 2009).

The linkage mapping was conducted with (LM3) Lep-MAP3 (Rastas, 2017) as follows. First, the input genotype likelihoods (called posteriors in LM3) were obtained from individual bam files using SAMtools pileup (Li et al., 2009), pileupParser2.awk and pileup2posterior.awk of LM3. Then the ParentCall2 (LM3) module was called on the likelihoods and the pedigree file to obtain final input data. Thereafter, the markers were clustered into paternal (informativeMask=1) and maternal (informativeMask=2) linkage groups by using a LOD score of 18 (parameter lodLimit=18) in a segregation distortion aware model (distortedLod=1) within SeparateChromosomes2. The groups found that were matched to chromosomes corresponded to 11 major scaffolds in the reference genome. Finally, the markers in each chromosome were ordered using module OrderMarker2 with default parameters. The total number of markers was about 720k and 610k, for paternal and maternal markers, respectively. The markers were used to determine the orientation and order along the pseudomolecules. The final assembly combining the two parental maps, solving conflicts as well as identification of haplotype alleles, was carried out manually (**Fig. S4**).

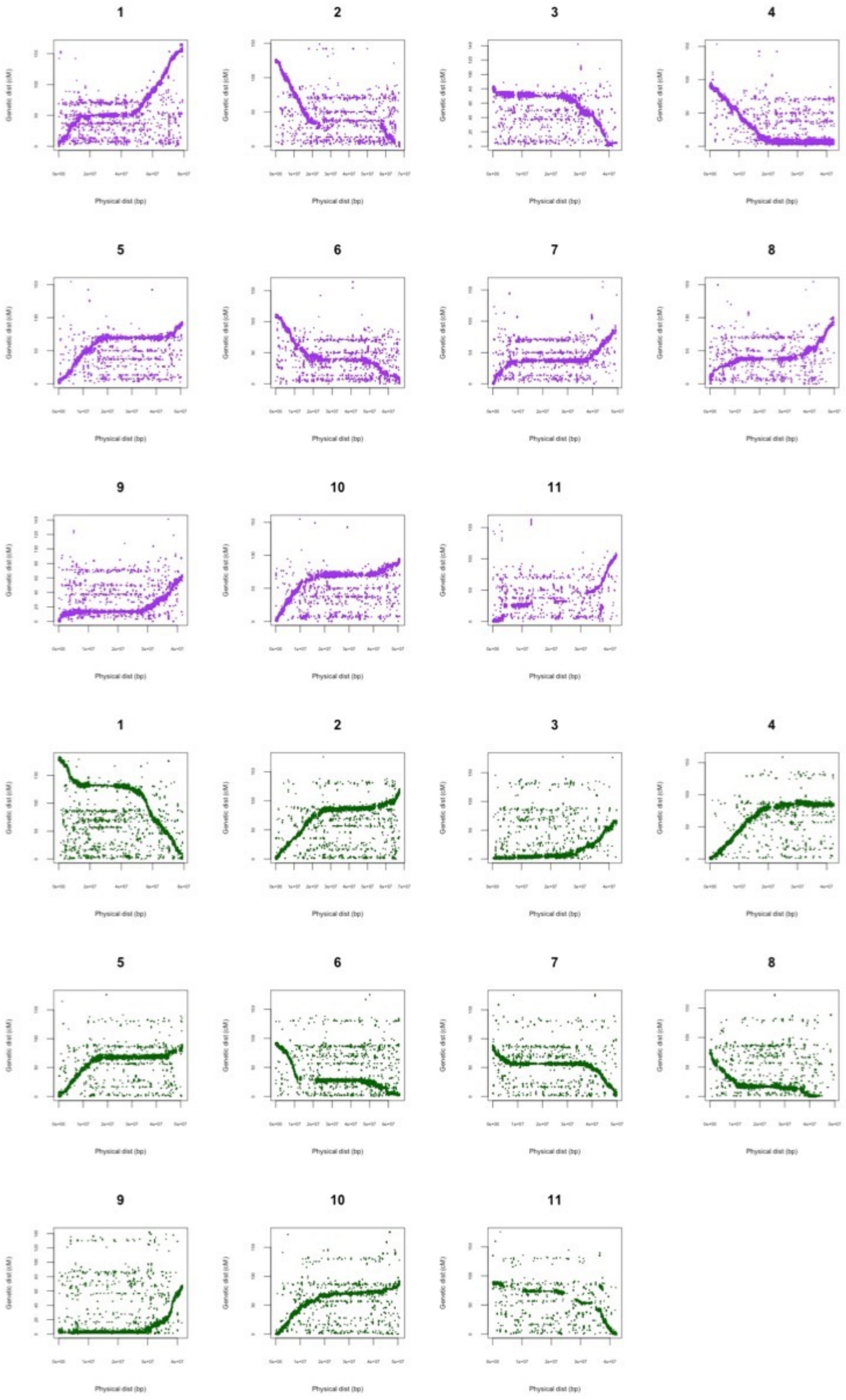


***Fig. S4.*** *Marey maps of* C. arabica. *Top: Maternal map, Bottom: Paternal map. X-axis shows the physical coordinate of the markers in the assembly, y-axis the genetic position.*

##### 2.5 Assembly metrics and quality control

The assemblies were compared to previously published assemblies of *Coffea canephora* (Denoeud et al., 2014) and *C. arabica* (Scalabrin et al., 2020) (**Table 1**).

### 3 Annotation

##### 3.1 RNA-Seq and IsoSeq

RNA-Seq libraries (**Table S17**) were prepared from leaf RNA using the TruSeq Stranded mRNA Kit (Illumina) following the manufacturer's instructions, and then paired-end sequenced at a coverage of 40-80 million reads per sample on the Illumina HiSeq2500 platform.

For Isoform Sequencing (IsoSeq) (**Table S18**), one RNA extraction from young leaves of the di-haploid Arabica accession was used for full-length transcript sequencing. The library was prepared starting from 1 µg of RNA following the PacBio protocol IsoSeq using the Clontech SMARTer PCR cDNA Synthesis Kit and the Bluepippin Size Selection system (PN 100-377-100-02). The library was divided into four fractions based on size selection, and each fraction was sequenced on the PacBio RSII platform at Nestlé Research. For every fraction, 18 SMRT cells were run using P6/C4 chemistry.

##### 3.2 Retrotransposons and other repetitive elements

**Construction and analysis of *de novo* transposable element database**

EDTA (https://doi.org/10.1186/s13059-019-1905-y) was used to *de novo* identify transposable elements (TEs) in the *Coffea canephora* and *C. eugenioides* genomes and *C. arabica* subgenomes. Inpactor2 (https://doi.org/10.1093/bib/bbac51) was used to recover full-length LTR retrotransposons in the three genomes and to classify them at the lineage level (https://github.com/simonorozcoarias/Inpactor2). EDTA and Inpactor2 libraries were merged and clustered using cd-hit (doi: 10.1093/bioinformatics/bts565). Clusters were manually inspected to remove nested and false predictions. After curation, libraries were used for annotation using Repeat Masker (default parameters). Annotations with length > 200bp were retained.

**Timing of LTR retrotransposon insertions**

The timing of LTR retrotransposon insertions was studied in the three genomes using individual sequences recovered by Inpactor2 and using an average base substitution rate of 1.3E–8 (Ma & Bennetzen Jeffrey, 2004), similarly to (Orozco-Arias et al., 2018) (**Fig. S5**). All three genomes showed the same trend of recent insertions of elements between 0-1.5 mya (**Fig. S5**).

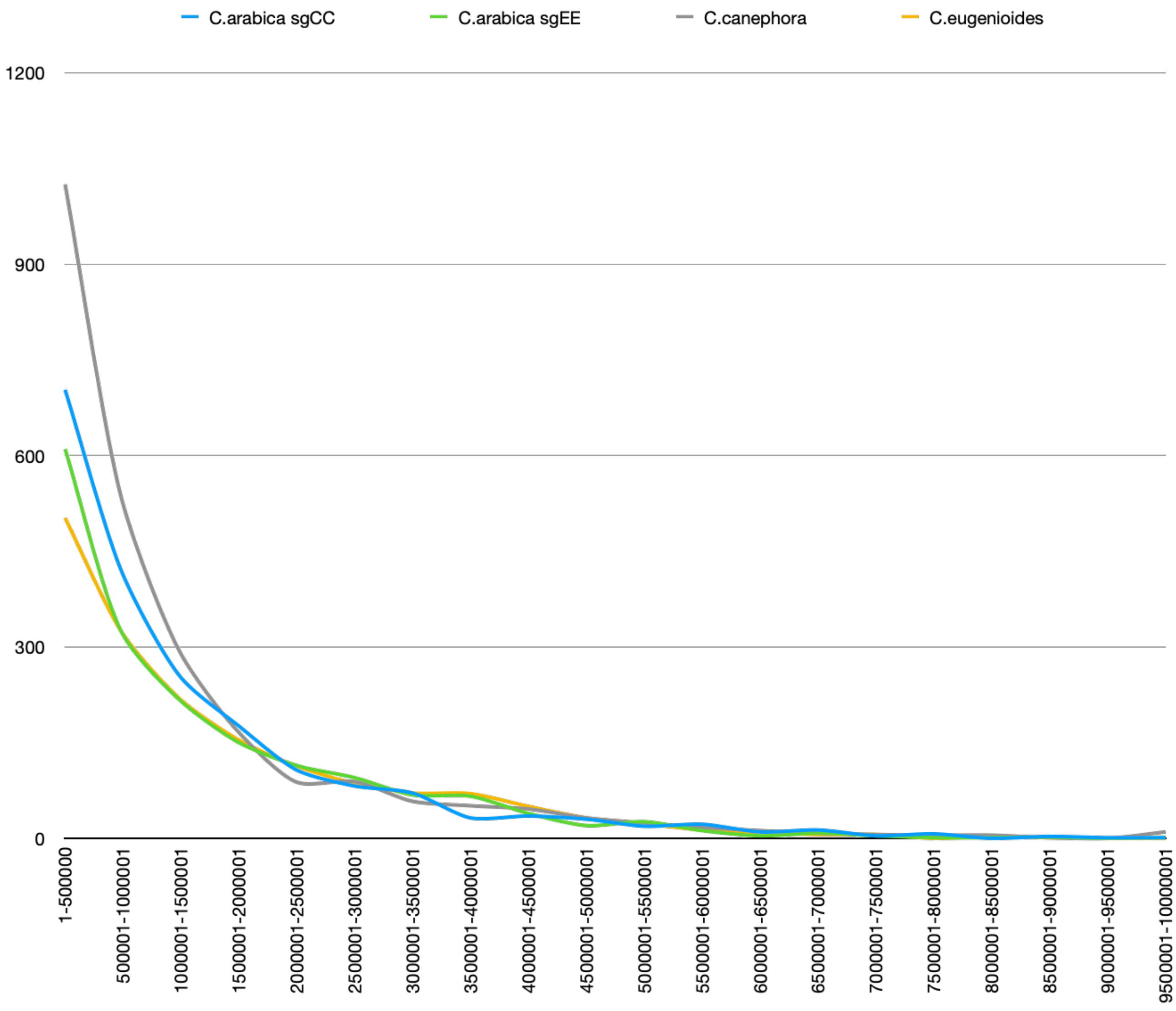


***Fig. S5.*** *Timing of LTR Retrotransposon insertions in the* Coffea arabica *subgenomes (subCC, blue; subEE, green),* C. canephora *(grey) and* C. eugenioides *(orange). Full-length LTR retrotransposons are identified by Inpactor2 per bins of 0.5 My. An average base substitution rate of 1.3E–8 was used.*

At the level of superfamilies (RLC: Copia; RLG: Gypsy), variations were observed for Copia elements (RLC). A small peak of insertions was observed at 1.5-2 My for *C. arabica* and at 2-2.5 My for *C. eugenioides*. No evident peak of insertion was observed for *C. canephora* (**Fig. S6**). At the level of lineages, clear variations were observed between all three genomes (**Fig. S7-S8**).

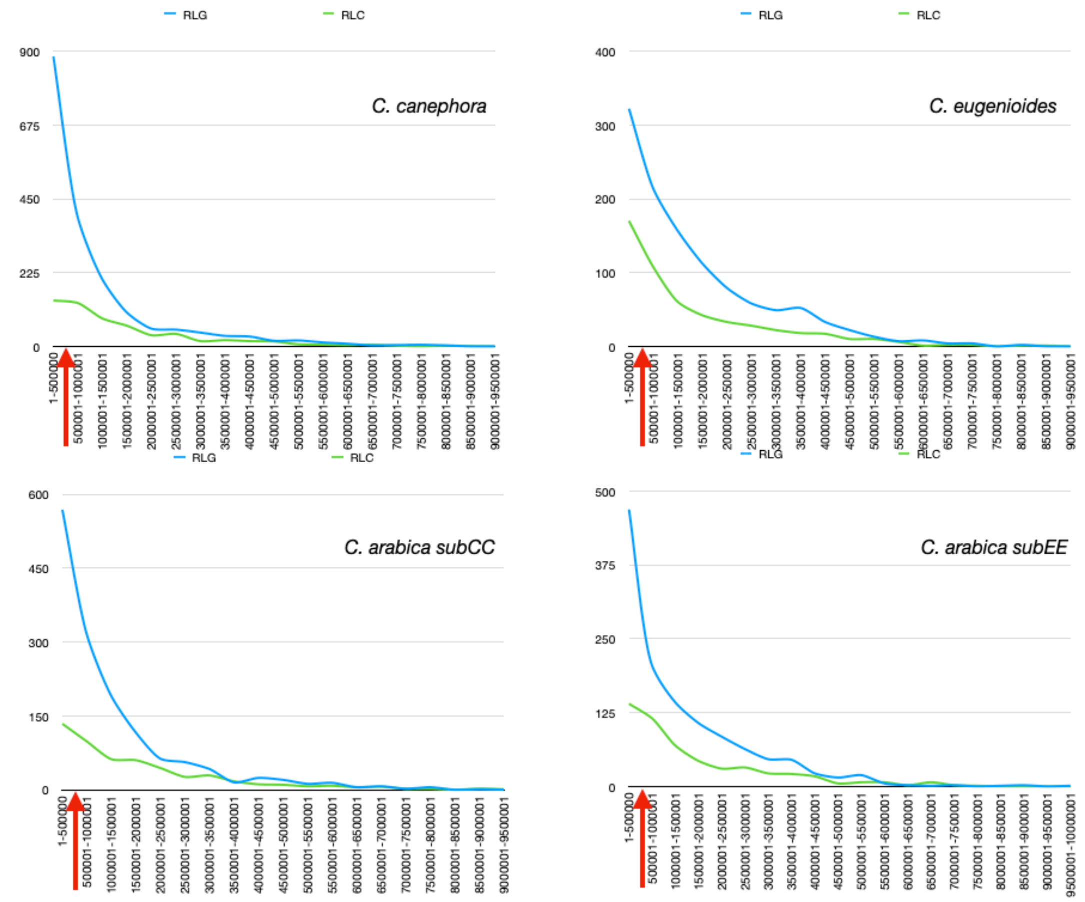


***Fig. S6.*** *Timing of RLG (Gypsy) and RLC (Copia) LTR retrotransposon insertions in* Coffea canephora, C. eugenioides, *and the subgenomes of* C. arabica. *Full-length LTR retrotransposons are identified by LTR_STRUC per bins of 0.5 My. An average base substitution rate of 1.3E–8 was used.*

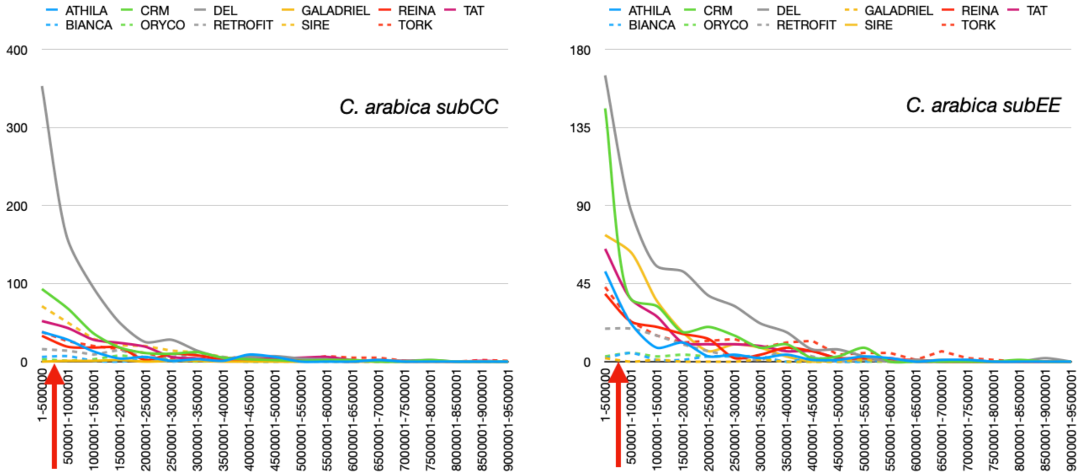


***Fig. S7.*** *Timing of LTR retrotransposon lineage insertions in* Coffea arabica *subgenomes. Full-length LTR retrotransposons were identified by LTR_STRUC per bins of 0.5 My. An average base substitution rate of 1.3E–8 was used.*

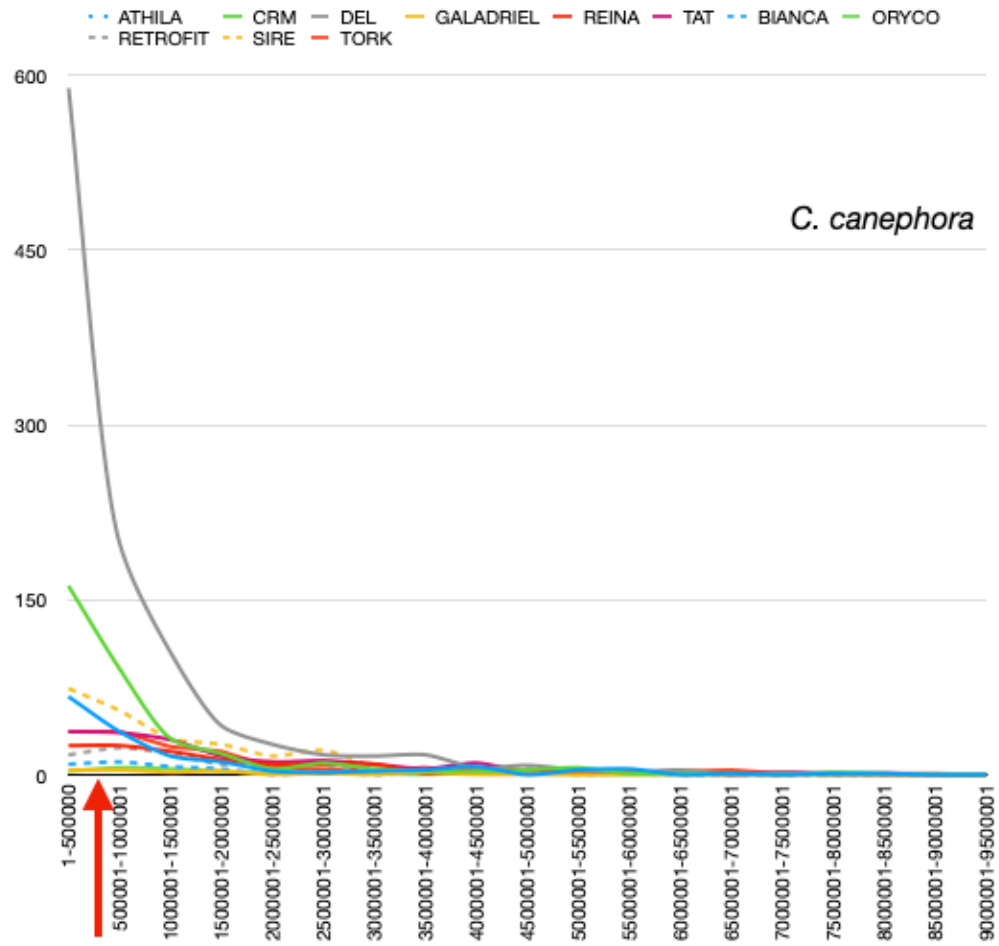


***Fig. S8.*** *Timing of LTR retrotransposon lineage insertions in* Coffea canephora*. Full-length LTR retrotransposons were identified by LTR_STRUC per bins of 0.5 My. An average base substitution rate of 1.3E–8 was used.*

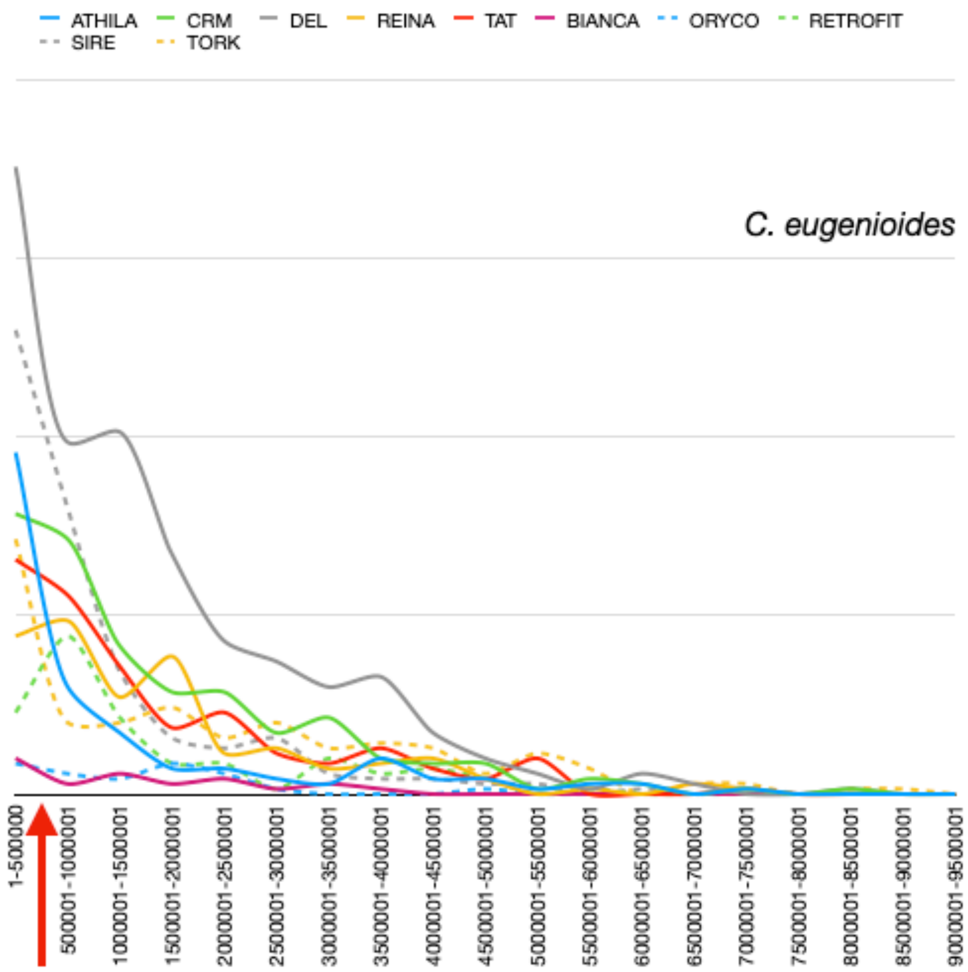


***Fig. S9.*** *Timing of LTR retrotransposon lineage insertions in* Coffea eugenioides*. Full-length LTR retrotransposons were identified by LTR_STRUC per bins of 0.5 My. An average base substitution rate of 1.3E–8 was used.*

#### Masking and annotating *Coffea* genomes

The EDTA/Inpactor2 libraries, merged into one file, were used for masking (RepeatMasker default parameters) the genomic sequences in *Coffea canephora, C. eugenioides* and *C. arabica.*

The merged library masked 67,47%, 59,72%, 63,14% and 63,82% for *C. canephora, C. eugenioides,* and *C. arabica* subCC and subEE, respectively. At the TE order and superfamily levels, no significant differences were observed between the three genomes (**Fig. S10**) (*C. canephora* vs subCC p=0.06805, *C. eugenioides* vs subEE p=0.6802; *C. canephora* vs *C. eugenioides* p=0.511; subCC vs subEE p=0.8797; pairwise t-test). However, individual TE families may still show marked differences, such as LTR Gypsy retrotransposons.

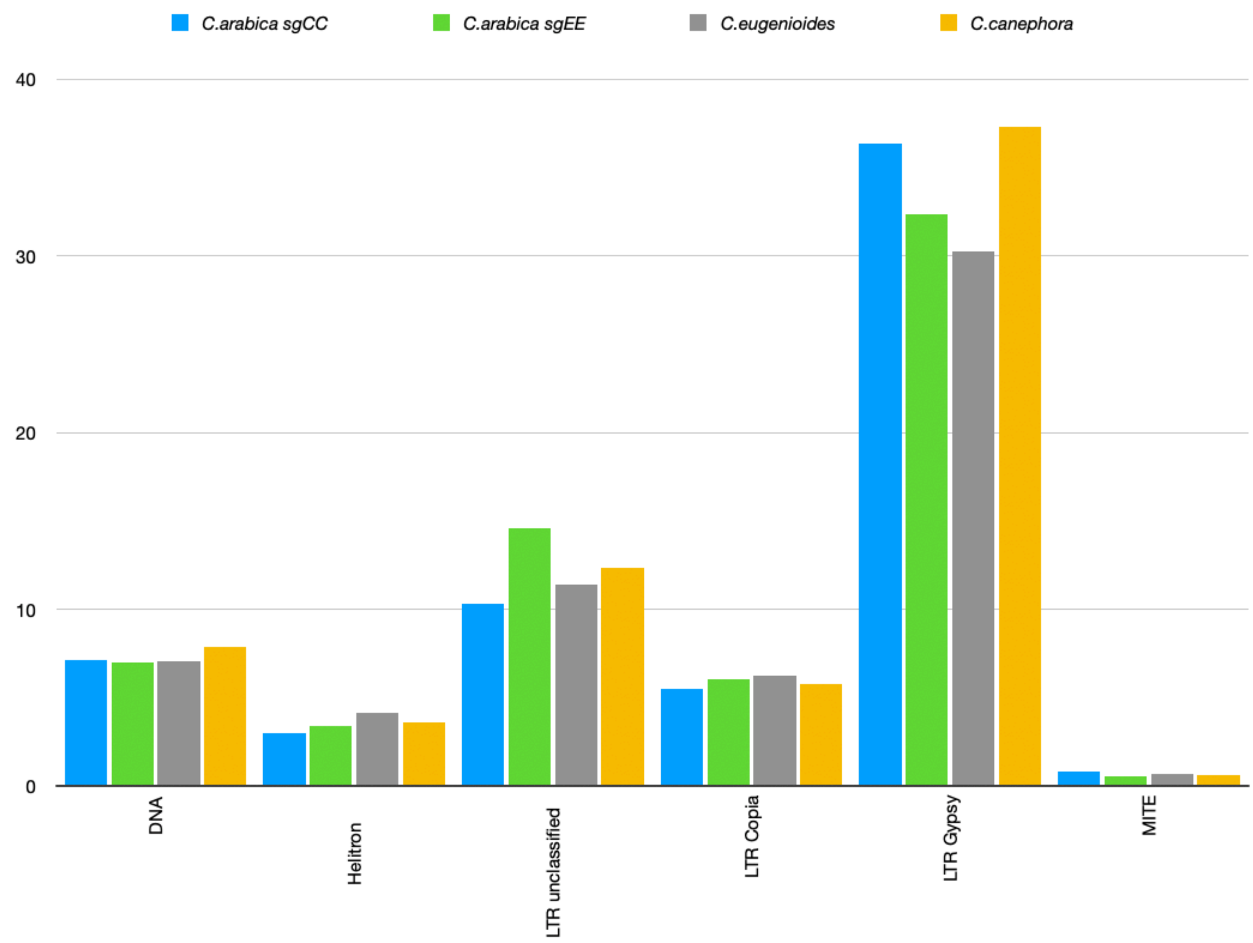


***Fig. S10.*** *Genome proportion (%) of transposable element categories in the* Coffea canephora*,* C. eugenioides *and* C. arabica *subgenomes.*

Similarly, no statistically significant differences in Gypsy or Copia LTR retrotransposon lineages were found. However, there were marked differences in the variation of genome proportion (%) for DEL, CRM, TAT and SIRE (**Fig. S11**).

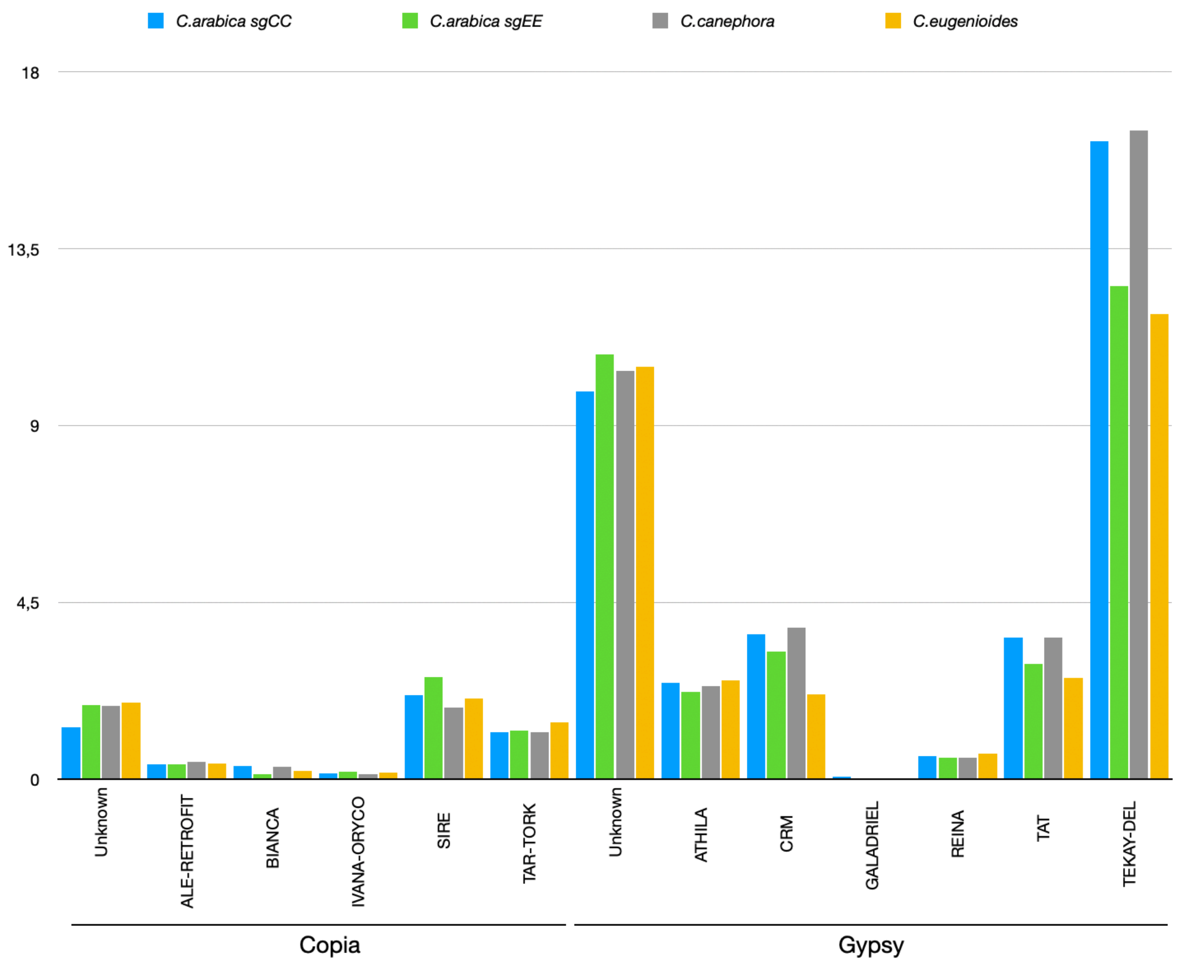


***Fig. S11.*** *Genome proportion (%) of LTR retrotransposon lineages in* Coffea canephora*,* C. eugenioides *and* C. arabica *subgenomes.*

**Distributions of transposable elements along *Coffea arabica, C. canephora* and *C. eugenioides* pseudomolecules**

The distributions of transposable elements along *Coffea arabica, C. canephora* and *C. eugenioides* pseudomolecules were studied with a window size of 200 kb using DensityMap (Guizard et al., 2016). A negative association between coding genes and TE distribution was observed for all pseudomolecules (**Fig. S12**). LTR retrotransposons were mainly located in pericentromeric regions, while DNA transposons were mainly positioned in distal regions. For LTR retrotransposons, Copia elements were equally distributed, while Gypsy elements were mainly pericentromeric. However, Copia SIREs were found mainly in pericentromeric regions, while the Gypsy ATHILA elements were found in distal regions. As previously published, CRMs formed a clear peak in pericentromeric regions, likely located in centromeric regions themselves. All of these observations were similar along the *C. arabica, C. canephora* and *C. eugenioides* pseudomolecules.

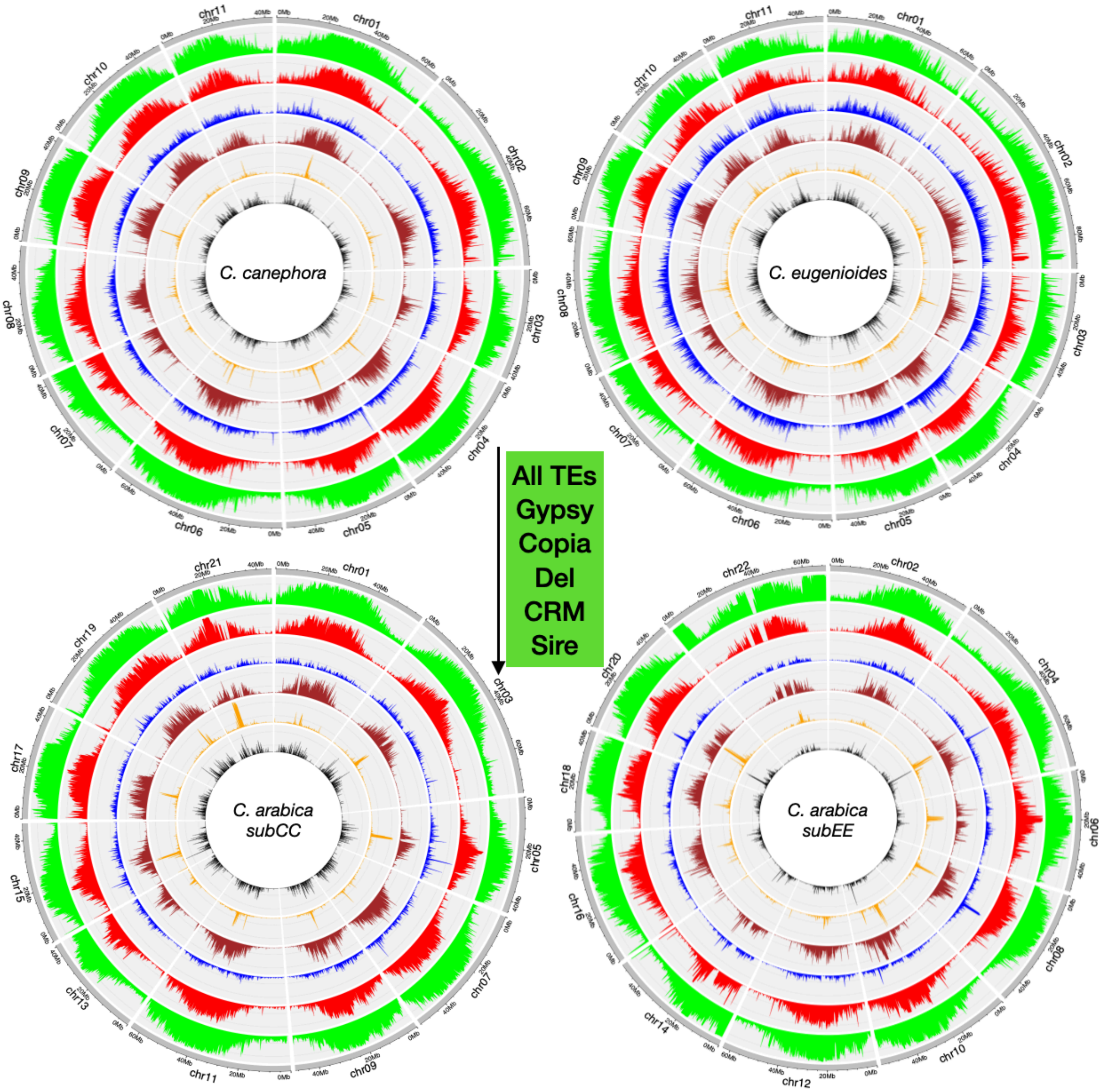


***Fig. S12.*** *Circos representations of the distribution of genes and TEs along pseudomolecules of* Coffea canephora*,* C. eugenioides *and subgenomes of* C. arabica*, using windows of 200 kb.*

**Phylogenetic analysis of Reverse Transcriptase (RT) domains.**

The phylogenetic analysis of RT domains was performed as in Yu et al. (doi: 10.1038/ng.3435). Briefly, RT domains (> 150 residues) were identified from the genomes using GeneWise (https://www.ebi.ac.uk/%7Ebirney/wise2/). Recovered sequences were aligned using Mafft (https://mafft.cbrc.jp/alignment/software/), and FastTree was used to infer approximately-maximum-likelihood phylogenetic trees. Trees were edited with itol (https://itol.embl.de). 5474 and 4824 domains were recovered for *C. canephora* and *C. arabica* subCC (**Fig. S13**), and 4616 and 4820 domains were recovered for *C. eugenioides* and *C. arabica* subEE (**Fig. S14**).

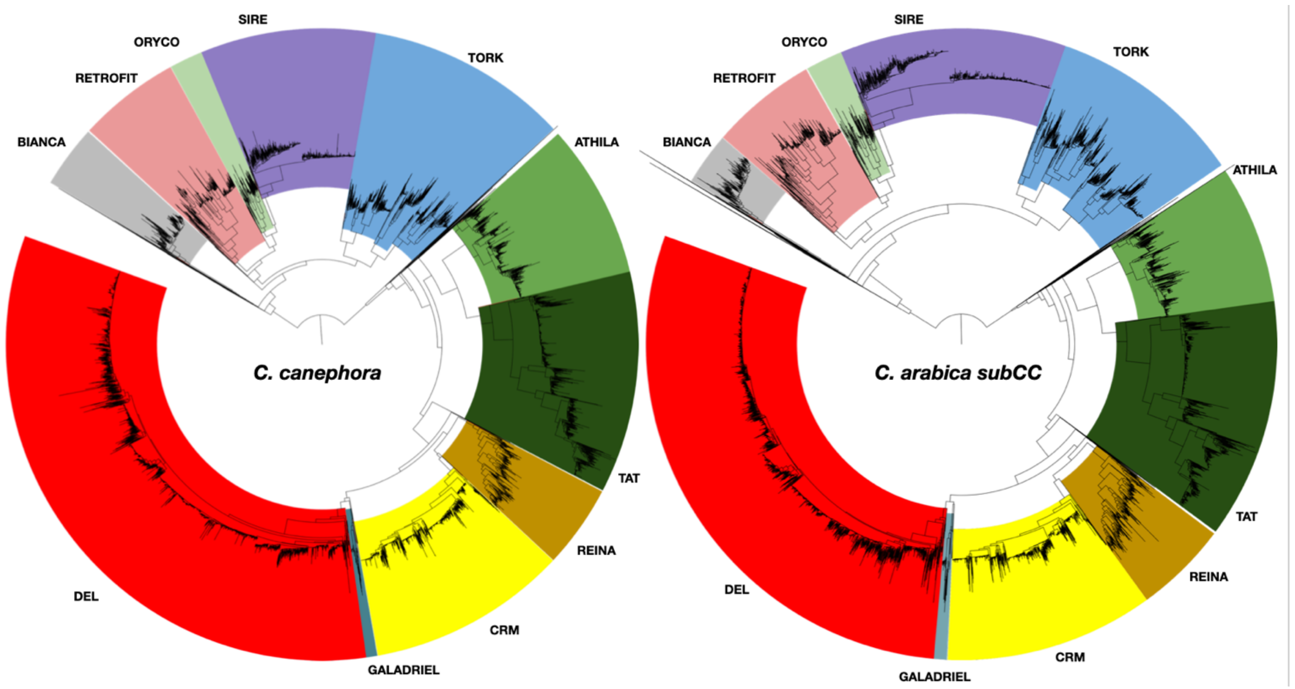


***Fig. S13.*** *Phylogenetic analysis of RT domains from* Coffea canephora *and* C. arabica *subCC reconstructed on the basis of 5474 and 4824 aligned sequences.*

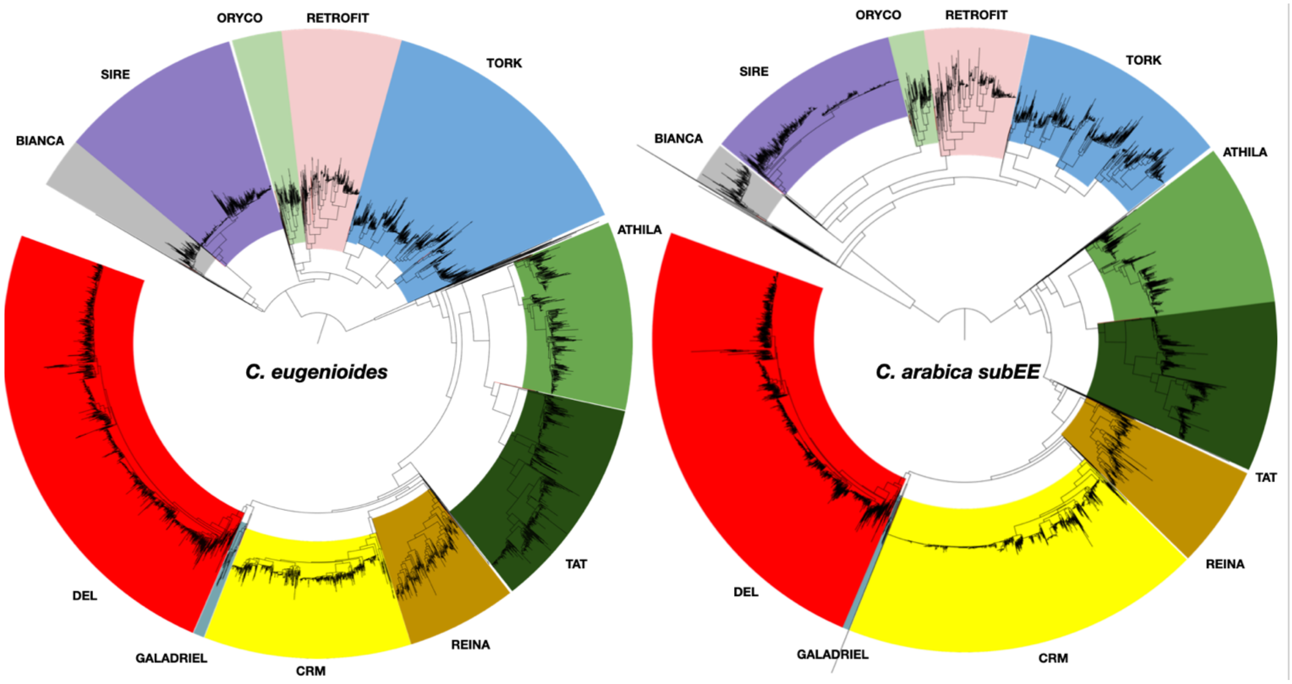


***Fig. S14.*** *Phylogenetic analysis of RT domains from* Coffea eugenioides *and* C. arabica *subEE reconstructed on the basis of 4616 and 4820 aligned sequences.*

#### 3.3 Telomeric repeats

*Arabidopsis* type telomeric repeat sequences $(TTTAGGG)_n$ were searched for (as the following polymer query: TTTAGGGTTTAGGGTTTAGGGTTTAGGGTTTAGGGTTTAGGG) in the *C. arabica* HiFi and *C. canephora* v1.8 genomes using BLASTN with e-value cutoff of 1E-5, word size 8, existence:5 Extension:2, Match/Mismatch 1,-2, and HSP maximum 35,000. The search was carried out using CoGeBlast (https://genomevolution.org/coge/CoGeBlast.pl).

To assess the completeness of the genome assembly, we analyzed the occurrence and position of telomeric repeats found in the three *Coffea* genomes. Altogether 33480 hits to the telomeric repeat query were annotated in the *C. arabica* genome, including 33 located at the end of chromosomes, one on unanchored scaffolds, and many interstitial clusters within chromosomes (**Fig. S15**). Fourteen *C. arabica* chromosomes contained telomeric repeats at both ends, indicating complete assembly of those chromosomes, while 5 chromosomes contained an array on one end only.

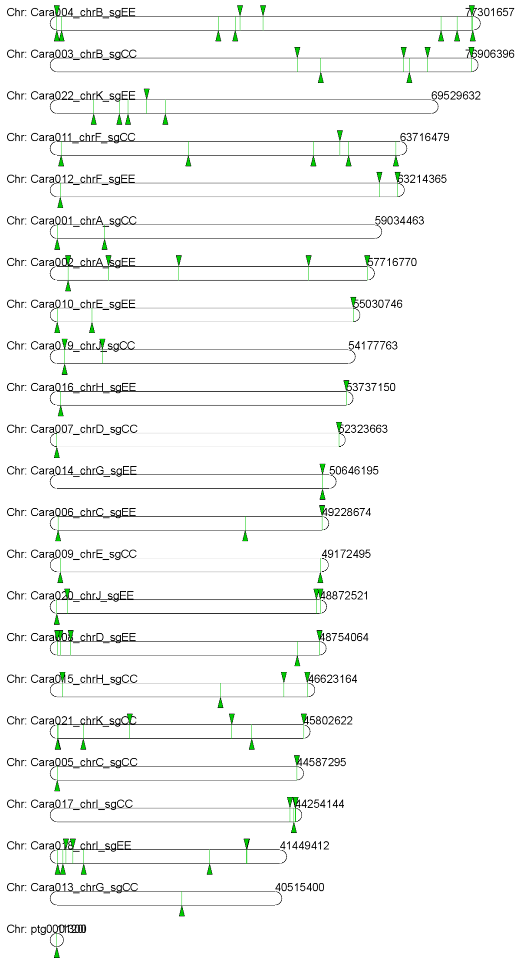


***Fig S15.*** *Telomere arrays in the* Coffea arabica *HiFi assembly. The green arrows show the locations in chromosome idiograms.*

For the *C. canephora* genome, telomeric arrays were found at both chromosomal ends for 7 chromosomes, and one chromosomal end for 4 chromosomes (**Fig. S16**). Many interstitial telomeric repeat sequences were found across chromosomes. The level of end-chromosome array completeness suggests that the chromosomal-level assembly of *C. canephora* is high quality. However, only one telomeric array was found in the *C. eugenioides* genome. This is likely due to the filtering of PacBio contigs with high repetitive sequences during the genome assembly procedure, with the result that telomeric repeats were not found in the final assembly.

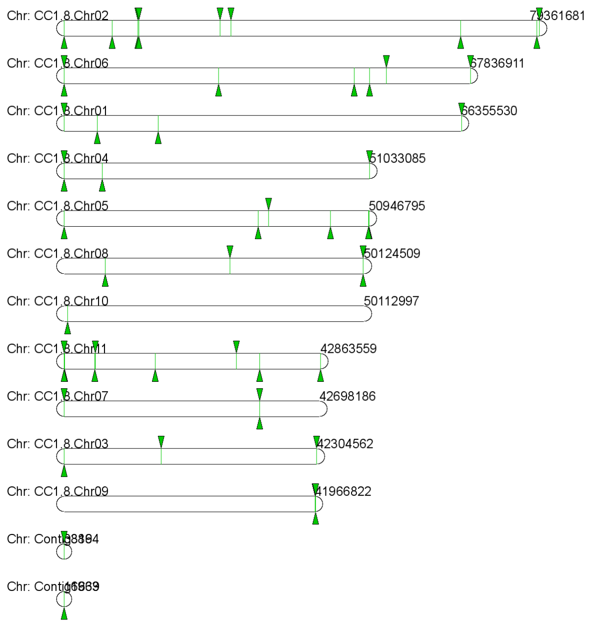


***Fig S16.*** *Telomere arrays in the* Coffea canephora *assembly. The green arrows show the locations in chromosome idiograms*

#### 3.4 Structural and functional gene annotations

##### 3.4.1 Annotation of protein coding genes

A MAKER-based annotation strategy was adopted and refined for gene annotation. IsoSeq and RNA-Seq data was mapped to the repeat-masked genome using default parameters in HiSAT (Kim et al., 2015) (**Fig. S17**). The resulting alignment files were read by the intron-exon junction predicting software Portcullis (Mapleson et al., 2018). The predicted exon-intron junctions along with alignment files in GTF format were used with Mikado software (Venturini et al., 2018) to predict gene models. Gene model predictions from Mikado, Augustus (Stanke et al., 2006), Genmark (Borodovsky & McIninch, 1993) and SNAP (Korf, 2004) were integrated into one final gene annotation using MAKER (Cantarel et al., 2008). In MAKER, each gene model was assigned an Annotation Edit Distance (AED) score based on the level of supporting evidence, where genes without any support have score of 1, and 0 denotes gene3 models with perfect agreement with the supporting evidence (Campbell et al., 2014). High evidence gene models with AED score <0.5 were selected for the annotation. Quality and completeness of the annotation was assessed by BUSCO scores (Simão et al., 2015) (**Table 1**).

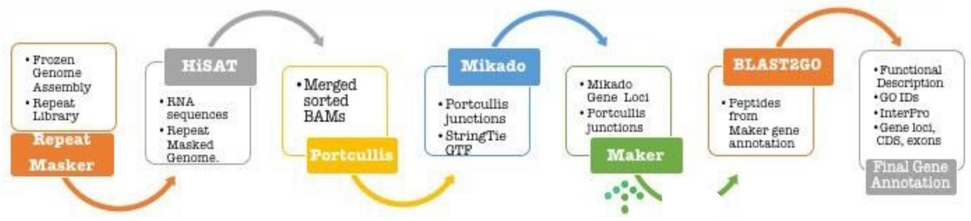


***Fig. S17.*** *Genome annotation pipeline.*

In the case of the *Coffea arabica* HiFi assembly, the high quality annotations for *C. canephora*, *C. eugenioides* and the earlier *C. arabica* assembly (v 0.99) were transferred using GeMoMa (Keilwagen et al., 2019) and then combined using the same toolbox.

##### 3.4.2 Manual gene curation

Altogether 210 genes from key metabolic pathways impacting coffee quality were manually curated (**Table S19)** and fully annotated using WebApollo software (Lee et al., 2013), including enzymes involved in caffeine, sucrose and chlorogenic acids (phenylpropanoid pathway) biosynthesis, among others, for the *Coffea canephora* (70 genes) and *C. eugenioides* (70 genes) and *C. arabica* (140 genes) genomes. The annotated sequences were detected by sequence homology using BLAST with previously published gene sequences as queries for the caffeine pathway (Denoeud et al., 2014; McCarthy & McCarthy, 2007; Perrois et al., 2015), sucrose pathway (Privat et al., 2008), and phenylpropanoid pathway including chlorogenic acids (Lepelley et al., 2007; Lepelley et al., 2012).

Importantly, this in-depth annotation confirmed the high quality of the genome sequences, the annotation using WebApollo as the annotation platform, and was consistent with gene sequences already published for key metabolic pathways in *C. arabica* and in *C. canephora*.

Overall, the availability of *C. arabica*, *C. canephora* and *C. eugenioides* genome sequences and their respective annotations will be particularly useful for assisting coffee breeding programs and association studies. The identification of genetic markers such as SNPs encoding proteins associated with key traits in coffee (coffee cup quality, technical and agronomical traits) (Merot-L'anthoene et al., 2019) will be crucial for the selection of coffee varieties with improved traits, e.g., cultivars with higher cup quality or higher reliance to diseases and drought stress.

#### 3.5 Mapping between *C. arabica* HiFi versus previous PacBio assembly

Gene model mappings between the two assembly versions (**Table S20**) was obtained by carrying out a syntenic alignment using the SynMap2 tool (Haug-Baltzell et al., 2017) available in the CoGe platform (genomevolution.org).

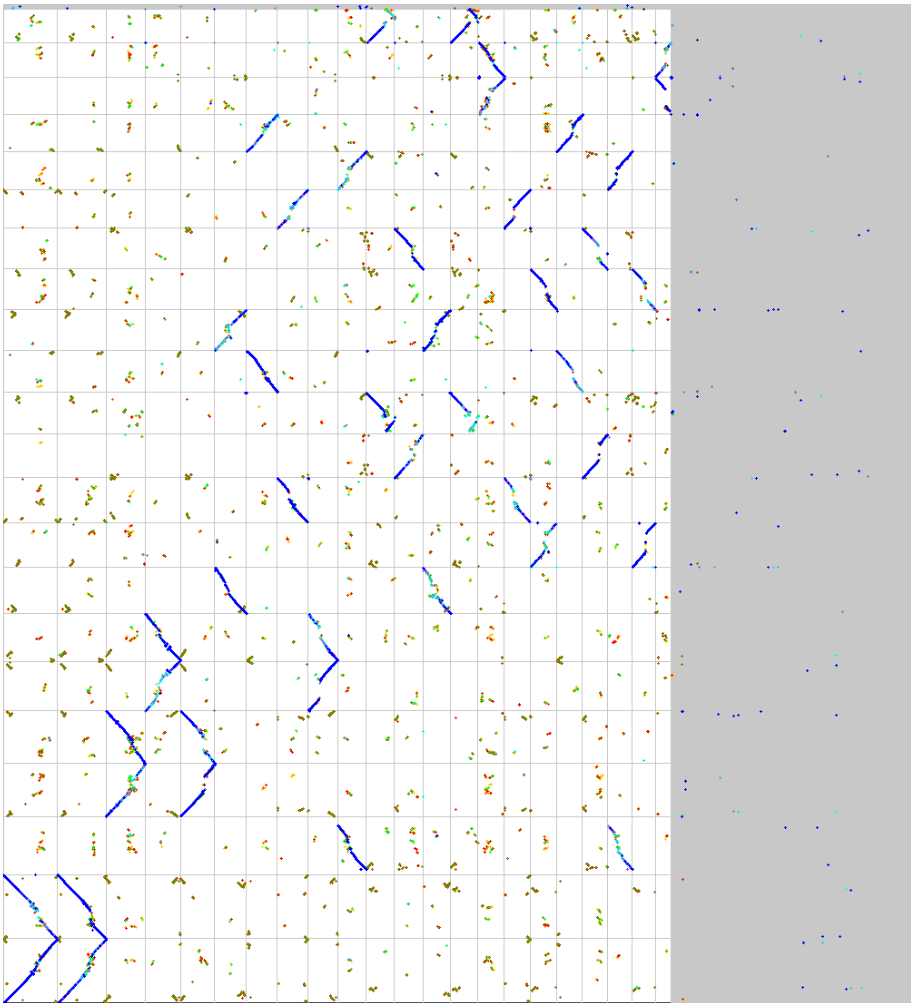

***Fig S18.*** *Syntenic alignment of the* Coffea arabica *HiFi assembly (y-axis) against* C. arabica *PacBio + Dovetail v0.9 (x-axis). The points reflect syntenic gene blocks between the assemblies, and the color palette corresponds to synonymous substitutional gene similarity (with blue reflecting no or very little difference, lighter colors suggesting greater distance).*

Overall, syntenic comparison of the two assemblies (**Fig. S18**) suggested that the gene space is assembled relatively similarly in the two assemblies. The largest differences were in the pericentromeric and telomeric repetitive regions, where the new PacBio HiFi assembly has more additional sequence. Note, for example, the second from bottom chromosome on the y-axis against the first and second chromosomes on the x-axis.

##### 3.6 Subgenome phasing using per-chromosome $K_s$ distributions

In order to test the split of the *Coffea arabica* assembly into two subgenomes, $K_s$ estimates between *C. arabica* and each of two diploid outgroups, *C. canephora* and *C. eugenioides*, were compared. First, $K_s$ values were extracted from CoGe SynMap (https://genomevolution.org/coge/SynMap.pl) output and $\log_{10}$ transformed. In each comparison, $\log_{10}$ $K_s$ values less than -3 and equal to or greater than 0 were removed, as these fell outside of the main peak corresponding to divergence between *C. arabica* and each of the diploid outgroups (**Fig. S19**).

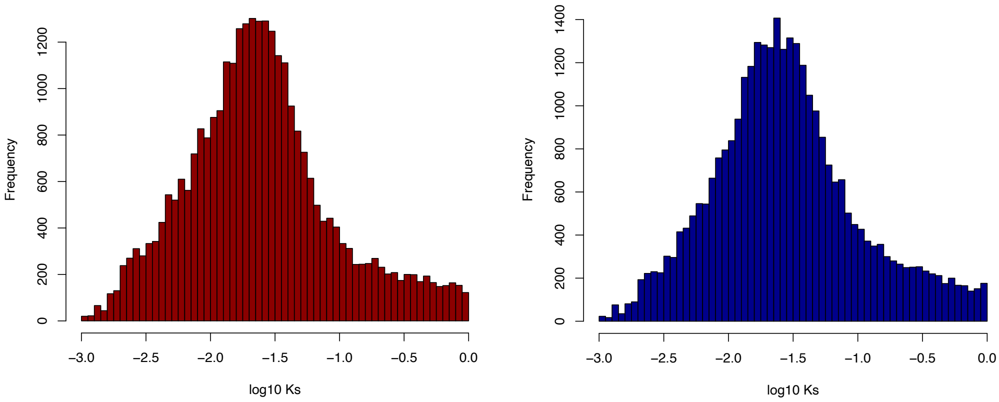


***Fig. S19.*** *Histograms depicting $log_{10}$ $K_s$ distributions following trimming for* Coffea arabica *versus* C. canephora *(left panel), and* C. eugenioides *(right panel).*

Next, mean Ks values per chromosome were estimated between *C. arabica* and each of the diploid outgroups, respectively. Pairs of homoeologous chromosomes were compared for each comparison to assess whether patterns of Ks match subgenome phasing from other datasets (**Fig. S20**).

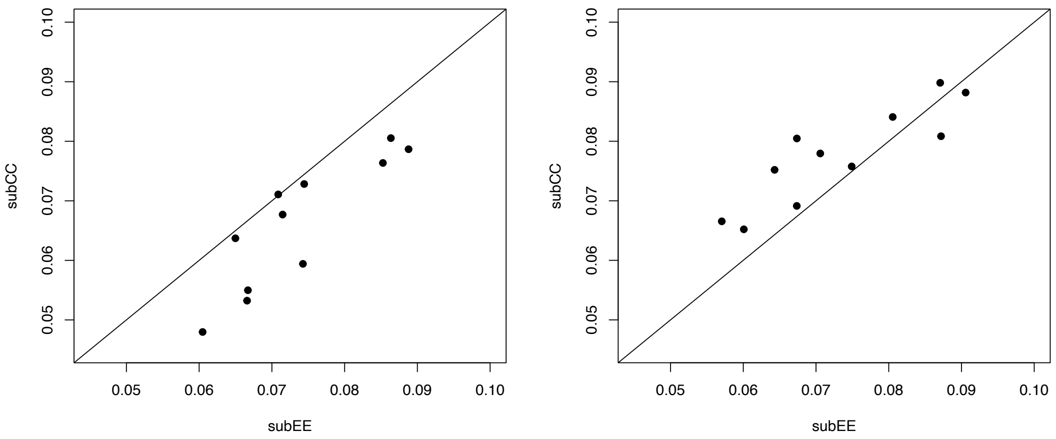


***Fig. S20.*** *Scatterplots of* Coffea arabica *homoeologous pairs (mean Ks per chromosome) versus* C. canephora *(left panel) and* C. eugenioides *(right panel) outgroups. For the comparison with* C. canephora *(left panel), the eugenioides subgenome chromosomes generally (10/11 pairs) have higher Ks values than the respective canephora homoeolog, suggesting that* C. canephora *is more similar to subCC than to subEE. For the comparison with* C. eugenioides *(right panel), the opposite is generally true (9/11 pairs).*

Finally, we compared $K_s$ values for chromosomes from the putative subCC subgenome versus *C. canephora* to $K_s$ values for chromosomes from the putative subEE subgenome to *C. eugenioides* (**Fig. S21**). Nine of the eleven homoeolog pairs showed greater similarity of the subCC subgenome to the *C. canephora* outgroup versus subEE subgenome to the *C. eugenioides* outgroup. Phasing using this $K_s$ approach is generally consistent with the subgenome designations outlined previously.

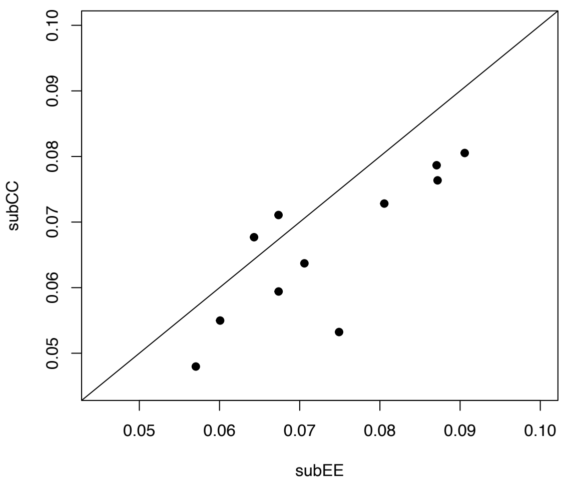


***Fig. S21.*** *Scatterplots of* Coffea arabica *homoeolog pairs depicting the mean chromosome-wide Ks values for the* C. eugenioides *outgroup versus putative subEE subgenome chromosomes (x-axis) versus Ks values for their respective homoeologous pairs between the* C. canephora *outgroup and the putative subCC subgenome chromosomes (y-axis).*

### 4 Genome Evolution

#### 4.1 Fractionation in *Coffea*

In a SynMap comparison of the two Arabica subgenomes, there were 7461 subCC genes in synteny blocks that did not have a subEE counterpart in that synteny block, and 7691 subEE genes in synteny blocks that did not have a subCC counterpart in that synteny block. Part of this was due to the minimum block size (5) criterion of SynMap. If this size was reduced to 2 or 1, the number of unpaired subCC genes dropped by 1190 or 2120, and the number of unpaired subEE genes dropped by 1240 or 2179.

Unexpectedly, most of these genes missing from a subgenome were not lost after tetraploidization. The great majority were already absent from expected syntenic positions in the ancestors of subgenomes. Only 835 of the missing subEE genes were found paired in a subCC-CE syntenic block, and only 950 of the missing subCC genes were found paired in a subEE-CC syntenic block. Only these are likely to have undergone fractionation after tetraploidization. Thus, 89% of the missing subEE genes were already missing before tetraploidization, as were 88% of the missing subCC genes.

Polyploidization is only one of the processes giving rise to paralogous pairs of genes. Tandem duplicates and other mechanisms endow genomes with many duplicate pairs, some of which may be almost contemporaneous with the genome-wide event under study. Thus, the same gene may be present in two or more synteny blocks, since it may be homologous to two tandem-originated copies of a single gene. After fractionation of one of these copies, the gene will be matched in one of the synteny blocks and not the other. Thus, aside from the 7461 unmatched subCC genes and 7691 unmatched subEE genes, another 1697 subCC genes and 1580 subEE genes, respectively, were unmatched in one synteny block with the other subgenome but matched in another synteny block (and counted as matched). Most of these, 1567 in subCC and 1479 in subEE, were matched in synteny blocks with Eugenioides and Robusta, respectively, almost all to only one gene. The remaining 130 and 101 genes, unmatched in any synteny block with a parent genome, may have jumped to their current synteny block from some other place in the genome, but this is speculative, lacking other evidence.

More importantly, 1233 of the 1567 subCC genes and 1242 of the 1479 subEE genes were matched in exactly one of two synteny blocks with Eugenioides and Robusta respectively, so that their partially fractionated status in the tetraploid was inherited from the parent. The remaining 325 and 237 genes, matched in only one synteny block with a parent genome, may also be candidates for jump translocation status, but this also remains speculation.

In summary, most of the apparent fractionation in *Coffea arabica* reflects the difference in gene complement between the parent genomes *C. canephora* and *C. eugenioides* prior to the tetraploidization event.

A substantial proportion of duplicate gene loss from synteny blocks, whether orthologous or paralogous, left alternative homological relations in other synteny blocks. The surviving gene originally had multiple homologs.

#### 4.2 Subgenome Fractionation

Gene loss from the subgenomes did not occur randomly over the length of the chromosomes. **Fig. S22** and **Fig. S23** illustrate how gene loss was more pronounced in the pericentromeric regions of all chromosomes.

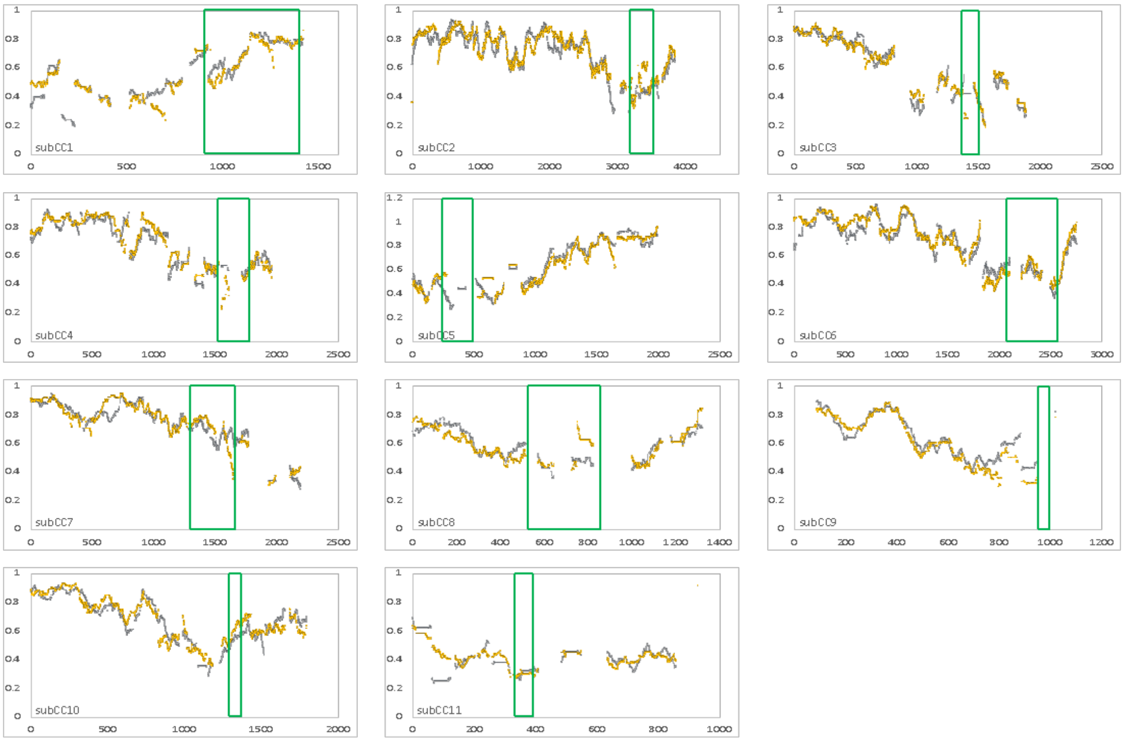


***Fig. S22.*** *For each pair of homoeologous chromosomes in* Coffea arabica*, the proportion of genes retained in synteny blocks, including singletons and paired genes over total number of original pairs, measured in a window of size +/- 50 genes for each gene, is plotted. Graphs are shown in coordinates of subCC gene order. Rectangle indicates centromeric region. Orange: subEE genes. Grey: subCC genes.*

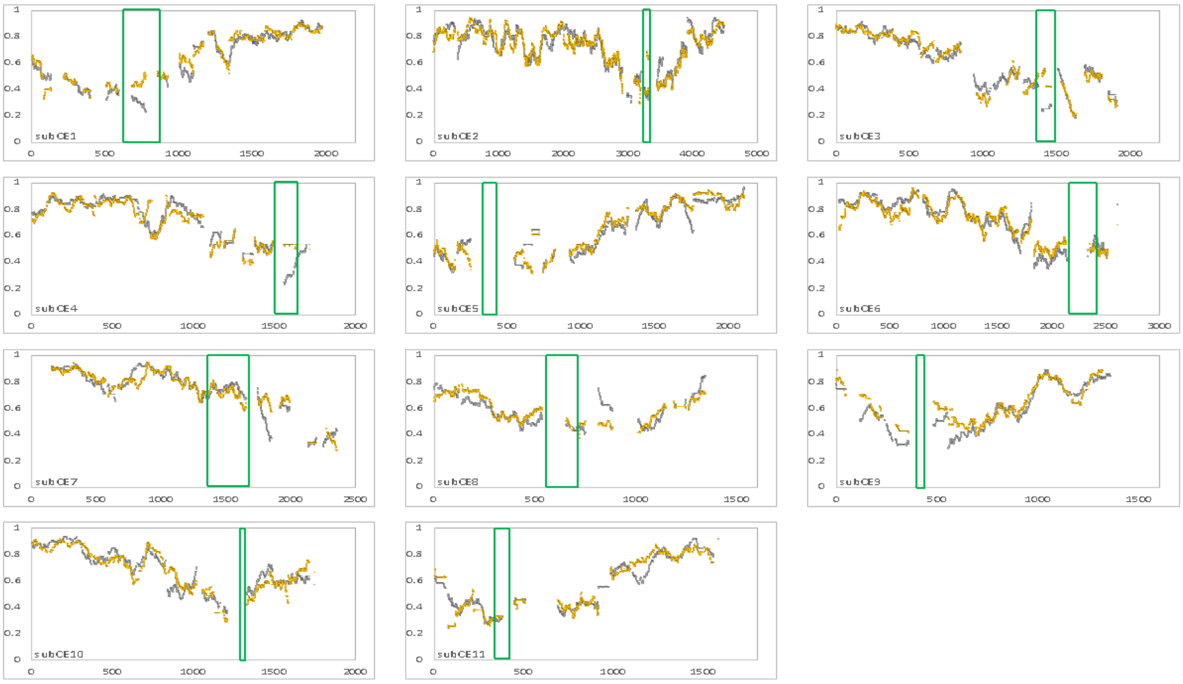


***Fig. S23.*** *For each pair of homoeologous chromosomes in* Coffea arabica*, the proportion of genes retained in synteny blocks, including singletons and paired genes over total number of original pairs, measured in a window of size +/- 50 genes for each gene, is plotted. In coordinates of subEE gene order. Rectangle indicates centromeric region. Orange: subEE genes. Grey: subCC genes.*

The parallelism between the fractionation patterns of the two subgenomes is striking, but even more so when we consider that these patterns were largely inherited from their progenitor genomes. **Fig. S24** and **Fig. S25** depict the parallel rates of depletion of synteny blocks in *Coffea canephora* and *C. eugenioides* in a SynMap comparison of the two genomes. The conclusion is that the pattern must reflect a structural tendency towards gene loss originating before the speciation of the two progenitors.

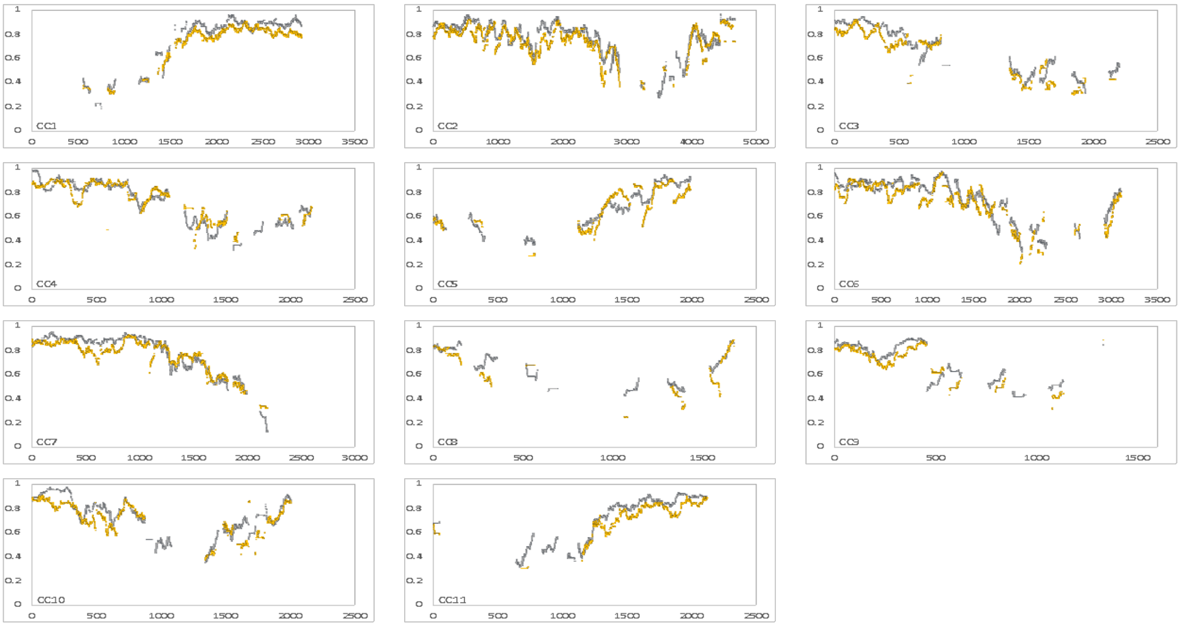


***Fig. S24.*** *For each pair of orthologous chromosomes in* Coffea canephora *and* C. eugenioides*, the proportion of genes retained in synteny blocks, including singletons and paired orthologs over total number of original pairs, measured in a window of size +/- 50 for each gene, is plotted. In coordinates of CC gene order. Orange: EE genes. Grey: CC genes.*

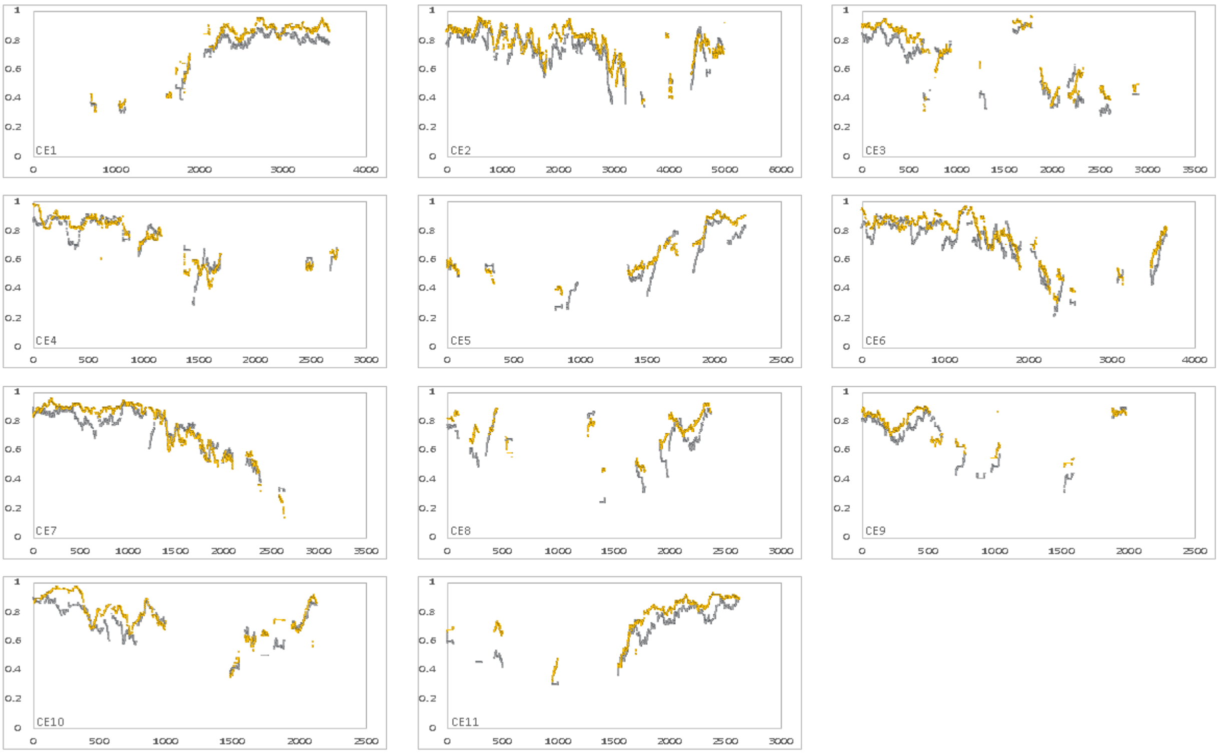


***Fig. S25.*** *For each pair of orthologous chromosomes in* Coffea canephora *and* C. eugenioides*, the proportion of genes retained in synteny blocks, including singletons and paired genes over total number of original pairs, measured in a window of size +/- 50 for each gen,e is plotted. In coordinates of EE gene order. Orange: EE genes. Grey: CC genes.*

#### 4.3 Excision versus Pseudogenization

To assess whether excision or pseudogenization accounts for the *C. arabica* gene loss data, we note that pseudogenization, leaving the gene intact, would not drastically shorten the length of the chromosomal region it is in. In contrast, excision of genes, including some or all of the flanking intergenic DNA, will shorten the region. **Fig. S26** shows that the length of the intergenic region remaining after the loss of one or more genes is relatively constant at about 10,000 bp, whereas the length of the corresponding region in the other genome, including the genes that were not lost and their intergenic regions, increases by about 20,000 bp per gene in subCC and 15,000 bp in subEE. So, the regions that have lost annotated genes have lost almost all their DNA sequence. This is striking evidence in favor of the predominance of excision events. Note that our analysis was confined to the quantitatively most important case where either the neighbors in subCC are adjacent or the neighbors in subEE are adjacent, excluding the cases where there are missing genes between neighbors on both subgenomes.

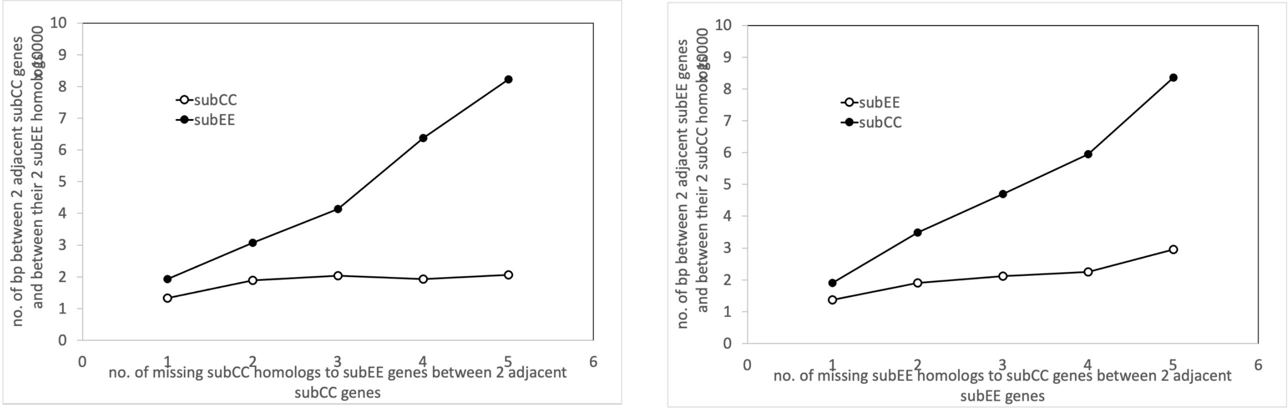


***Fig. S26.*** *Lengths of DNA segments between genes made adjacent by fractionation of intervening genes still present in the opposite subgenome. Left: fractionation in subCC. Right: in subEE*

**a)**

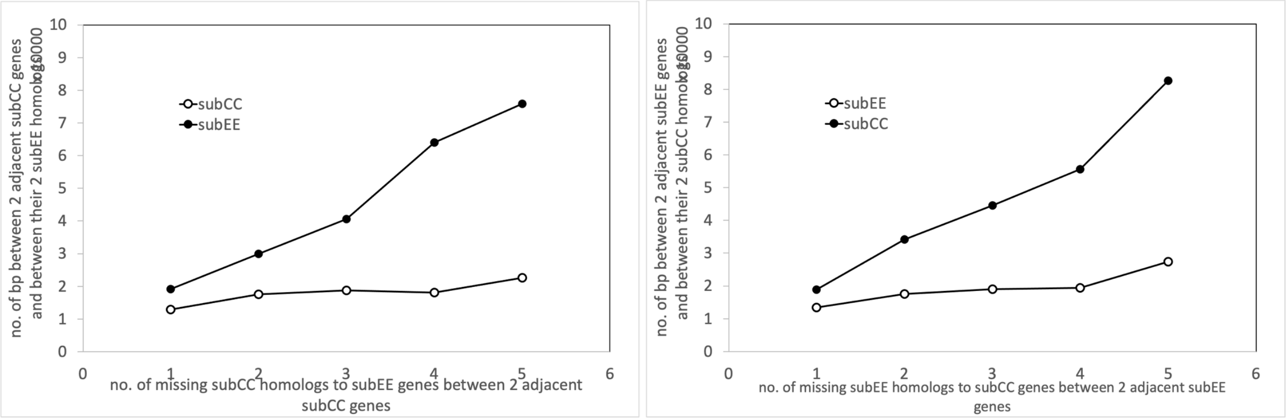

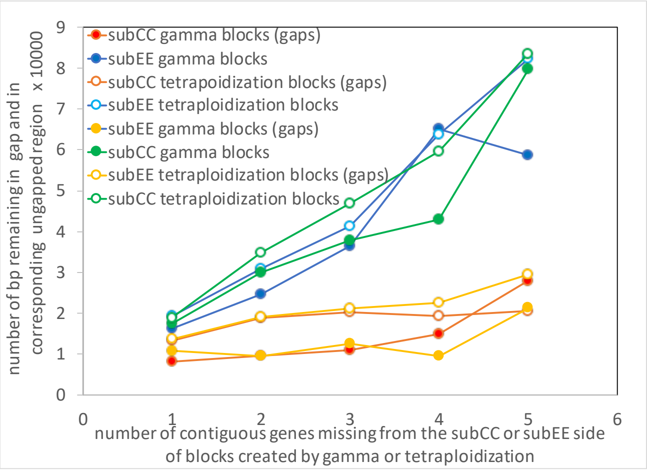


**b)**

**Fig S27. a)** *Lengths of DNA segments between genes made adjacent by fractionation of intervening genes still present in the opposite subgenome. Includes synteny blocks created by gamma and those created by tetraploidization. Left: subCC, right: subEE.* ***b)*** *Comparison of the synteny blocks created by the* gamma *event with those created by tetraploidization.*

**Fig. S26** represents in large measure the unmatched genes that were already absent in the progenitor genome. To try to distinguish between patterns of loss pre- and post-tetraploidization, we repeated the preceding analysis with restriction to the relatively small subset (11-13%) of genes unmatched in a subCC-subEE synteny block but matched in a synteny block between a subgenome and the opposite progenitor genome. The results in **Fig. S27** showed substantially the same pattern as in **Fig. S26**, indicating that the excisions were not just the product of many millions of years of independent evolution, including gene losses, in the progenitor genomes, but also reflect the fractionation process in the tetraploid. **Fig. S27b** compares results from the synteny blocks created by *gamma* with those created by Arabica tetraploidization.

The aggregate statistics represented in **Figs. S26, S27** do not rule out the possibility of occasional pseudogenization. Compared to the decreasing length between adjacent genes in **Fig. S26** the pattern in **Fig. S27** suggests more pseudogenization occurs during fractionation. PseudoPipe revealed thousands of pseudogenes in the *C. arabica* genome, but these were focused on retrotransposed pseudogenes and highly duplicated pseudogenes, presumably derived from specific functional classes such as housekeeping genes. There was no specific concern with sets of pseudogenes created almost simultaneously after a polyploidization event. The largest category in the output was that of gene fragments, but these were not controlled for syntenic context. Our search of the full PseudoPipe output found only a handful of pseudogenes in syntenic block gaps, out of many thousands of such gaps, that seemed to be derived from a duplicate of an unmatched gene in the opposite subgenome.

This prompted us to search directly for fragments of coding DNA in these gaps homologous to parts of unmatched genes in the opposite subgenome side of the synteny block. We required that the level of similarity between these parts and the fragments be greater than 92%, which is the average level of similarity we require for the coding sequence in synteny blocks created by the tetraploidization. (This constraint could be relaxed considerably without any large change in the kinds of fragments found.) Many unmatched genes had different parts connected to disjoint fragments, in which case we used the combined length of the matching parts to characterize the match.

**Fig. S28** shows that fragments of coding sequence, most of them less than 600 bp, and almost all less than 1000 bp – while the average annotated gene contains about 1500 bp – remain in the DNA for a few hundred cases of fractionation. Whether these ever passed through a pseudogene stage is dubious; the original gene seems to have been inactivated by a series of deletions of the coding sequence, not necessarily at the 5' or 3' end, but internal to the gene so that several fragments can be matched to different parts of the surviving gene in the opposite subgenome.

Summarizing, statistical evaluation of the massive duplicate gene cohorts created by speciation or polyploidization showed that pseudogenization is either a very rare process contributing to fractionation, or that does not result in much stable structure. At present, the clear impression is that fractionation simply excises the DNA of a gene or several contiguous genes. It is of course still possible that once a pseudogene is created, or a gene otherwise silenced, its DNA is immediately vulnerable to repeated small deletions, so that the pseudogene itself would be transient. The distinction between this and some single-event excision becomes a matter of semantics.

In summary, both fractionation in the tetraploid, or gene loss from the parent diploids in *Coffea* genomes, appears to have proceeded mostly by excision, resulting in shortening of the genome. Pseudogenization is either very rare, or transient. Some large gene fragments remain after an annotated gene could no longer be detected, in a substantial minority of cases, as detected by similarity to the surviving paralog.

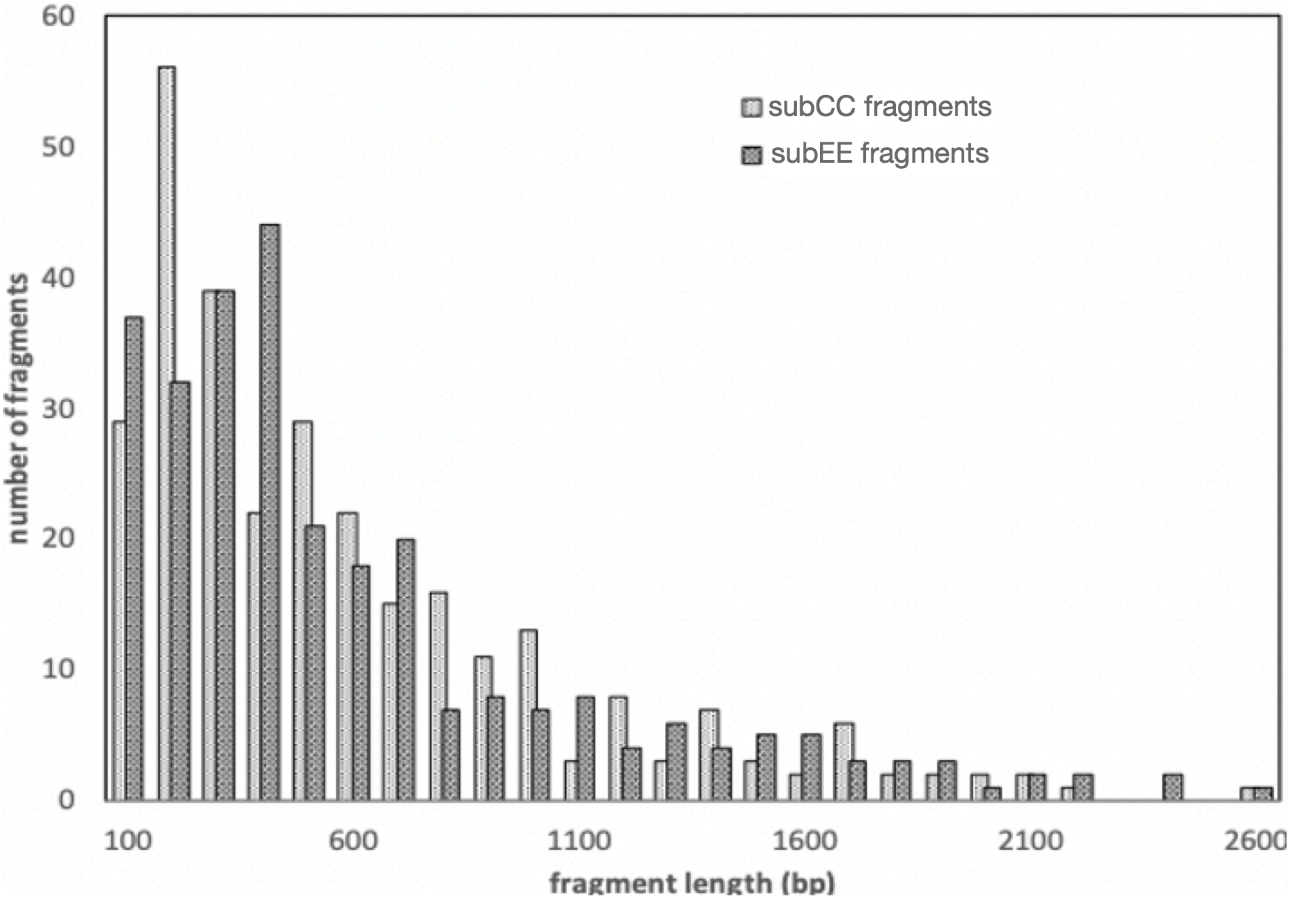


***Fig. S28.*** *Distribution of lengths of fragments matching part(s) of an unmatched gene in the opposite subgenome in a synteny block.*

The pattern of excision predominating over pseudogenization is not peculiar to *Coffea*. It is the common sequel to polyploidization and gene loss more generally (Yu et al., 2020).

##### 4.4 The evolution of syntenic homology from *gamma* to the present

We analyzed the homologous gene pairs in syntenic context as produced from the data on pairs of genomes using the SynMap procedure on the CoGe platform (Lyons & Freeling, 2008; Lyons et al., 2008). This allowed us to study the evolution of both paralogous and orthologous synteny blocks. We studied only genes within the region of the blocks, including gene pairs and singleton genes in each genome, that have lost their counterparts in the other genome due to fractionation or other gene loss. We used four genomes/genomic compartments, Robusta, Eugenioides, subCC and subEE, producing six comparisons of pairs, and four self-comparisons.

Certain operational definitions allowed us to analyze evolution coherently across all evolutionary eras. For more than 10 genes in a gap on one genome between two adjacent gene pairs, the synteny block was broken into two at that point. This was justified by the regular decrease in frequency in gap sizes from 0, 1, 2, until there were almost none of size 8, 9, or 10, except between neighboring synteny blocks, which can be separated by large numbers of unpaired genes in either of both of the genomes. We wished to study the nature of the distribution of gap size due to fractionation or gene deletion, and this convention avoids biasing estimates by inclusion of gaps produced by mechanisms other than fractionation. We used the default parameters of SynMap, except for the maximum number of non-duplicate genes interrupting any neighboring gene pairs, which we set at 10.

We used the "peaks" method (Yu et al., 2020) for the three events that generate duplicate genomes in the evolution of Arabica: the *gamma* hexaploidization, Robusta/Eugenioides speciation and Arabica tetraploidization (which is effectively a speciation of Robusta/subCC and of Eugenioides/subEE). In this method, the local modal values (peaks) of the distribution of the entire set of homologous gene pairs, as calculated by the R function geom_density, are estimates of the timing of the event.

###### 4.4.1 The sequence of evolutionary events

All ten self- and pairwise comparisons showed a cluster of homologous pairs dating from the early hexaploidization of the core eudicots. **Table S21** comparing averages over all pairs with less than 87% similarity, indicated tight clustering of these estimates, in terms of peak gene similarity (over CDS regions) and Ks.

The speciation of Robusta and Eugenioides generated orthologous gene pairs visible in the Robusta (or subCC) vs Eugenioides (or subEE) comparisons, as can be seen in **Figs S22-S25**. **Table S22**, presents the peak similarity and Ks for these comparisons.

The Arabica tetraploidization event, which for our purposes consists of the synchronous speciation of Robusta/subCC and Eugenioides/subEE, was detected as a peak of gene pairs in Robusta vs subCC and the Eugenioides vs subEE. These are depicted in **Table S23**.

The current best estimates of $\gamma$ and Robusta/Eugenioides speciation are on the order of 120 My and 4.5-7.2 My, while the Arabica tetraploidization is less than 1 My old. The similarity measures largely correspond with this timeline. The tetraploidy event seems to be 15-20% of the speciation age, suggesting an interval of 0.6-1.4 My. However, recent divergence estimates may suffer from difficulties interpreting similarities very close to 100% or very small Ks. Altogether, the rate of gene loss and the divergence time estimate from Ks point to similar time frames (**Fig. S29**).

It can be noted that in all of our comparisons, there has been a symmetry between Robusta and Eugenioides, and between subCC and subEE. If $\gamma$ fractionation rates or evolutionary divergence rates of CC and CE or subgenome dominance played a role, their effects must have been relatively small. Independent gene loss from Robusta-Eugenioides synteny blocks since speciation has been up to 25%.

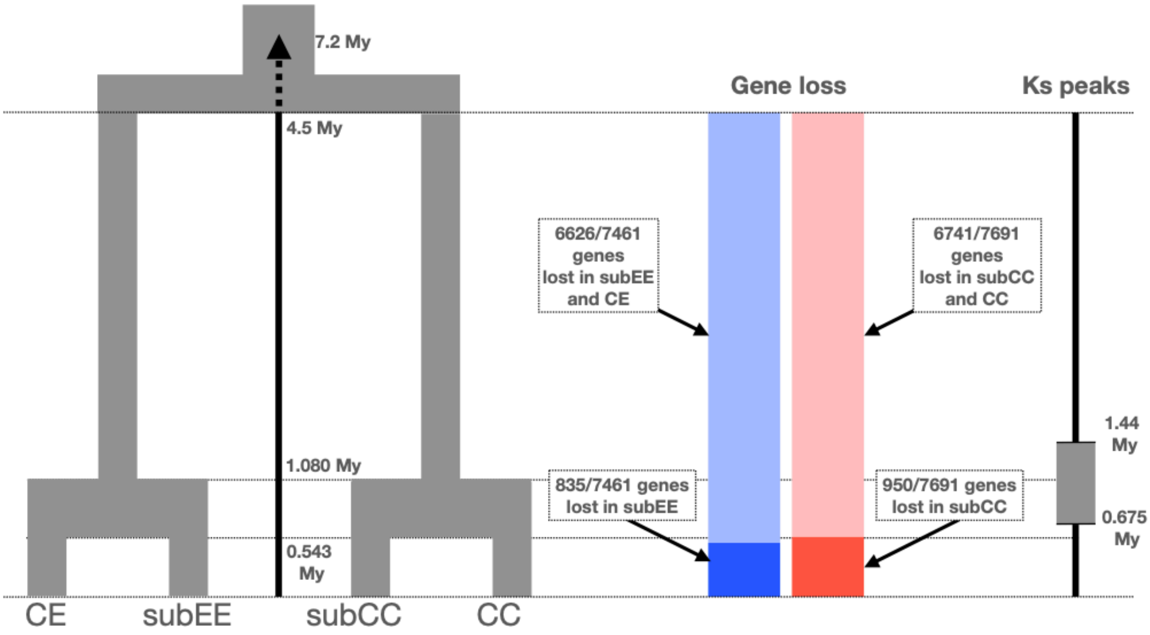


***Fig. S29.*** *A summary of the genome fractionation rate and divergence of syntenic gene models. The timing of the splits in the phylogeny (left) reflects the most recent estimates from (Bawin et al., 2020). The rate of gene loss (barplot) is presented as the percent of syntenic genes lost in the Eugenioides/subEE common ancestor (light blue) or only in subEE (blue). Similar analysis was carried out for Robusta-derived genomes, where the percent of genes lost in Robusta/subCC is shown in light red and genes lost only in subCC with dark red. The Ks peaks method (right) scales the divergence time between the subgenomes, estimated from number of synonymous mutations between syntenic genes, to the timing of the speciation event.*

To identify functional biases in the regions with different fractionation rates, we next identified the genes located in high fractionation (more than 45% of the genes have been lost) and low fractionation (more than 85% of genes have been retained) regions. Significant overlap with genes originating from whole genome duplications was found with regions with high retention rate, as well as an overlap between tandemly duplicated genes and regions with low retention rate (**Table S1**). Following the dosage balance hypothesis, we found significant overlap with genes in regulatory roles in high retention rate regions (**Tables S2-S3**), and genes associated with defense responses and secondary metabolism in low retention rate regions (**Tables S4-S5**).

#### 4.5 Gene family identification and sub-genome differential expression

##### 4.5.1 RNA-Seq analysis

To aid these analyses, the gene expression patterns for each subgenome in the beans of *Coffea arabica* (EMBL accession ID PRJEB24137) were analyzed at three maturation stages (Cheng et al., 2018): fruits at immature (Green; UG), intermediate (Yellow; UY), and mature stages (Red; UR). After quality control, reads were aligned against the *C. arabica* genome using STAR aligner v 2.7.10b (Dobin et al., 2013) with parameters (outFilterMultimapNmax=1, outFilterMismatchNmax=999, outFilterMismatchNoverLmax=0.016, outFilterMismatchNoverReadLmax=0.016). Differential expression analysis was first run in R using the DESeq2 package without taking into account homoeologous exchange, by calculating differential expression using green stage as control, that is, UY vs UG and UR vs UG. Differentially expressed genes (genes with FDR corrected p-values <0.05) were then split according to the subgenomes and tallied up (**Table S6**). Significance of subgenome-specific bias was tested with the Fisher exact test.

##### 4.5.2 Global expression bias

Subgenome-specific allelic biases were analyzed by first normalizing the raw counts from STAR into transcripts per million (tpm) values in R. The set of 10,281 genes that were syntenic between the subgenomes (syntenic alignments were found using SynMap in GoGe platform; see section 4.1 below), and did not show signs of homoeologous exchange (see **Supplementary section 4.1** below) in any of the accessions, were used in the subgenome dominance analysis. Statistical significance was tested using Student's t-test and the resulting *p*-values were corrected for multiple testing using Benjamini-Hochberg correction, using adjusted *p*-value < 0.05 as the threshold for significance. Statistical significance of the expression biases between subgenomes was tested using the Fisher exact test (**Table S7**). This transcriptome analysis highlighted the lack of dominance of one subgenome in the global contribution to the transcriptome, with only some genes expressed highly or exclusively from one of the two subgenomes.

##### 4.5.3 Gene family analysis

Gene family analysis was initiated by running OrthoFinder v2.3.2 with 21 species, plus the *Coffea* proteomes published in this work (**Table S24**).

Coffee flavors and health functions are mostly attributable to several characteristic secondary metabolites, including caffeine and terpenoids accumulated in coffee beans. InterProScan version 5.39-77.0 (Jones et al., 2014) was run on all protein sequences for target genomes. For fatty acid desaturase 2 (FAD2) sequences, proteins that included InterPro domain IPR005804 were extracted and OrthoFinder (Emms & Kelly, 2019) orthogroups containing these protein sequences were concatenated and used as input for the PASTA (Practical Alignment using Saté and TrAnsitivity) algorithm (Mirarab et al., 2014) to generate gene trees. InterPro family IPR005299 was used to identify N-methyltransferase (NMT) proteins, and InterPro domain IPR005630 was used to identify terpene synthase (TPS) proteins. InterProScan was run with default options and match calculations were run locally. PASTA version 1.9.0 was run with default options for protein data. FAD2, NMT, and TPS gene trees were created for protein sequences from all target genomes as well as a subsample of *Coffea arabica*, *C. canephora*, *C. eugenioides*, and *Arabidopsis thaliana*.

**NMT gene family.** Caffeine is a purine alkaloid responsible for the psychotropic properties of many plant derived beverages, such as coffee, tea, yerba mate and guaranà (Ashihara et al., 2017). The metabolite appeared in different plant genera through convergent evolution mediated by *NMT* gene duplication (Denoeud et al., 2014; Xia et al., 2017; Z. Xu et al., 2020). Caffeine biosynthesis is mediated by three different *NMT* genes (**Fig. S30**), all of which retain duplicates from Robusta- and Eugenioides-derived subgenomes in the Arabica genome. Furthermore, the *DXMT* gene, catalyzing the last step in caffeine biosynthesis, is tandemly duplicated in subEE, and both copies are expressed in fully ripe fruits at decreased levels compared to subCC, providing a molecular basis for the lower caffeine content of CA with respect to CC (Ashihara, 2008; Ashihara & Crozier, 1999; Campa et al., 2005). More generally, the *NMT* gene family in Arabica shows clear, though not extensive, mosaicism in subgenome-wise expression.

**TPS gene family.** Terpenes derived from geranyl diphosphate strongly contribute to coffee aroma (Del Terra et al., 2013). Here, we observed expression dominance by subEE for genes encoding $\alpha$-terpineol synthases and a putative nerolidol synthase. In contrast, a tandem duplication and expression of several putative $\alpha$-

farnesene synthases in subCC suggested expression dominance there, whereas the genes encoding for isoprene synthase also showed dominance for subCC, but only at the green maturation stage.

**FAD2 gene family.** Polyunsaturated fatty acids form an energy reserve in coffee beans and contribute to coffee flavor, aroma, and its shelf life (Speer & Kölling-Speer, 2006). *FATTY ACID DESATURASE 2* (*FAD2*) encodes the key enzymes that desaturate oleic acid to linoleic acid, the major unsaturated fatty acid in coffee. *Arabidopsis* encodes a single, constitutively expressed *FAD2* gene, whereas some oil-producing plants, and coffee, have multiple copies of *FAD2*, some of which are mainly expressed in seeds. Based on phylogenetic analyses, these seed-type *FAD2*s evolved from housekeeping FAD2-encoding genes by gene duplication (Dar et al., 2017; Hernández et al., 2005). In Arabica, the housekeeping *FAD2* syntelogs were present in both subgenomes, with expression dominated by subEE, whereas seed-type *FAD2* was duplicated in both subgenomes, where overall, the duplicates showed expression patterns suggesting dominance by subCC. Interestingly, the genic region encoding the N-terminal signal peptide of the only highly expressed subEE homolog (Cara002g032150) differed from those of the other seed-type paralogs; this could result, for example, from a gene conversion event (Deb et al., 2023; Guo et al., 2014).

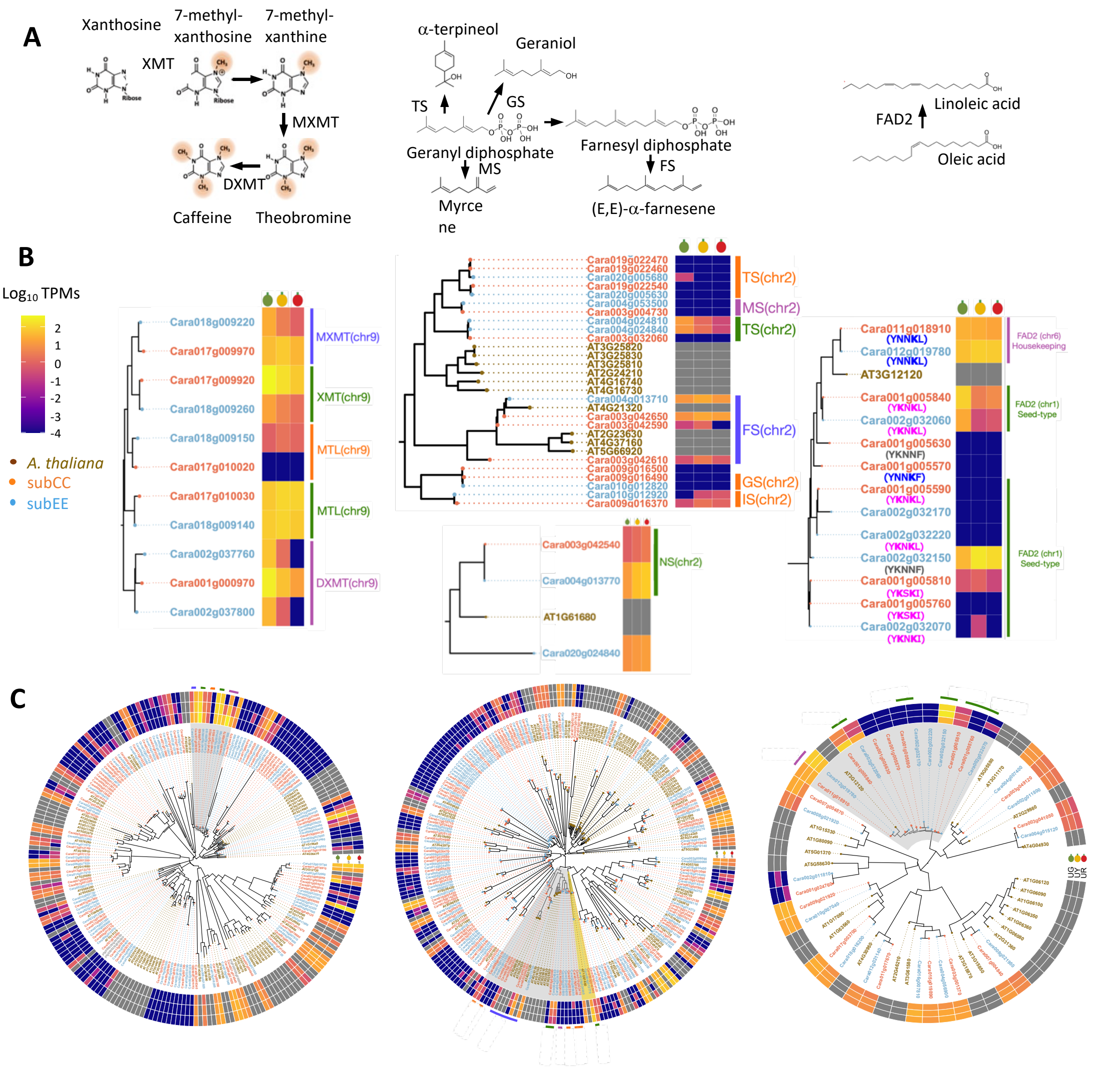


*Figure S30. Composition and expression of* exemplar Arabica *gene families contributing to bean quality traits. A. Schematic biosynthesis of caffeine (left), terpenoids (middle), and unsaturated fatty acids (right)* ***B.*** *Phylogenies and expression during fruit development of* CA *genes for N-methyltransferases (NMTs) mediating caffeine biosynthesis (left), terpene synthases (TPS) (middle), and fatty acid dehydrogenase 2 (FAD2) (right).*

*RNA sequencing was carried out for three biological replicates from three different fruit maturation stages (green, yellow, and red) of the K7 cultivar.* ***C.*** *Genome-wide NMT (left), TPS (middle), and FAD2 (right) phylogenies and expression patterns during fruit development. Genes located in the two subgenomes are indicated by font color; subCC (red) and subEE (blue). Grey areas highlight the parts of phylogenies shown in B. XMT: xanthosine methyltransferase; MXMT: 7-methylxanthine methyltransferase; DXMT: 1,7-dimethylxanthine methyltransferase; MTL: N-methyltransferase-like; FS: (E,E)-a-farnesene synthase; GS: Geraniol synthase; IS: Isoprene synthase; MS: myrcene synthase; TS: (-)-a-terpineol synthase; FAD2: Fatty acid desaturase 2.*

###### 4.5.3.1 Evolutionary analysis of the FAD2 family

Fatty acid desaturases (FADs) catalyze the desaturations of fatty acids, which are important for coffee's fragrance, body and flavor (Anagbogu et al., 2021). In the endoplasmic reticulum (ER), FAD2 and FAD3 sequentially convert oleic acid (18:1) to linoleic acid (18:2) and linolenic acid (18:3) (Okuley et al., 1994; Yadav et al., 1993).

The FAD2 gene is tandemly duplicated multiple times in the *Coffea arabica* genome, which is not observed with other FADs (Denoeud et al., 2014). The expansion of the FAD2 gene family is likely responsible for the unusually high amount of linoleic acid (18:2) in *C. arabica* seeds, where it accounts for over 40% of total lipids (Dussert et al., 2008). While Arabidopsis has only one FAD2 gene, oilseed crops such as sesame, corn, canola, olive, soybean, sunflower and cotton have more than one copy of FAD2 (Dar et al., 2017).

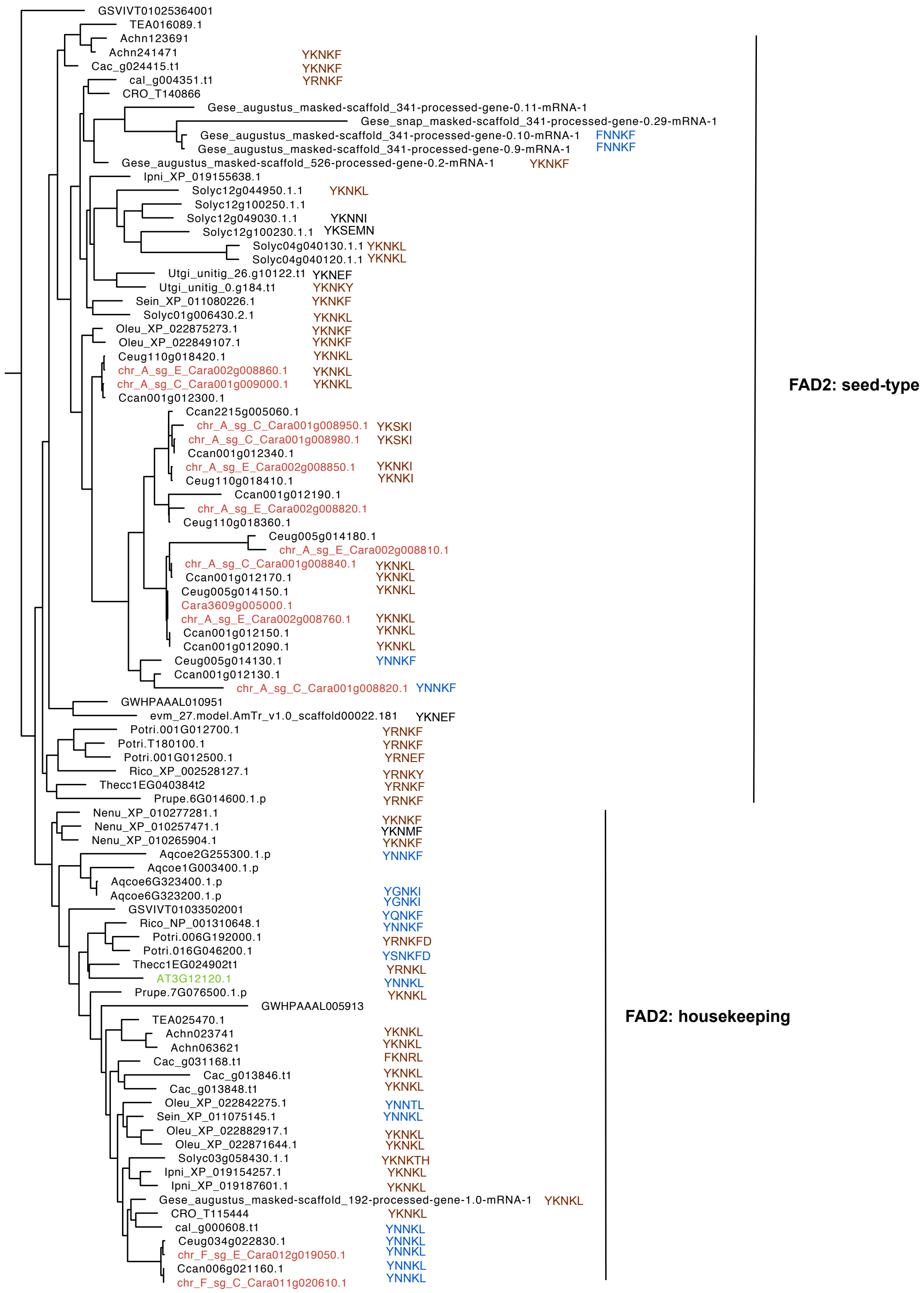


***Fig. S31.*** *A phylogenetic tree of FAD2 from 22 plant species. FAD2s are grouped into two clades: the housekeeping and the seed-type. Arabidopsis FAD2 (AT3G12120) is marked in green. Coffea arabica FAD2s are labeled in red. FAD2s in other plant species are black. The C-terminal ER retrieval peptides are next to each gene. The C-terminal sequences in blue have the "YNNKX" motif while the ones in brown have the "YKNKX" or "YRNKX" motif, with "X" being hydrophobic amino acids. Annotations of species used: GSVIV: Vitis vinifera; Cara: C. arabica; Ceug: C. eugenioides; Ccan: C. canephora; TEA: Camellia sinensis; Achn: Actinidia chinensis ; Cac: Camptotheca acuminata; cal: Calotropis gigantea; CRO: Catharanthus roseus; Gese: Gelsemium sempervirens; Ipni: Ipomaea nil; Solyc: Solanum lycopersicum; Utgi: Utricularia gibba;*

*Sein: Sesamum indicum; Oleu: Olea europaea; GWHPAAAL: Eucommia ulmoides; evm_27.model.AmTr: Amborella trichopoda; Potri: Populus trichocarpa; Rico: Ricinus communis; Thecc: Theobroma cacao.*

Phylogenetic analysis of FAD2 genes in *C. arabica* revealed two distinct clades: one housekeeping and one seed-type (Dar et al., 2017; Hernández et al., 2005); the split was also visible in Orthofinder analyses (Fig. S30). The two housekeeping FAD2 genes in *C. arabica* were indeed expressed at constantly high levels in all stages of fruit development. Among the seed-type FAD2s, Cara001g005840 and Cara002g032060 were highly expressed during early fruit development but gradually reduced as the fruit ripens. The other seed-type FAD2s showed diverse expression patterns during fruit development.

We also noticed that the housekeeping and the seed-type FAD2s have different C-terminal ER retrieval signals. The actively expressed housekeeping FAD2 in *C. arabica*, similar to the FAD2 of Arabidopsis, has a C-terminal retrieval signal YNNKX that is necessary and sufficient to prevent the movement of FAD2 from the ER to the Golgi (McCartney et al., 2004) **(Fig. S30**). However, the actively expressed *C. arabica* seed-type FAD2s have a di-lysine C-terminal motif YKNKX as the classic ER retrieval signal (Jackson et al., 1990). Further, in the phylogenetic analysis of FAD2 genes from 22 different plant species, the putative seed-type FAD2 genes almost always have the C-terminal ER retrieval signal with two positively charged amino acids: either YKNKX or YRNKX. In contrast, putative housekeeping FAD2 genes of these species contain either the di-lysine YKNKX or the mono-lysine YNNKX motifs (Hernández et al., 2005). The conclusion that housekeeping and seed-type FAD2 genes can be distinguished by their C-terminal ER retrieval signal may well be extended to other plants.

### 5 Homoeologous exchange

Homoeologous exchange analysis was carried out in a similar manner as (Bird et al., 2021). To assess possible homoeologous exchange between the two subgenomes, a pseudoreference was first made by concatenating the modern representatives of the diploid progenitors into one common reference. Next, fastq files consisting of reads where PCR duplicates had been removed were obtained from .bam files from Picard v2.18.14 MarkDuplicates (see **Supplementary section 5.2**. for preprocessing of sequencing data).

A set of homoelogous gene pairs was obtained by syntenic alignments between the *C. canephora* and *C. eugenioides* assemblies using the SynMap tool in CoGe. Additionally, syntenic alignments were carried out between *Coffea arabica* subCC vs. *C. canephora* and *C. arabica* subEE vs. *C. eugenioides*. Since the SynMap algorithm collapses tandemly duplicated genes into one representative, the syntelog pairs were further filtered for tandemly duplicated genes. Coverages on syntelog pairs were calculated using BEDtools bedcov (Quinlan & Hall, 2010). Differential coverage across the chromosomes was visualized using custom R scripts. To reduce noise, a sliding window of 10 genes was used to calculate the average coverage along chromosomes. The allele balance was calculated as A= 4*((CC/(CC+EE))-0.5), where CC and EE are the subCC and subEE syntelog coverages, respectively. Allele balances < -1.5 or > 1.5 were considered homozygous for EE, or CC, respectively, while balances <0.5 and > -0.5 were considered equal.

***Coffea canephora* allele balances.** As a control, we first mapped the *C. canephora* accessions against the pseudoreference. The analyses showed balances corresponding to homozygous for CC, as expected (**Fig S32)**.

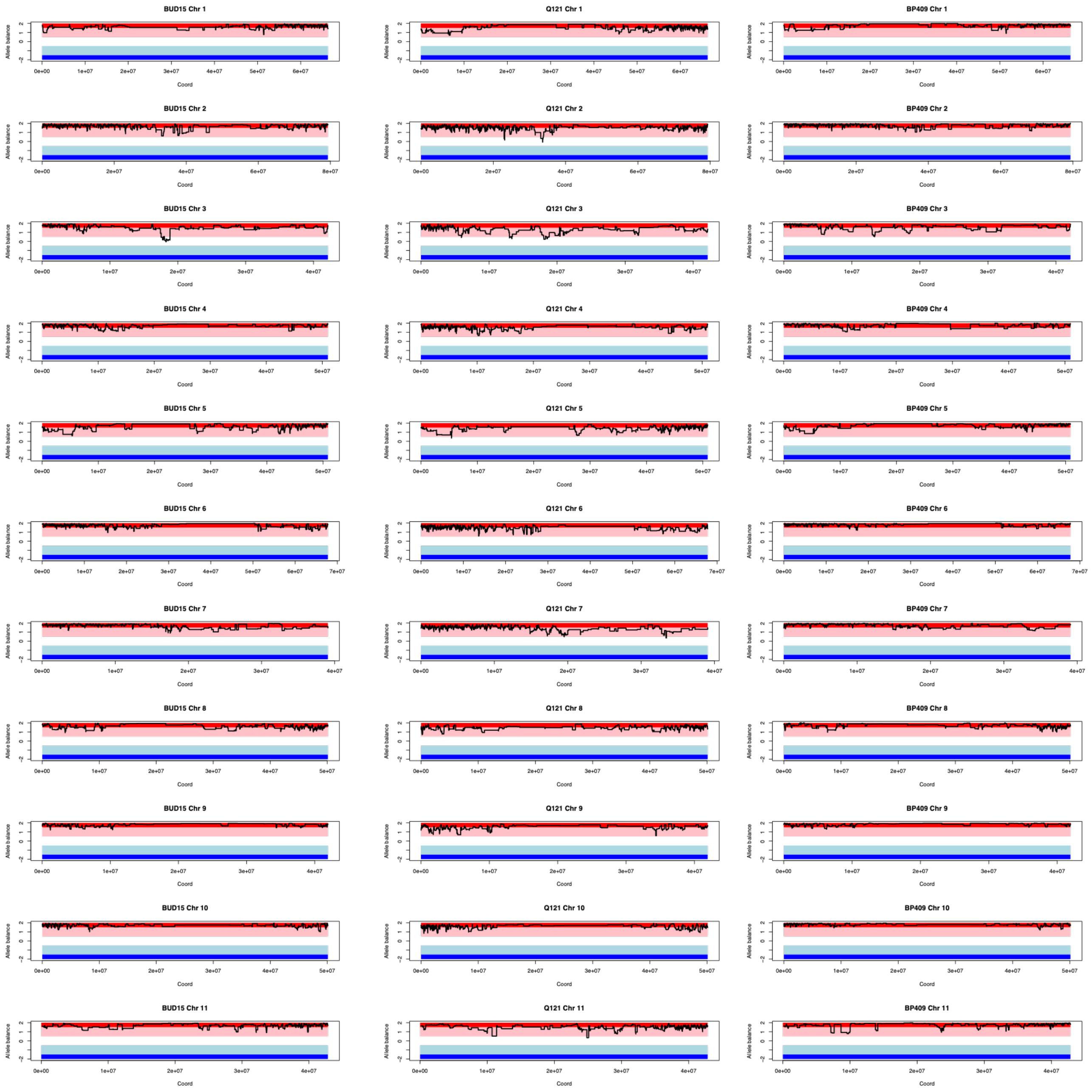


***Fig S32.*** *Homoeologous exchange analysis for* Coffea canephora *accessions BUD15, Q121 and BP409. The dark red region indicates 4:0 allele balance in favor of subCC, while the pink region illustrates 3:1, white 2:2, light blue 1:3 and dark blue 0:4 balances, respectively. The black line indicates the observed allele balances in syntelog gene pairs.*

***Coffea eugenioides* allele balances.** As a second control, we next mapped the *Coffea eugenioides* wild accession DA56 against the pseudoreference (**Fig S33**). The analyses showed balances corresponding to homozygous for subEE, as expected. However, there were some regions in chromosomes 1, 9 and 11 where the mapping coverage was equal between the two progenitors. This could be due to ancestral polymorphism (incomplete lineage sorting) or introgression in the DA56 lineage. However, distinguishing between these and other alternatives will require further research, preferably with more *C. eugenioides* accessions.

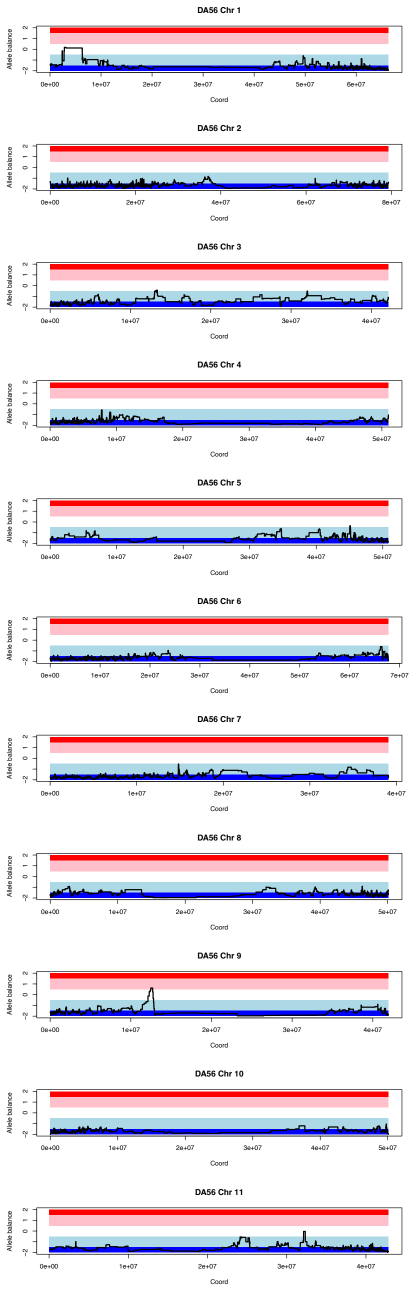


***Fig S33.*** *Homoeologous exchange analysis for* Coffea eugenioides *accession DA56. The dark red region indicates 4:0 allele balance in favor of subCC, while the pink region illustrates 3:1, white 2:2, light blue 1:3 and dark blue 0:4 balances, respectively. The black line indicates the observed allele balances in syntelog gene pairs.*

***Coffea arabica* wild accessions.** The first case study in the HE analysis was the Arabica wild individuals. Overall, the allele balances showed variation between 1:3 – 2:2 balance, except for one region, the beginning of chromosome 7, which was fixed for subEE (**Fig S34**). The high number of regions with 1:3 / 3:1 ratios suggested relatively frequent homoeologous exchanges between the two subgenomes. Several accessions showed similar patterns from shared homoeologous exchange events, resulting from common ancestry (see relationship analysis with KING; **Fig 4**; **Fig S48**; **Table S25-S26**).

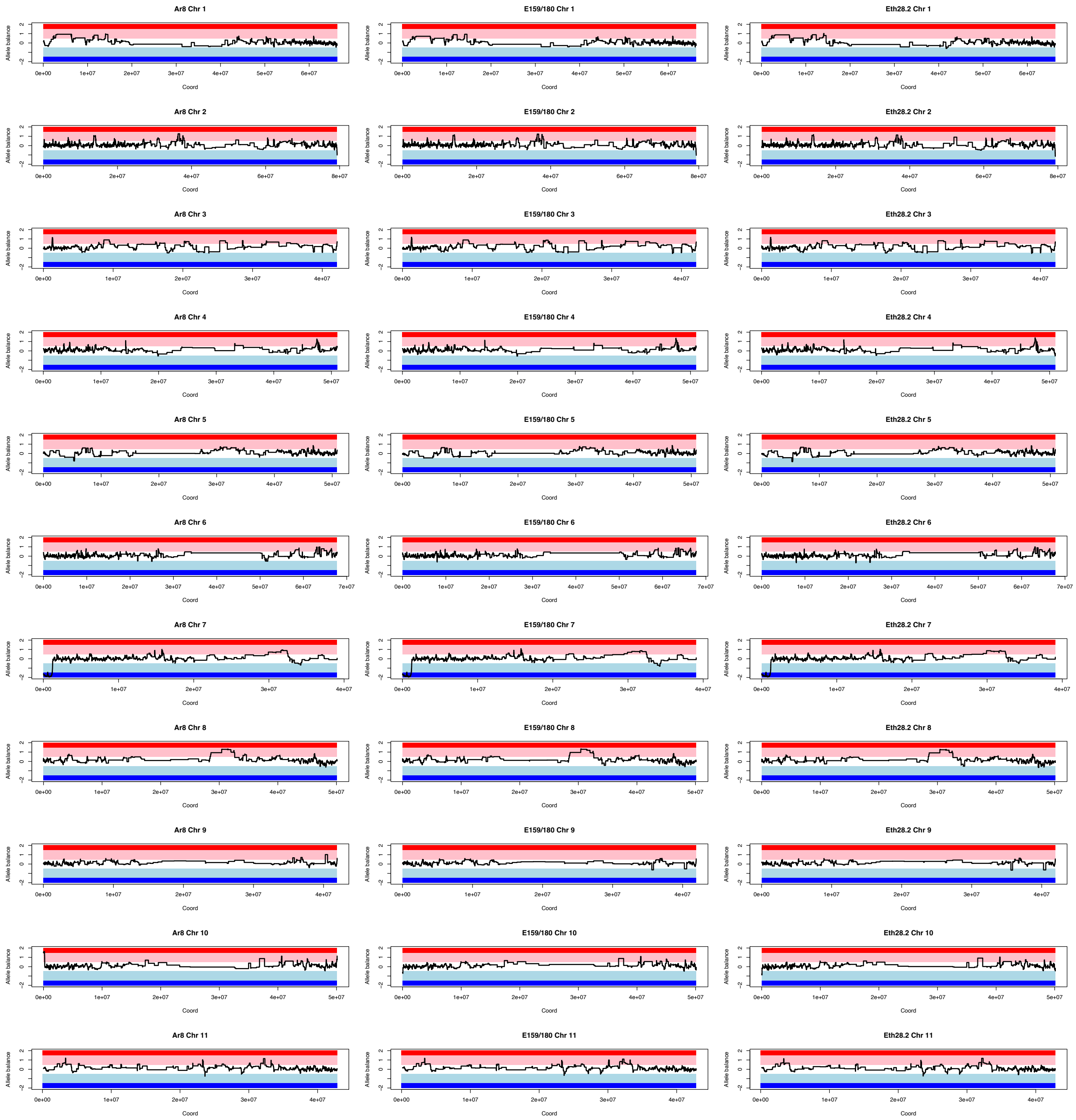
Ar8 Chr 1
E159/180 Chr 1
Eth28.2 Chr 1
Ar8 Chr 2
E159/180 Chr 2
Eth28.2 Chr 2
Ar8 Chr 3
E159/180 Chr 3
Eth28.2 Chr 3
Ar8 Chr 4
E159/180 Chr 4
Eth28.2 Chr 4
Ar8 Chr 5
E159/180 Chr 5
Eth28.2 Chr 5
Ar8 Chr 6
E159/180 Chr 6
Eth28.2 Chr 6
Ar8 Chr 7
E159/180 Chr 7
Eth28.2 Chr 7
Ar8 Chr 8
E159/180 Chr 8
Eth28.2 Chr 8
Ar8 Chr 9
E159/180 Chr 9
Eth28.2 Chr 9
Ar8 Chr 10
E159/180 Chr 10
Eth28.2 Chr 10
Ar8 Chr 11
E159/180 Chr 11
Eth28.2 Chr 11
Allele balance
Coord

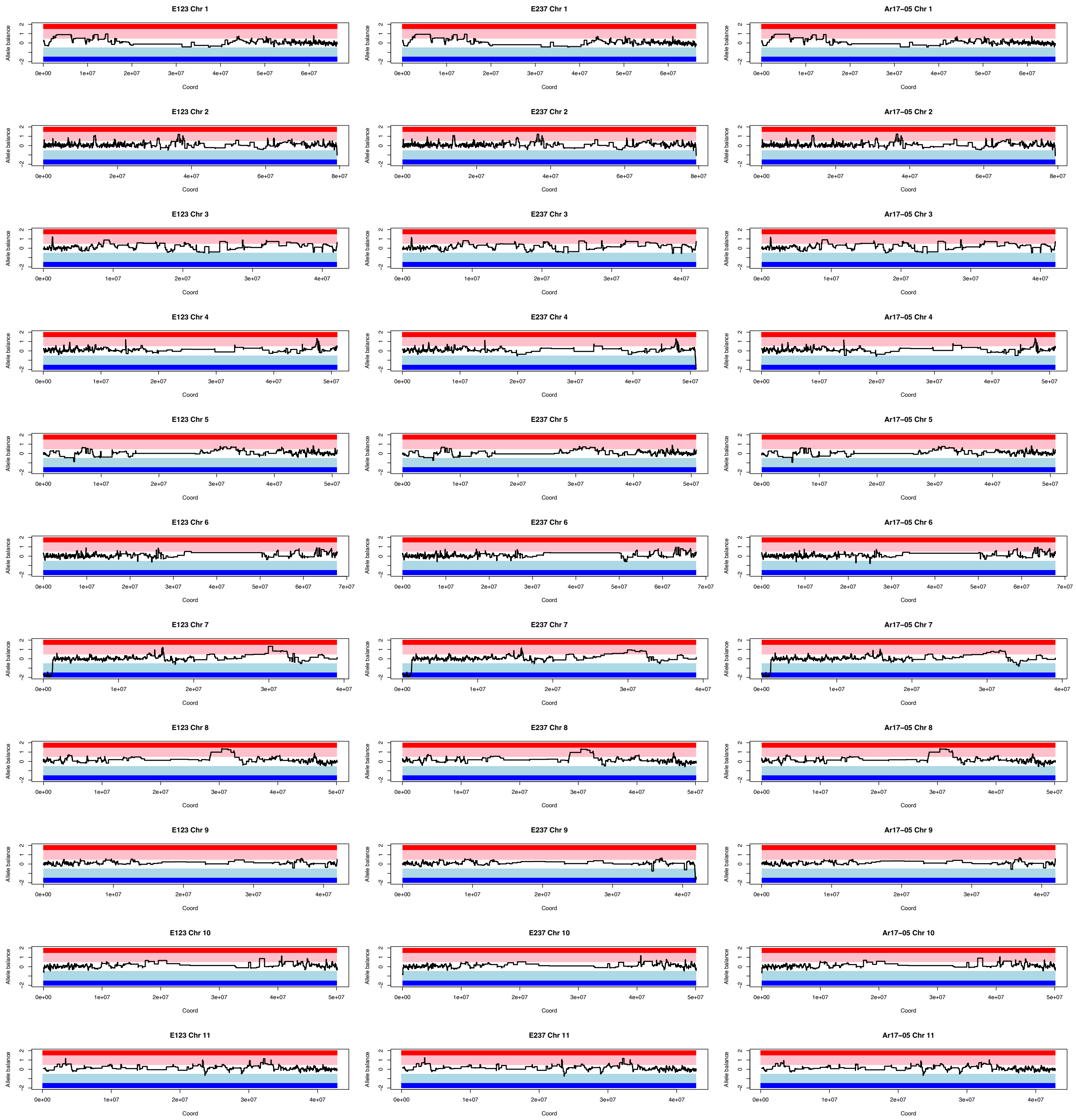
E123 Chr 1
E237 Chr 1
Ar17–05 Chr 1
E123 Chr 2
E237 Chr 2
Ar17–05 Chr 2
E123 Chr 3
E237 Chr 3
Ar17–05 Chr 3
E123 Chr 4
E237 Chr 4
Ar17–05 Chr 4
E123 Chr 5
E237 Chr 5
Ar17–05 Chr 5
E123 Chr 6
E237 Chr 6
Ar17–05 Chr 6
E123 Chr 7
E237 Chr 7
Ar17–05 Chr 7
E123 Chr 8
E237 Chr 8
Ar17–05 Chr 8
E123 Chr 9
E237 Chr 9
Ar17–05 Chr 9
E123 Chr 10
E237 Chr 10
Ar17–05 Chr 10
E123 Chr 11
E237 Chr 11
Ar17–05 Chr 11
Allele balance
Coord

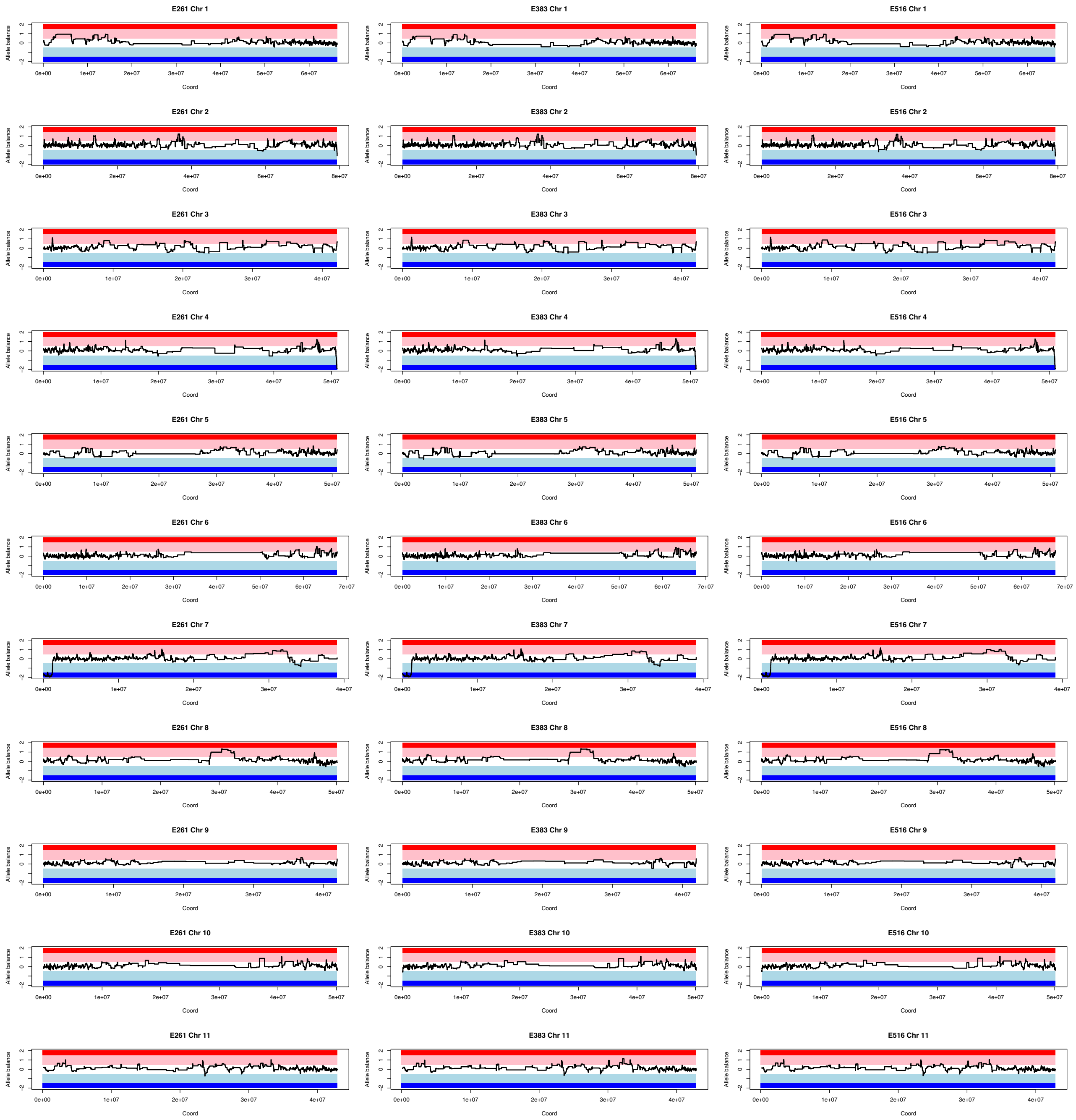
E261 Chr 1
E383 Chr 1
E516 Chr 1
E261 Chr 2
E383 Chr 2
E516 Chr 2
E261 Chr 3
E383 Chr 3
E516 Chr 3
E261 Chr 4
E383 Chr 4
E516 Chr 4
E261 Chr 5
E383 Chr 5
E516 Chr 5
E261 Chr 6
E383 Chr 6
E516 Chr 6
E261 Chr 7
E383 Chr 7
E516 Chr 7
E261 Chr 8
E383 Chr 8
E516 Chr 8
E261 Chr 9
E383 Chr 9
E516 Chr 9
E261 Chr 10
E383 Chr 10
E516 Chr 10
E261 Chr 11
E383 Chr 11
E516 Chr 11
Allele balance
Coord

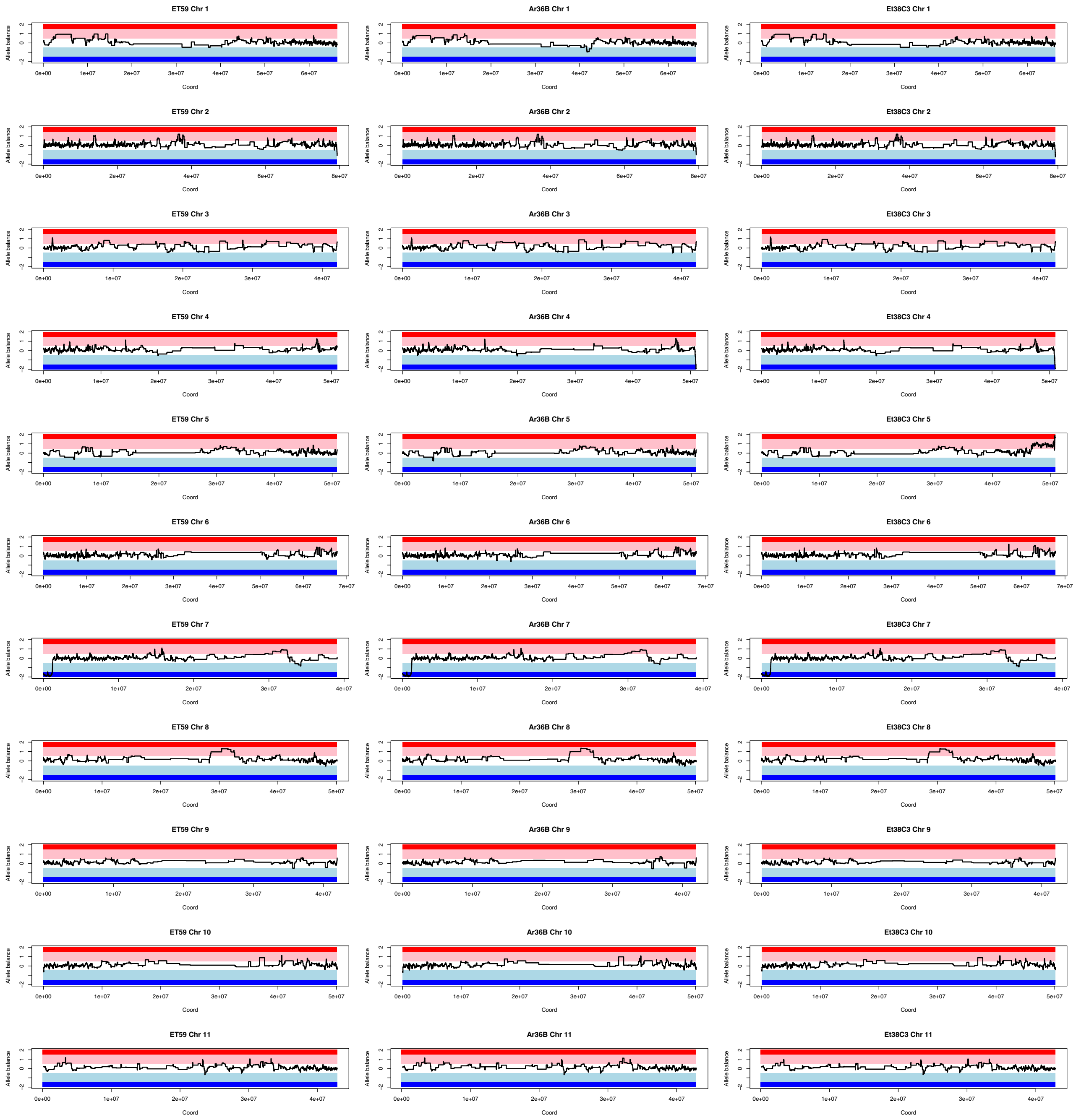
ET59 Chr 1
Ar36B Chr 1
Et38C3 Chr 1
ET59 Chr 2
Ar36B Chr 2
Et38C3 Chr 2
ET59 Chr 3
Ar36B Chr 3
Et38C3 Chr 3
ET59 Chr 4
Ar36B Chr 4
Et38C3 Chr 4
ET59 Chr 5
Ar36B Chr 5
Et38C3 Chr 5
ET59 Chr 6
Ar36B Chr 6
Et38C3 Chr 6
ET59 Chr 7
Ar36B Chr 7
Et38C3 Chr 7
ET59 Chr 8
Ar36B Chr 8
Et38C3 Chr 8
ET59 Chr 9
Ar36B Chr 9
Et38C3 Chr 9
ET59 Chr 10
Ar36B Chr 10
Et38C3 Chr 10
ET59 Chr 11
Ar36B Chr 11
Et38C3 Chr 11
Allele balance
Coord

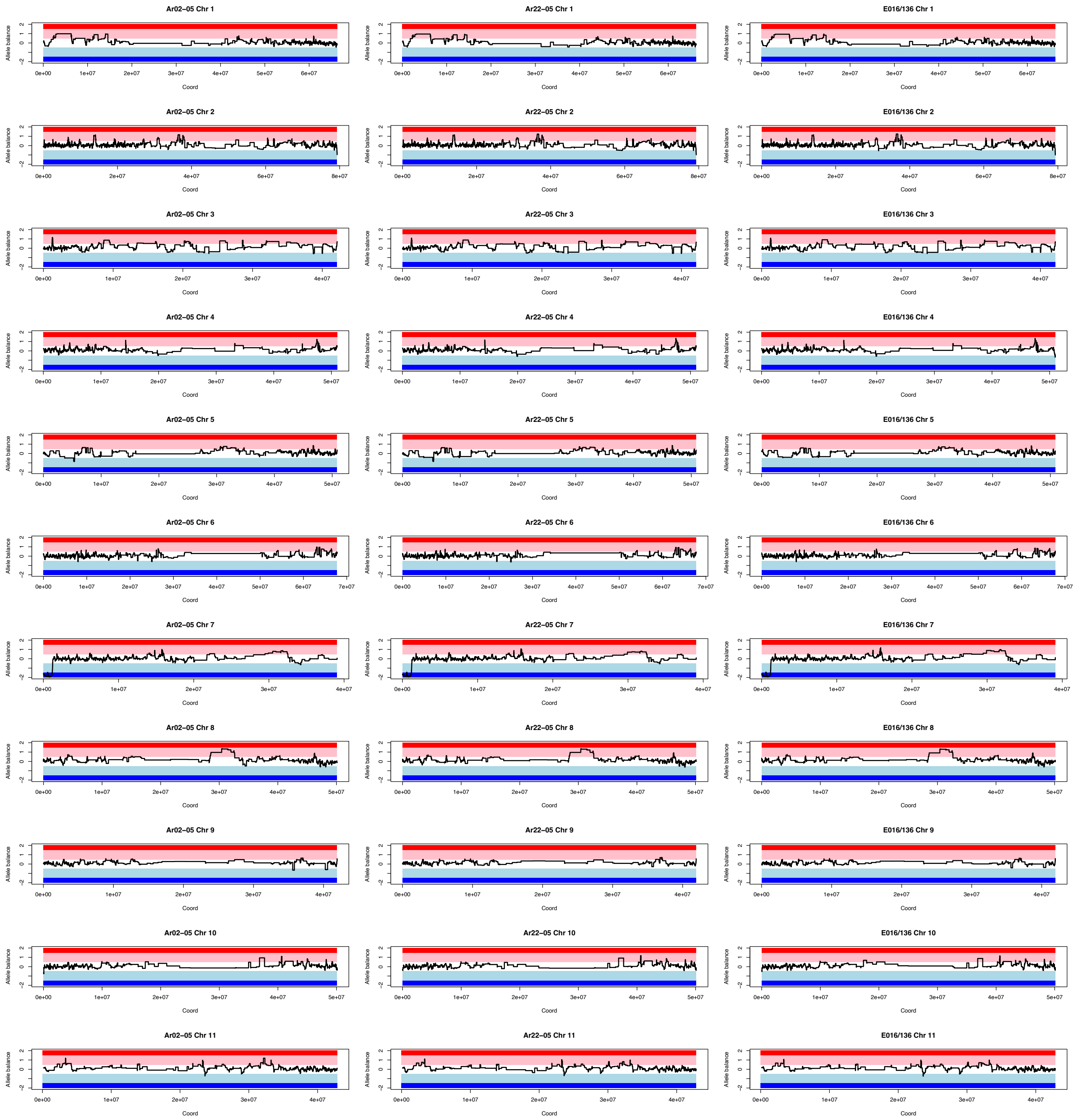
Ar02–05 Chr 1
Ar22–05 Chr 1
E016/136 Chr 1
Ar02–05 Chr 2
Ar22–05 Chr 2
E016/136 Chr 2
Ar02–05 Chr 3
Ar22–05 Chr 3
E016/136 Chr 3
Ar02–05 Chr 4
Ar22–05 Chr 4
E016/136 Chr 4
Ar02–05 Chr 5
Ar22–05 Chr 5
E016/136 Chr 5
Ar02–05 Chr 6
Ar22–05 Chr 6
E016/136 Chr 6
Ar02–05 Chr 7
Ar22–05 Chr 7
E016/136 Chr 7
Ar02–05 Chr 8
Ar22–05 Chr 8
E016/136 Chr 8
Ar02–05 Chr 9
Ar22–05 Chr 9
E016/136 Chr 9
Ar02–05 Chr 10
Ar22–05 Chr 10
E016/136 Chr 10
Ar02–05 Chr 11
Ar22–05 Chr 11
E016/136 Chr 11
Allele balance
Coord

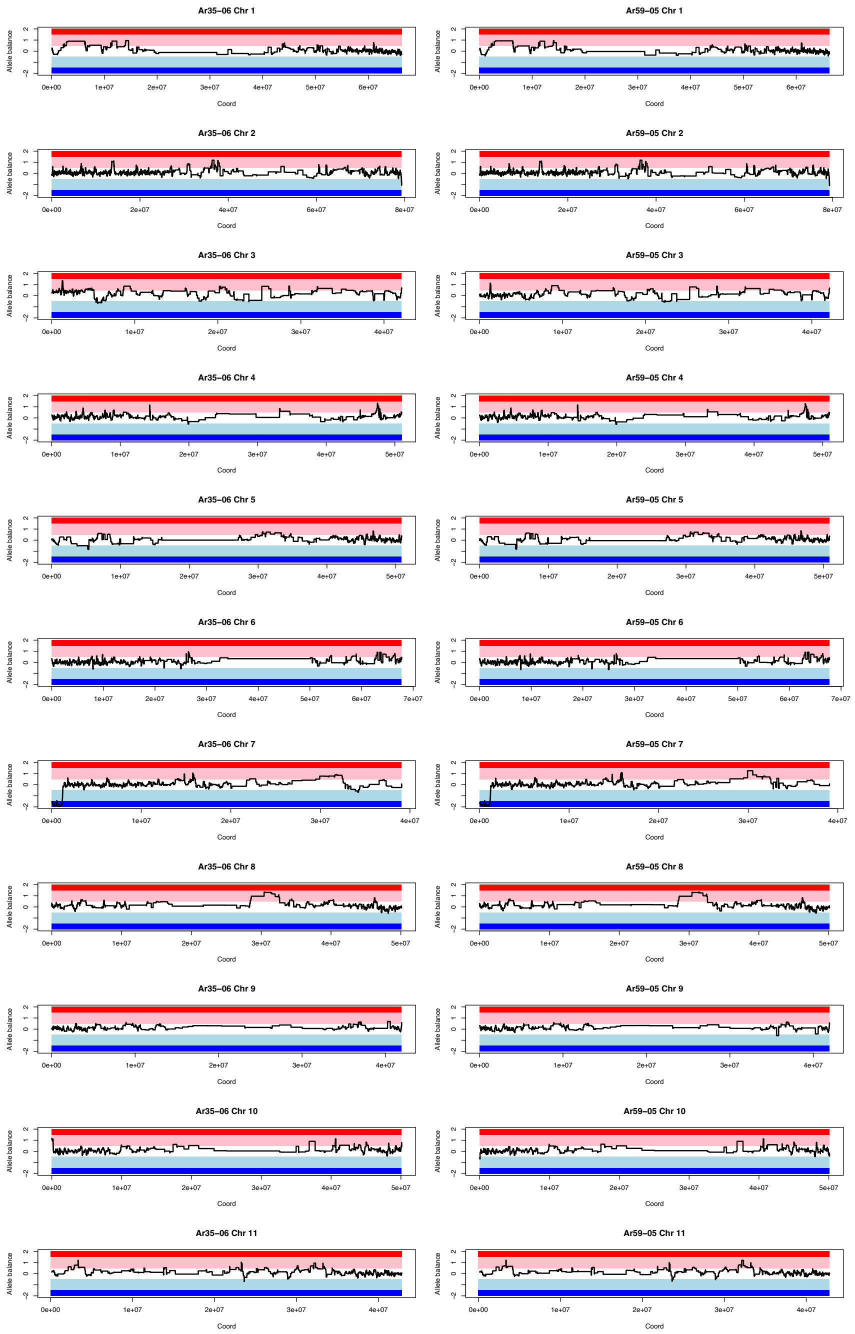


***Fig S34.*** *Homoeologous exchange analysis for* Coffea arabica *wild accessions. For each subplot, the title gives the accession ID followed by chromosome number. The dark red region indicates 4:0 allele balance in favor of subCC, while the pink region illustrates 3:1, white 2:2, light blue 1:3 and dark blue 0:4 balances, respectively. The black line indicates the observed allele balances in syntelog gene pairs.*

**Cultivated *Coffea arabica*.** Next, we analyzed the cultivated Arabica accessions from the Bourbon, Typica and Geisha lines (**Fig S35**). The cultivars showed shared homoeologous exchange events due to their close relatedness; this was also true for the Geisha accession. Interestingly BMJM had an accession-specific homoeologous exchange event from subEE in chromosome 1.

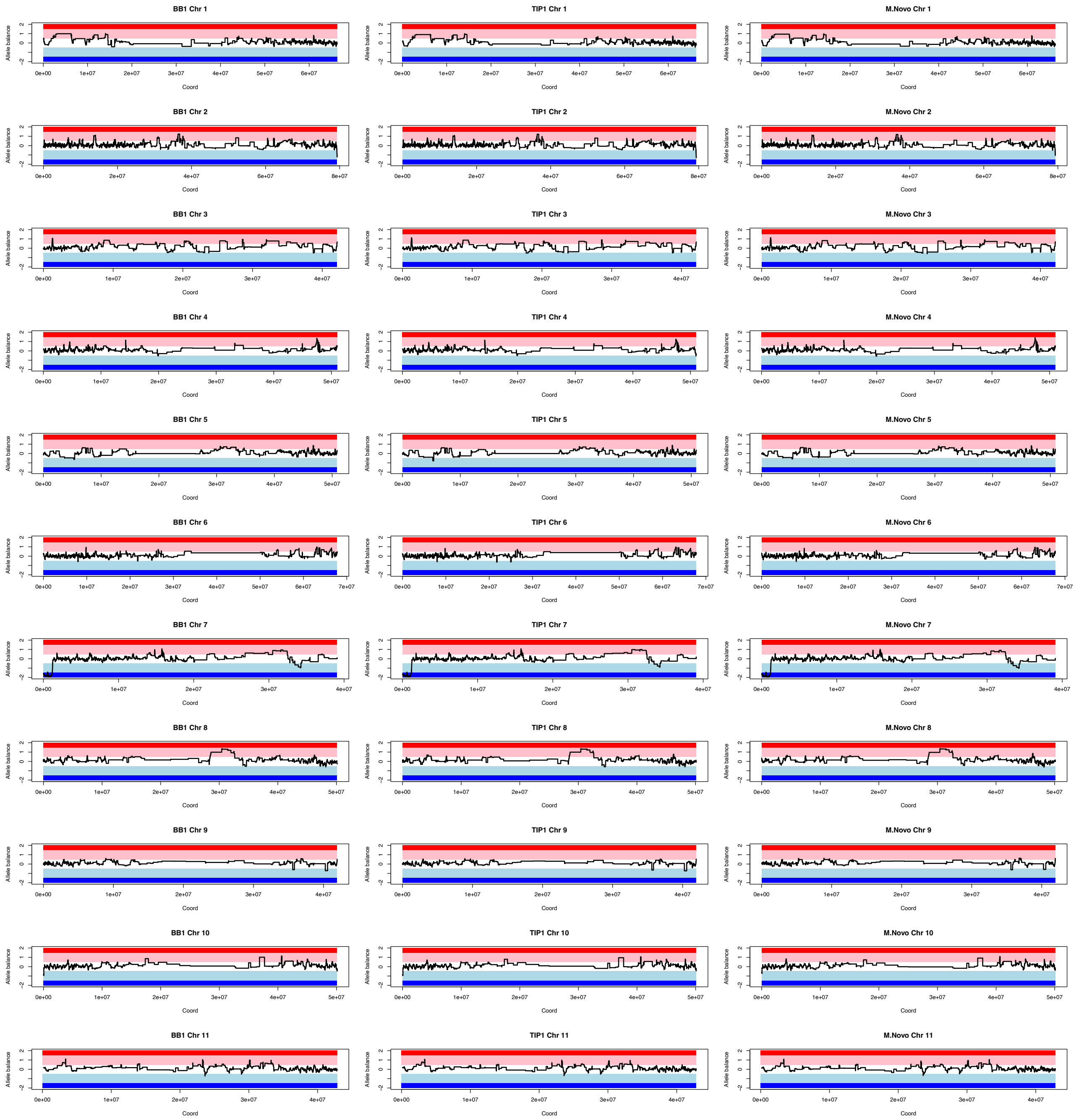
BB1 Chr 1
TIP1 Chr 1
M.Novo Chr 1
BB1 Chr 2
TIP1 Chr 2
M.Novo Chr 2
BB1 Chr 3
TIP1 Chr 3
M.Novo Chr 3
BB1 Chr 4
TIP1 Chr 4
M.Novo Chr 4
BB1 Chr 5
TIP1 Chr 5
M.Novo Chr 5
BB1 Chr 6
TIP1 Chr 6
M.Novo Chr 6
BB1 Chr 7
TIP1 Chr 7
M.Novo Chr 7
BB1 Chr 8
TIP1 Chr 8
M.Novo Chr 8
BB1 Chr 9
TIP1 Chr 9
M.Novo Chr 9
BB1 Chr 10
TIP1 Chr 10
M.Novo Chr 10
BB1 Chr 11
TIP1 Chr 11
M.Novo Chr 11
Allele balance
Coord

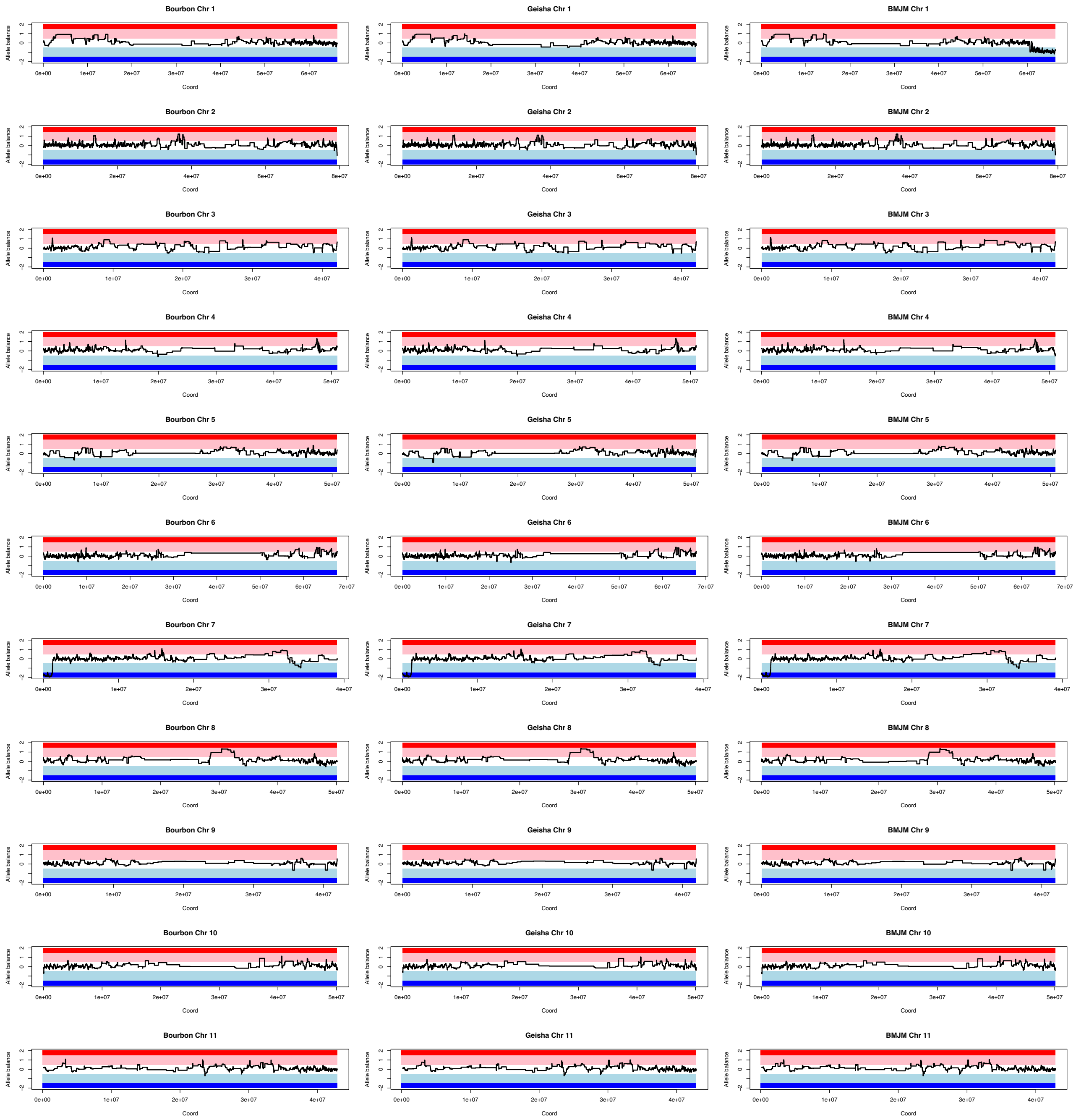
Bourbon Chr 1
Geisha Chr 1
BMJM Chr 1
Bourbon Chr 2
Geisha Chr 2
BMJM Chr 2
Bourbon Chr 3
Geisha Chr 3
BMJM Chr 3
Bourbon Chr 4
Geisha Chr 4
BMJM Chr 4
Bourbon Chr 5
Geisha Chr 5
BMJM Chr 5
Bourbon Chr 6
Geisha Chr 6
BMJM Chr 6
Bourbon Chr 7
Geisha Chr 7
BMJM Chr 7
Bourbon Chr 8
Geisha Chr 8
BMJM Chr 8
Bourbon Chr 9
Geisha Chr 9
BMJM Chr 9
Bourbon Chr 10
Geisha Chr 10
BMJM Chr 10
Bourbon Chr 11
Geisha Chr 11
BMJM Chr 11
Allele balance
Coord

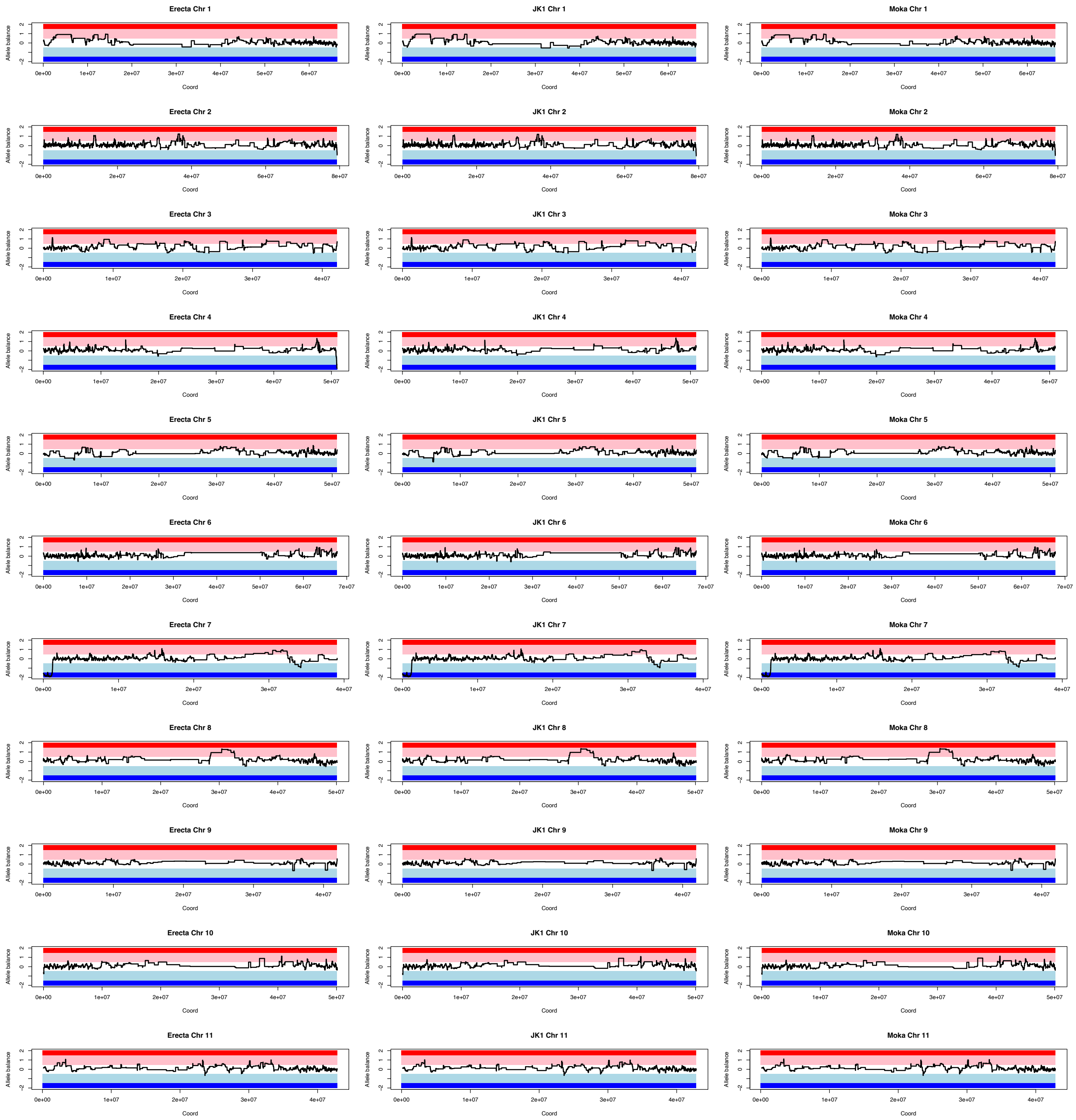
Erecta Chr 1
JK1 Chr 1
Moka Chr 1
Erecta Chr 2
JK1 Chr 2
Moka Chr 2
Erecta Chr 3
JK1 Chr 3
Moka Chr 3
Erecta Chr 4
JK1 Chr 4
Moka Chr 4
Erecta Chr 5
JK1 Chr 5
Moka Chr 5
Erecta Chr 6
JK1 Chr 6
Moka Chr 6
Erecta Chr 7
JK1 Chr 7
Moka Chr 7
Erecta Chr 8
JK1 Chr 8
Moka Chr 8
Erecta Chr 9
JK1 Chr 9
Moka Chr 9
Erecta Chr 10
JK1 Chr 10
Moka Chr 10
Erecta Chr 11
JK1 Chr 11
Moka Chr 11
Allele balance
Coord

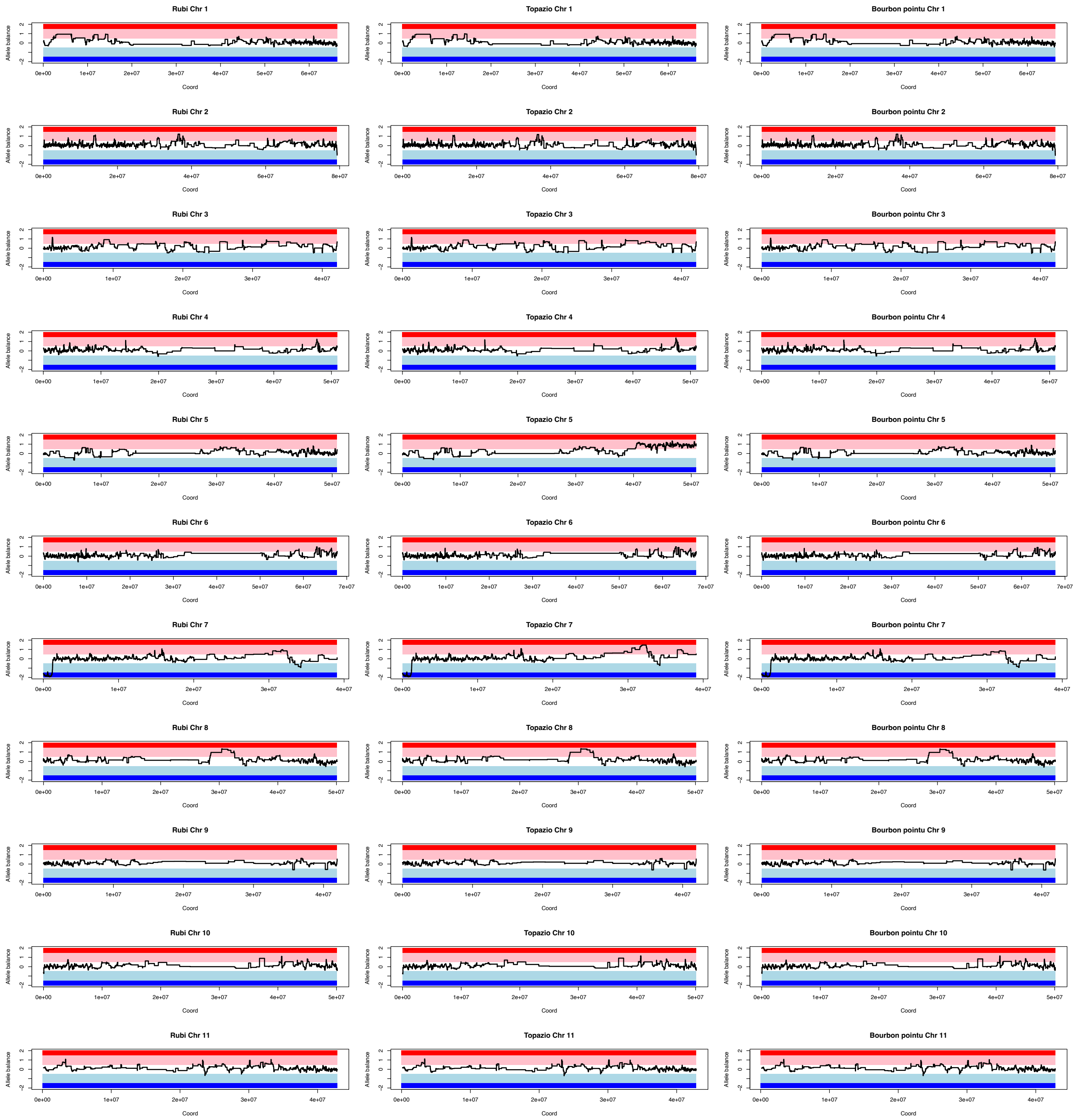

Rubi Chr 1
Topazio Chr 1
Bourbon pointu Chr 1
Rubi Chr 2
Topazio Chr 2
Bourbon pointu Chr 2
Rubi Chr 3
Topazio Chr 3
Bourbon pointu Chr 3
Rubi Chr 4
Topazio Chr 4
Bourbon pointu Chr 4
Rubi Chr 5
Topazio Chr 5
Bourbon pointu Chr 5
Rubi Chr 6
Topazio Chr 6
Bourbon pointu Chr 6
Rubi Chr 7
Topazio Chr 7
Bourbon pointu Chr 7
Rubi Chr 8
Topazio Chr 8
Bourbon pointu Chr 8
Rubi Chr 9
Topazio Chr 9
Bourbon pointu Chr 9
Rubi Chr 10
Topazio Chr 10
Bourbon pointu Chr 10
Rubi Chr 11
Topazio Chr 11
Bourbon pointu Chr 11
Allele balance
Coord

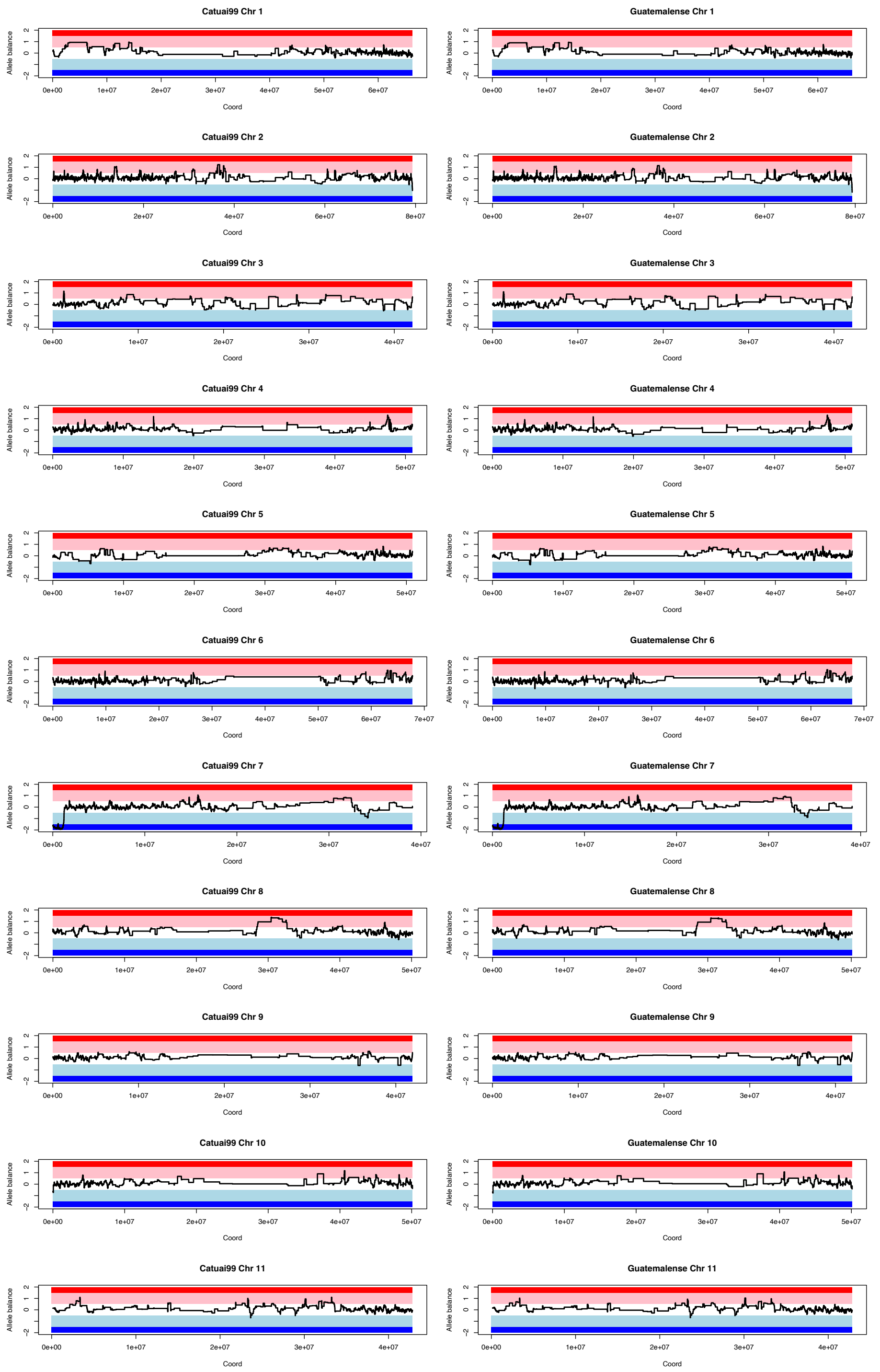


***Fig S35.*** *Homoeologous exchange analysis for cultivated* Coffea arabica *accessions. For each subplot, the title gives the accession ID followed by chromosome number. The dark red region indicates 4:0 allele balance in favor of subCC, while the pink region illustrates 3:1, white 2:2, light blue 1:3 and dark blue 0:4 balances, respectively. The black line indicates the observed allele balances in syntelog gene pairs.*

**Timor hybrids.** Timor hybrid lines are backcrosses of an original spontaneous *Coffea arabica* x *C. canephora* hybrid to cultivated Arabicas (Bourbon or Typica lines). Therefore, we expected to see larger amounts of homoelogous exchange the closer to the original hybrid the lines were. The introgression was most pronounced in Timor hybrid chromosome 6, where one chromosome pair appeared to still originate from *C. canephora*, resulting in a chromosome-wide 3:1 ratio (**Fig S36**).

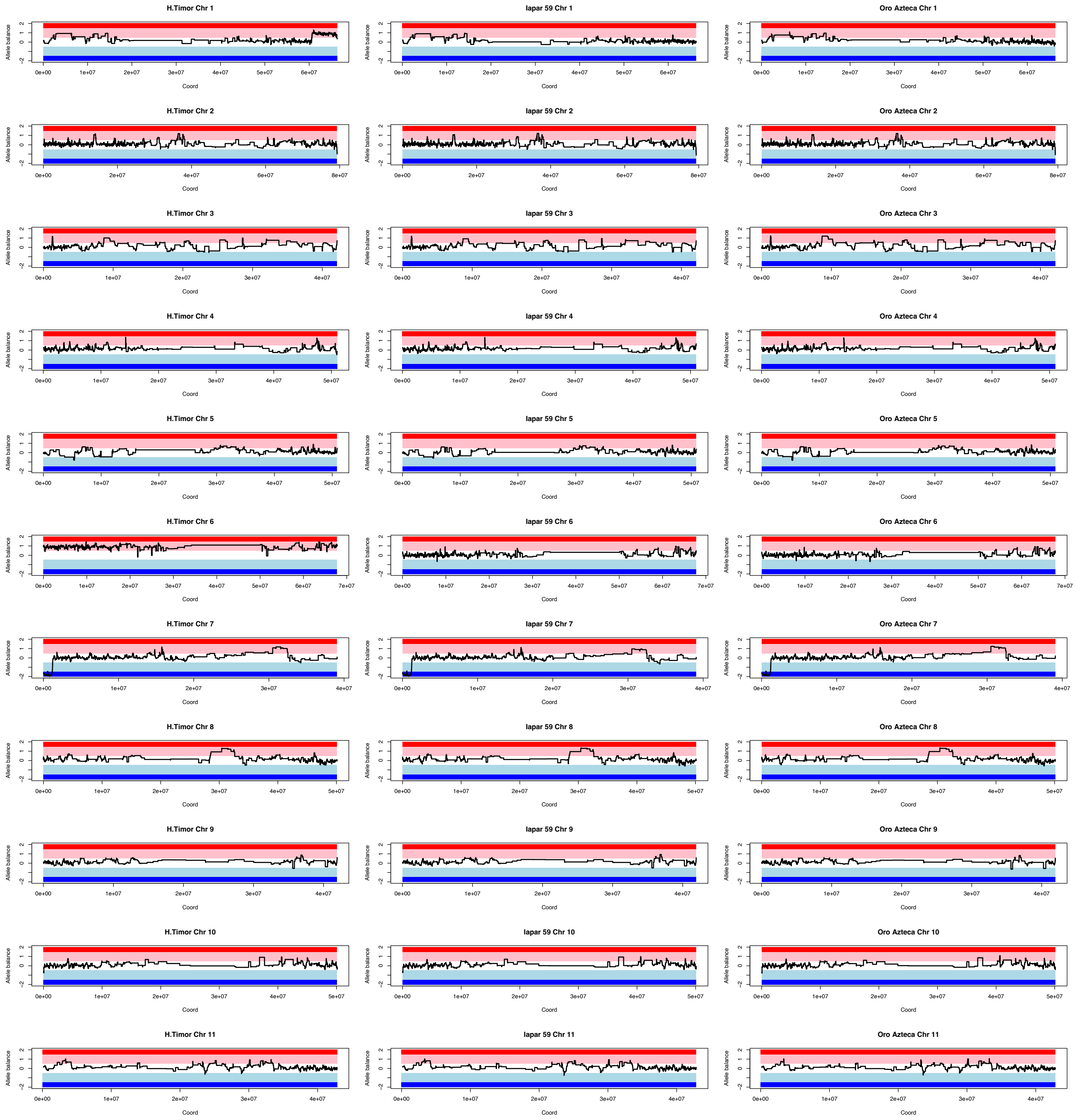
H.Timor Chr 1
Iapar 59 Chr 1
Oro Azteca Chr 1
H.Timor Chr 2
Iapar 59 Chr 2
Oro Azteca Chr 2
H.Timor Chr 3
Iapar 59 Chr 3
Oro Azteca Chr 3
H.Timor Chr 4
Iapar 59 Chr 4
Oro Azteca Chr 4
H.Timor Chr 5
Iapar 59 Chr 5
Oro Azteca Chr 5
H.Timor Chr 6
Iapar 59 Chr 6
Oro Azteca Chr 6
H.Timor Chr 7
Iapar 59 Chr 7
Oro Azteca Chr 7
H.Timor Chr 8
Iapar 59 Chr 8
Oro Azteca Chr 8
H.Timor Chr 9
Iapar 59 Chr 9
Oro Azteca Chr 9
H.Timor Chr 10
Iapar 59 Chr 10
Oro Azteca Chr 10
H.Timor Chr 11
Iapar 59 Chr 11
Oro Azteca Chr 11
Allele balance
Coord

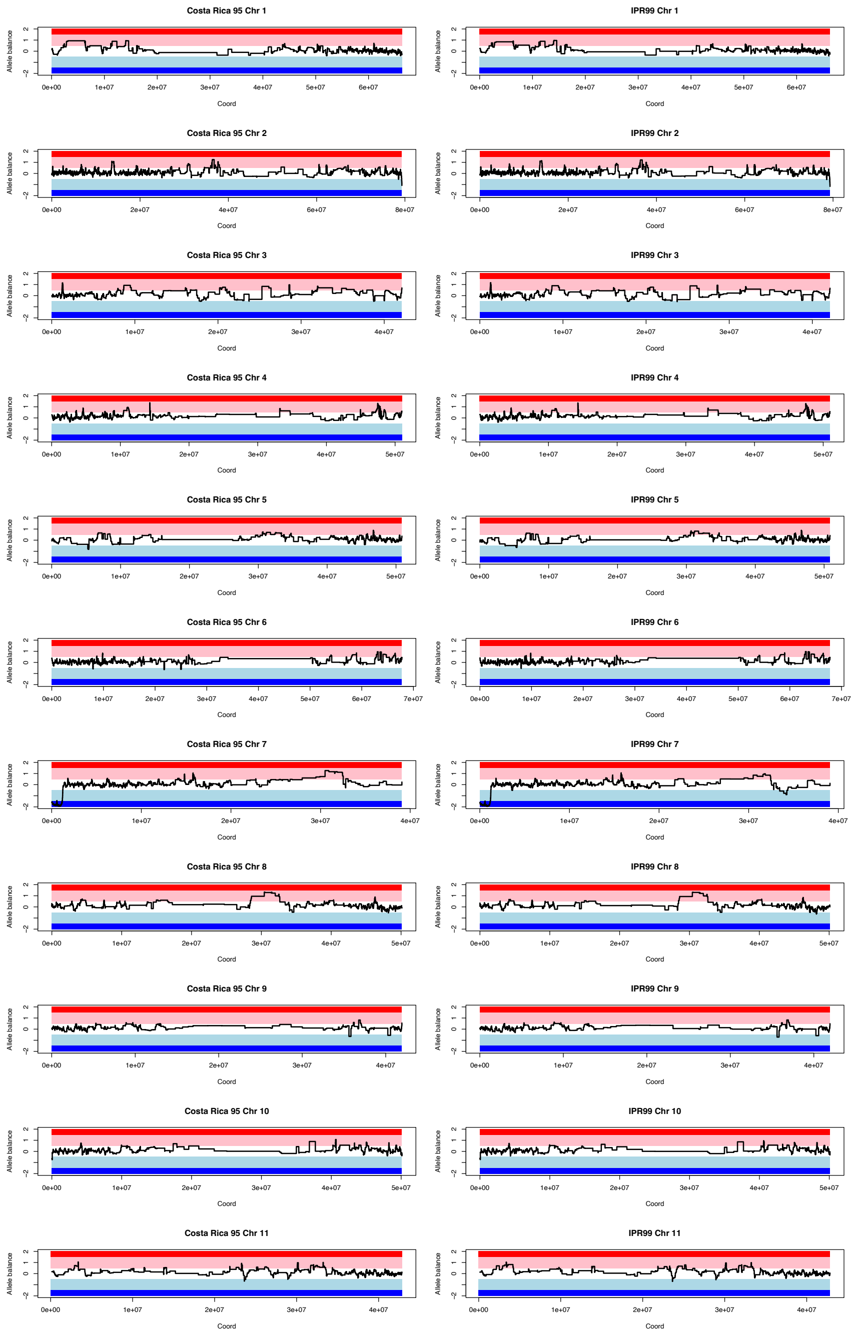


***Fig S36.*** *Homoeologous exchange analysis for Timor hybrid-derived* Coffea arabica *accessions. For each subplot, the title gives the accession ID followed by chromosome number. The dark red region indicates 4:0 allele balance in favor of subCC, while the pink region illustrates=3:1, white 2:2, light blue 1:3 and dark blue 0:4 balances, respectively. The black line indicates the observed allele balances in syntelog gene pairs.*

***Coffea arabica* x *C. canephora* hybrid.** Arabusta pizze is a synthetic hybrid that has been obtained from *C. arabica* crossed with *C. canephora*. As expected, the entire genome showed 3:1 allele ratios (**Fig S37**).

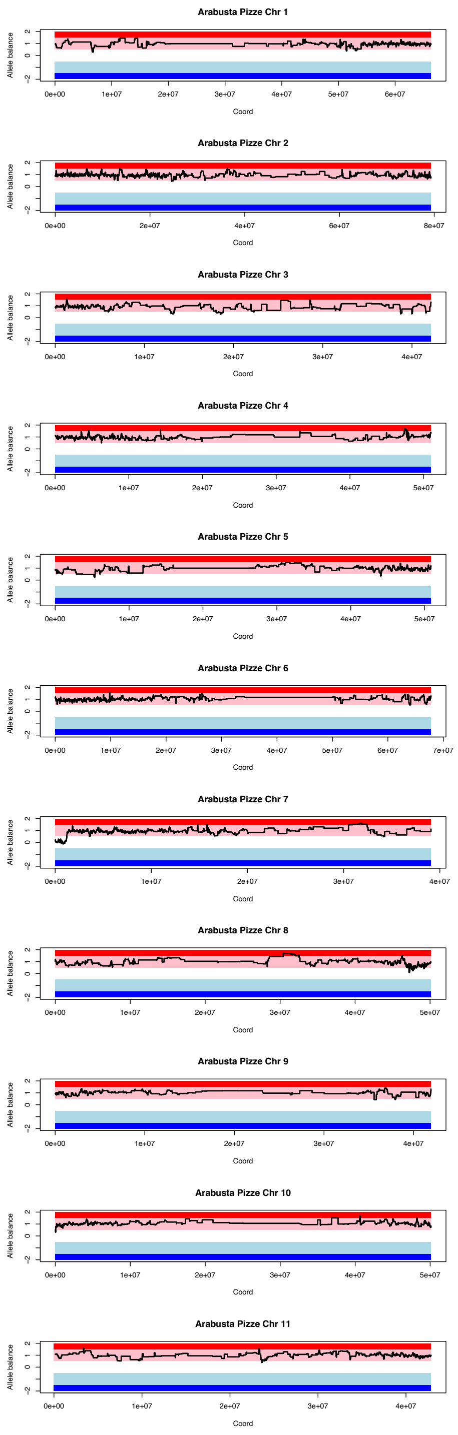


***Fig S37.*** *Homoeologous exchange analysis for Arabusta Pizze, a* Coffea arabica *x* C. canephora *hybrid accession. For each subplot, the title gives the accession ID followed by chromosome number. The dark red region indicates 4:0 allele balance in favor of subCC, while the pink region illustrates=3:1, white 2:2, light blue 1:3 and dark blue 0:4 balances, respectively. The black line indicates the observed allele balances in syntelog gene pairs.*

Since the accessions were closely related and appeared to share the same HE events, we plotted them together in the same plot (**Fig S38**). The regions with similar HE patterns clearly overlapped much more than would be expected from independent HE events, suggesting common ancestry.

Chr 1

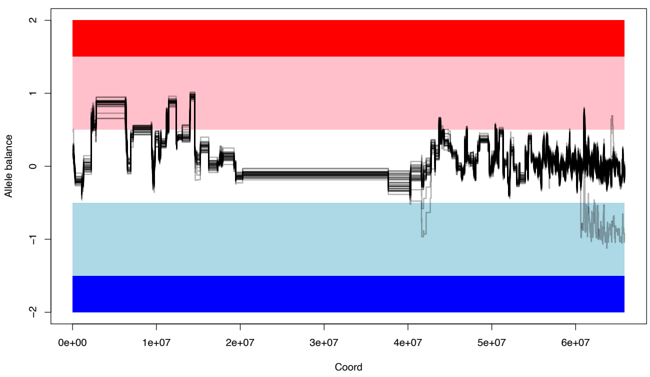


Chr 2

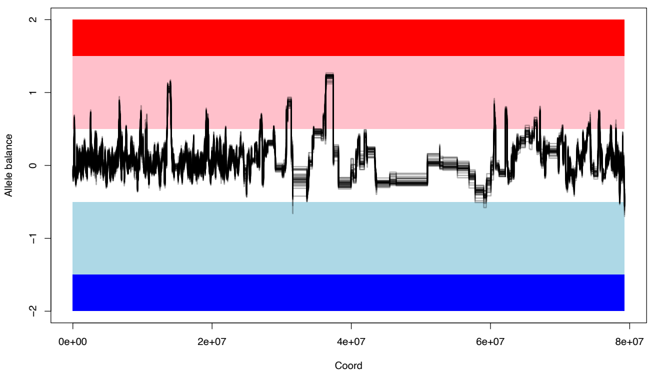


Chr 3

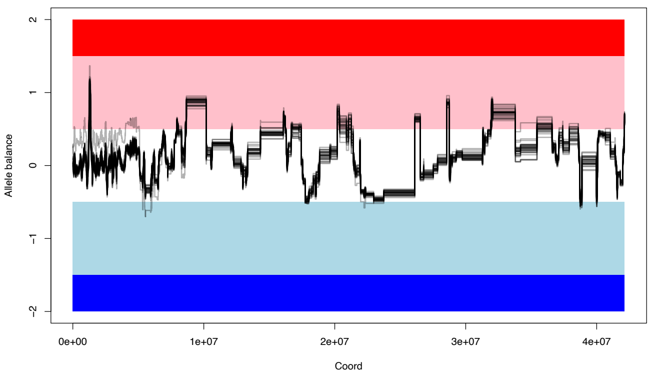


Chr 4

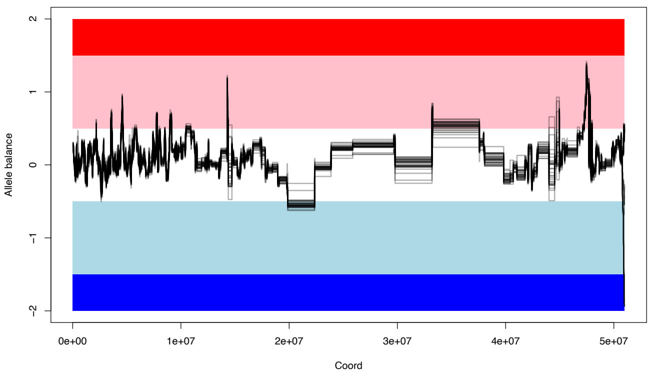

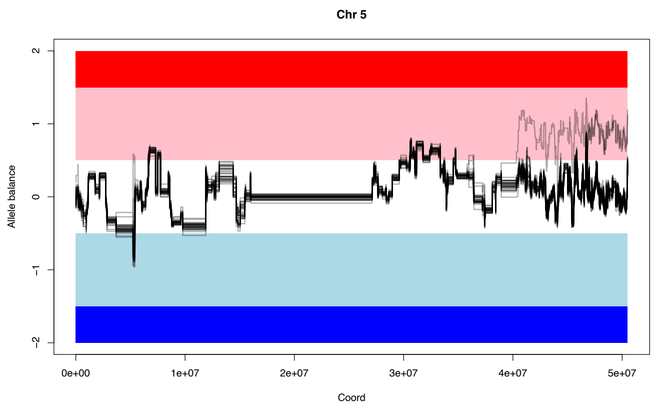
Chr 5
Allele balance
2
1
0
−1
−2
0e+00
1e+07
2e+07
3e+07
4e+07
5e+07
Coord

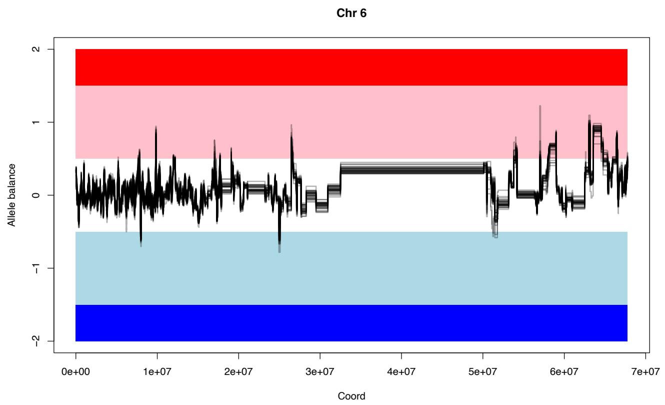
Chr 6
Allele balance
2
1
0
−1
−2
0e+00
1e+07
2e+07
3e+07
4e+07
5e+07
6e+07
7e+07
Coord

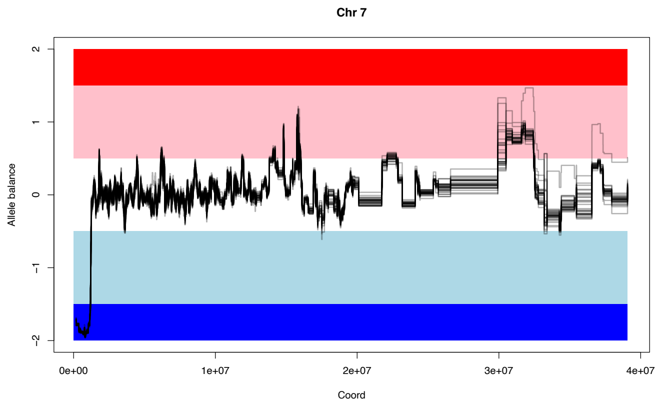
Chr 7
Allele balance
2
1
0
−1
−2
0e+00
1e+07
2e+07
3e+07
4e+07
Coord

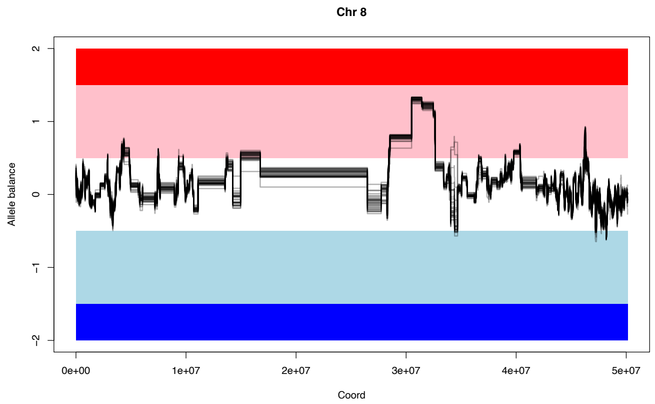
Chr 8
Allele balance
2
1
0
−1
−2
0e+00
1e+07
2e+07
3e+07
4e+07
5e+07
Coord

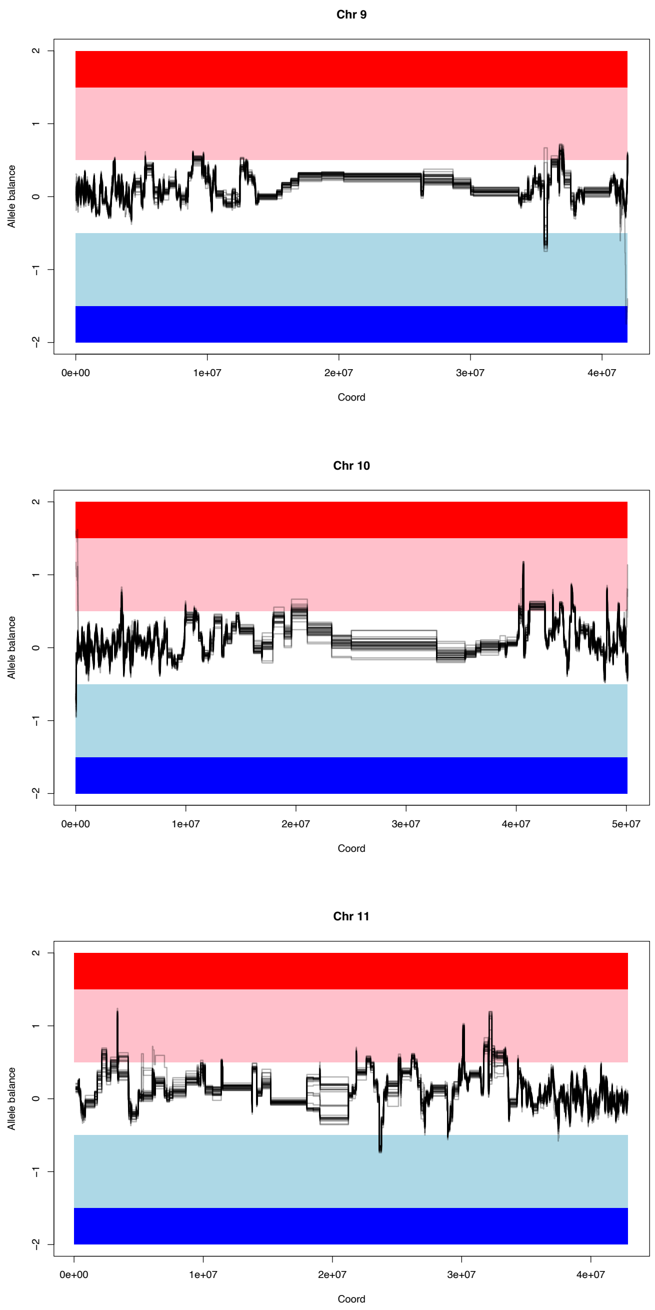


**Figure S38.** *Homoeologous exchange analysis by overlaying all Coffea arabica accessions in this study. Each subplot shows a different homoeologous chromosome. The dark red region indicates 4:0 allele balance in favor of subCC, while the pink region illustrates=3:1, white 2:2, light blue 1:3 and dark blue 0:4 balances, respectively. The gray lines indicate the observed allele balances in syntelog gene pairs for the different Arabica accessions (Individual plots are shown above in Figs S33-S36). For a detailed view of the genes involved, see* ***Table S15****.*

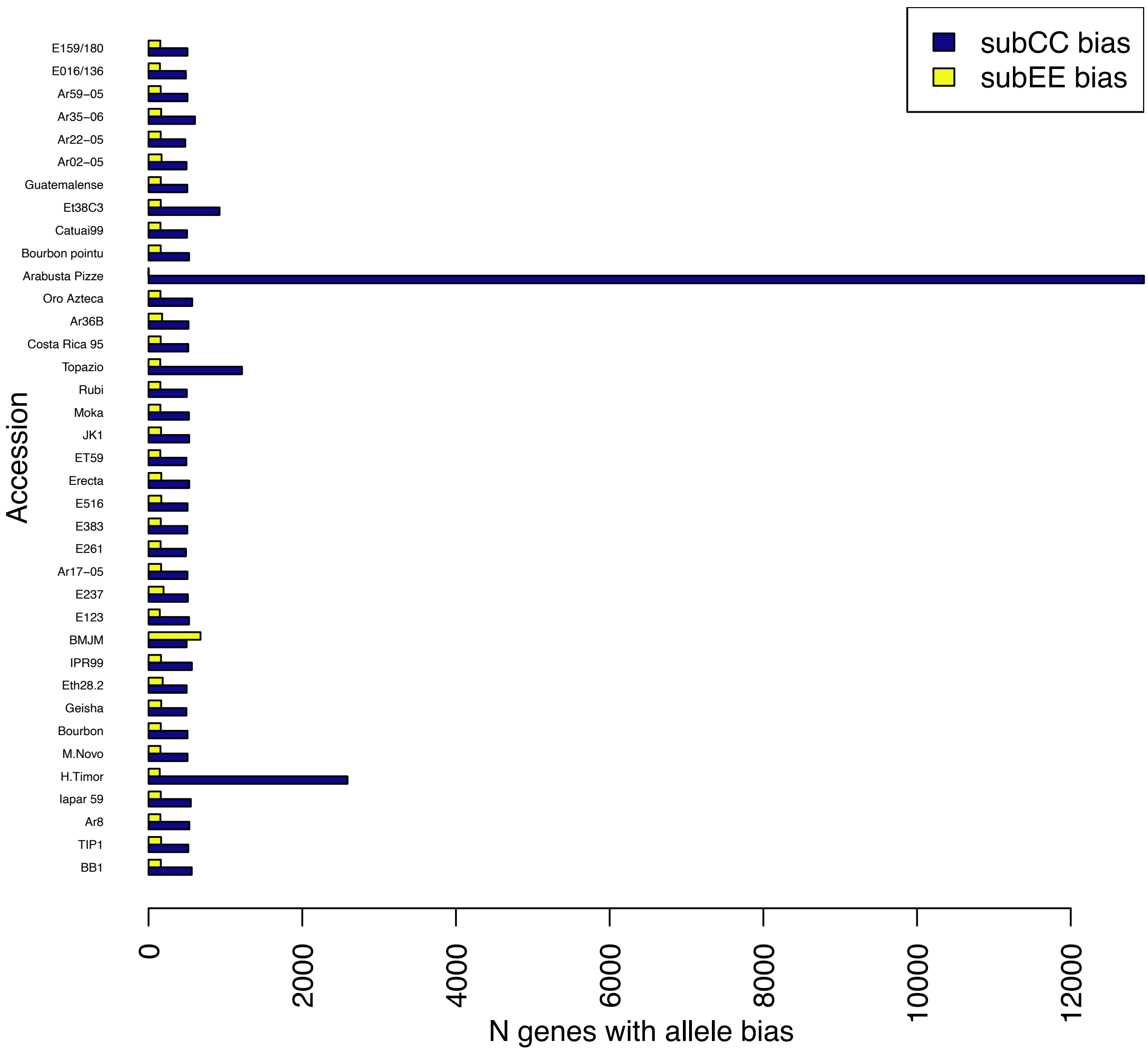


***Fig S39.*** *A summary of homoeologous exchange between subgenomes. The blue bar indicates genes with 3:1 allele bias towards subCC, whereas the yellow bar indicates genes with allele bias (1:3 or 0:4) towards subEE.*

To assess the overall bias between homoeologous exchange events between the subgenomes we next collected the genome information for each of the accessions (**Fig S39**). Most of the HE events resulted in 3:1 allele ratios, and only one event with full 4:0 bias towards subEE was observed. We tested functional enrichment of this region using GOATOOLS (Klopfenstein et al., 2018), and identified enrichment for genes associated with the chloroplast (**Table S9**). Since *C. eugenioides* is the maternal parent of *C. arabica*, this HE is most likely due to organellar compatibility. All but one of the accessions showed a significant bias towards subCC (p-values <<9.8e-37; Chi-square test); BMJM had a significant bias towards subEE (p<1.35e-05; Chi-square test).

We also calculated the frequency spectrum of the HE events (**Fig S40**). There were many events that were common among all accessions, as well as events that were present in only a few individuals, reflecting their recent occurrence (see also **Fig S38** above). The U-shape suggests strong selection for the HE events, such that they were rapidly fixed or purged from the genome, resulting in low numbers of HE events with intermediate frequencies.

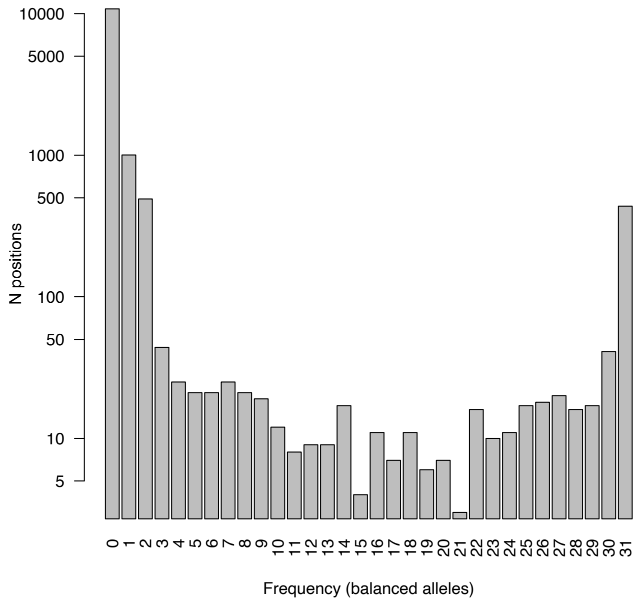


***Fig S40.*** *Site frequency spectrum of alleles with shared homoeologous exchange.*

### 6 Population data analysis

#### 6.1 Sample collection and characteristics

For this study, 38 *Coffea arabica* genotypes including 15 cultivated, 17 wild-type, and six introgressed *C. arabica* plants, as well as 3 *C. canephora* and 2 *C. eugenioides* accessions were sequenced with high ~40x coverage (**Table S8**). Plant material was provided by Embrapa Cerrados (Planaltina, DF) - Brazil; IAPAR institute (Londrina, Parana) - Brazil; R & D Nestlé -Ecuador and France; IRD - France; and Hortus Botanicus Amsterdam - The Netherlands (Sup. Excel Sheet Accessions DB). Wild-type plants were originally collected in the West and East side of the Great Rift Valley in Ethiopia from the FAO (Meyer et al., 1968) and ORSTROM (Halle, 1978) missions in the 1960s. For the FAO mission, seeds from original plant collections from CATIE (Costa Rica) were sent to IAPAR Paraná (via Agronomic Institute of Campinas-IAC Campinas) and also to Embrapa Cerrados. Some accessions were also collected in different germplasm banks of these institutions and provided in the frameworks of several research projects. This was, for example, the case for the accessions BMJM, ET59 and JK1 furnished by Embrapa but originally provided by IRAD (Cameroon) and CIRAD in the framework of the "Coffee genetic diversity in relation to drought tolerance (COFDROnet)" research project supported by Embrapa MarketPlace Brazil/Africa (n°2011.3.415) and Agropolis Fondation-CAPES (n° 1102-002).

Historically, *C. arabica* dispersion around the world relied on two main varieties: Typica and Bourbon. Variety Typica was introduced in the northern Brazil region (Pará State) in 1727, coming from seeds from French Guyana. *C. arabica* var. Bourbon was introduced in 1859, from Réunion Island. At the Instituto Agronômico de Campinas (IAC), both were introduced in 1887, in studies related especially to plant productivity and nutrition. In 1932, coffee trees of both varieties were introduced into the *Coffea* germplasm bank maintained, *ex situ*, by the institute. These Typica and Bourbon varieties have been used for more than 50 years as the main references for genetic analyzes, and for clarifying the origin of mutants and variations of different nature in *C. arabica* (Filho & Carvalho, 1957; Krug & Carvalho, 1951).

Introgressed genotypes originated from *C. arabica* crosses with the Timor Hybrid (HT), a spontaneous hybrid between *C. arabica* and *C. canephora*, found in 1927 in Timor Island. The Sarchimors Iapar-59 and IPR99 accessions result from a *C. arabica* Villa Sarchi x HT 832/2 cross, whereas the Caturra x HT 832/1 cross resulted in Catimors Costa Rica 95 and Oro Azteca.

The Amsterdam sample was provided by Amsterdam Botanical Garden, accession 20120026. The Linnaean material is from a herbarium sample, a lectotype for *C. arabica*, provided by the Linnean Society in London; it was probably collected in Yemen in the 18$^{th}$ century (http://linnean-online.org/2490/).

*C. arabica* Bourbon (BB1) and Typica (TIP1) samples were provided by IAC– SP – Brazil.

Backgrounds of cultivated material:

- Mundo Novo - Issued from a natural cross between Typica and Bourbon
- Bourbon Teksic - Typica mutant, Bourbon Variety after mass selection from Guatemala
- BMJM - Blue Mountain (Typica mutant) from Jamaica
- Rubi - Issued from a cross between Catuaí Vermelho and Mundo Novo
- Topazio - Issued from a cross between Catuaí Amarelo and Mundo Novo
- Bourbon pointu – Bourbon mutant from Réunion Island, single locus mutation called "Laurina"
- Catuai99 - Issued from a cross between Mundo Novo and Caturra (Caturra is a mutated Bourbon)
- Guatemalense - Dwarf mutant probably corresponding to "Pache colis" found in Mataquescuintla (Guatemala)
- Geisha - Ethiopian accession, a wild accession directly field-planted but following some breeding cycles
- Erecta - Typica mutant probably from Java , Indonesia
- Moka – A Bourbon dwarf mutant

#### 6.2 Data preprocessing, SNP calling and annotation

Following quality control with FastQC (Andrews, 2010), the reads were trimmed using Trimmomatic v0.36 (Bolger et al., 2014) and mapped on the *Coffea arabica* reference assembly with BWA mem v0.7.16a-r1181 (Li & Durbin, 2009). Subsequently, the genome analysis toolkit (GATK) pipeline (Poplin et al., 2018) was used for SNP calling. First, libraries for the same sample that were run on different lanes were given individual headers with SAMtools *addreplacerg* and the libraries were then merged using SAMtools merge (Li et al., 2009). Subsequently, duplicates were marked and removed using Picard v2.18.14, and genotype likelihoods were called as a GVCF file using HaplotypeCaller in GATK v3.8.0.

For the Linnaean herbarium sample, sequenced to 46x coverage with Ion Torrent technology, the reads were processed according to the protocols recommended for degraded DNA analysis. First, BWA samse (Li &

Durbin, 2009) was used for mapping the Ion Torrent raw reads against the *C. arabica* assembly. Next, MapDamage v. 2.0.8 (Jónsson et al., 2013) with default settings was used for adjusting the alignment qualities, taking into account the specific nature of DNA degradation. Low quality reads with alignment Phred scores <10 where then filtered out using SAMtools view (Li et al., 2009) and the resulting bam alignment file was then incorporated into the GATK SNP calling pipeline, including duplicate removal (MarkDuplicates) and read group addition (AddOrReplaceReadGroups) using Picard v.2.18.14, and calling genotype likelihoods with HaplotypeCaller in GATK v.3.8.0.

For the diploid progenitors, to allow interspecies comparisons, the mapping was done to each of the subgenomes separately, including chromosome zero, i.e., contigs not assembled into pseudomolecules, in both mappings.

Joint calling was carried out using GenotypeGVCFs in the same GATK version. For each subgenome, the Arabica samples were combined with diploid progenitor data mapped against subEE or subCC, respectively, and after joint calling the subgenome-specific calls were extracted into their own VCF files.

To remove regions with cross-species mappings, we additionally mapped the ET39 sequencing data to the Arabica reference genome. Since the accession is a di-haploid the heterozygous positions necessarily result from mapping between the subgenomes. These heterozygous loci were then filtered out from the VCF files during the SNP filtering step. In addition to removing the SNPs, only biallelic SNP loci were selected for further analysis. All individuals with depth smaller or equal to 3 and mapping quality < 10 were removed. Additionally, the loci with Hardy-Weinberg Equilibrium test $p$<1e-07 were filtered out.

To remove regions with low confidence mappings, a higher quality SNP file was also produced; in this case all the SNPs in repeat regions were filtered out using BEDtools (Quinlan & Hall, 2010) and the gff3 file with repeat annotations from Repeat Finder.

SnpEff v4.3t (Cingolani et al., 2012) was used to assess the functional impacts of the SNPs.

##### 6.2.1 Analysis of GBS data

Read data from 736 *Pst*I GBS libraries of *Coffea arabica* were downloaded from the NCBI sequence read archive (SRA BioProject PRJNA554647). The libraries were 100 bp single-end sequenced on an Illumina HiSeq2000 instrument. The 5' restriction site remnant, the 3' restriction site remnant, and the common adapter sequence were removed with Cutadapt v2.10 (Martin, 2011). Trimmed reads with an average base quality score below 25 were excluded using a custom python script, while reads with internal restriction sites were discarded using Cutadapt v2.10. After trimming and quality filtering, reads were mapped onto the *C. arabica* reference genome sequence using the BWA-mem algorithm with all default settings in BWA v0.7.17 (Li & Durbin, 2009). Upon the first mapping of the GBS data, only 50% of the reads could be mapped with a Phred quality score of 20 or higher. Many of the discarded reads mapped (equally) well onto both subgenomes, resulting in a lower quality Phred score. This non-unique mapping of reads onto a polyploid reference genome sequence is especially problematic for short single-end sequencing data (read length < 86 bp). To increase the percentage of mapped reads, a "genome read categorization" approach was applied. First, all reads were mapped onto each separate subgenome of the *C. arabica* reference genome sequence. Afterwards, a read was assigned to one of the subgenomes using a custom python script if: (i) the read could be mapped onto one subgenome but not onto the other; (ii) the read could be mapped onto both subgenomes but its mismatch score with one subgenome was lower than its mismatch score with the other. The mismatch score was calculated as the sum of the number of nucleotide differences, the length of insertions and deletions in the read divided by 2, and the length of hard- and soft-clipped regions in the read divided by 10. Reads that matched equally well with both subgenomes, with a mismatch score higher than 20, or with a mapping score below 20, were discarded. By using this approach, the percentage of mapped reads with a MAPQ of 20 or higher increased on average from 50% to 70%. Although this approach is not perfect, it substantially increased the number of correctly mapped GBS reads.

Mapped reads were sorted and indexed with SAMtools v1.10 (Li et al., 2009), and read groups were added using Picard (Picard_toolkit, 2019). Single Nucleotide Polymorphisms (SNPs) were called using the Unified Genotyper of GATK v3.7 (McKenna et al., 2010), and filtered for a minimum minor allele count of 4 and a minimum SNP quality of 20 using VCFtools v0.1.16 (Danecek et al., 2011). Genotype calls were converted into missing data if their read depth was below 30, their quality score below 30, and their allele depth below 3 with VCFtools and a custom python script. Non-polymorphic and multi-allelic SNPs were discarded as well with GATK v3.7, resulting in a set of 935 SNPs. 313 out of 935 SNPs had a heterozygous genotype call in a minimum of 368 individuals (50%). These SNPs were considered to be fixed polymorphisms between subCC and subEE in *C. arabica*, corresponding to ancestral polymorphisms between the *C. arabica* progenitor species. As the species *C. arabica* likely emerged after a single polyploidization event (Scalabrin et al., 2020), *between*-subgenome polymorphisms were assumed to be present in all *C. arabica* individuals. Such positions are not informative in an assessment of genetic differences among individuals, hence they were excluded from the dataset. Finally, 88 SNPs were removed because they had data in less than 19 out of 736 samples (5%). In total, 534 SNPs (235 SNPs on the subCC, 278 SNPs on the subEE, and 21 SNPs on unanchored contigs) were identified in 3,966 loci with data in a minimum 19 out of 376 samples (5%), representing 345,031 resequenced nucleotides (0.03%) of the 1.1 Gbp *C. arabica* reference genome sequence. The python scripts for the assignment of reads to the *C. arabica* subgenomes and for the additional filtering steps that were applied on the VCF file are available on GitLab (https://gitlab.com/ybawin/sequence-data-processing-tetraploids).

The genotype calls of the 534 SNP positions in 736 GBS samples and the 39 whole genome resequenced *C. arabica* samples were combined into one VCF file for each *C. arabica* subgenome separately (775 samples in total) with VCFtools. A principal component analysis (PCA) was run on those VCF files. The first principal component explained 13.2% and 21.0% of the total variation in the SNP positions in the subCC and subEE, respectively. The second principal component explained 10.0% of the variation in the subCC and 8.6% of the variation in the subEE. The 39 whole genome resequencing samples aligned well to the set of 736 GBS samples, comprising a substantial part of the genetic variation in the GBS dataset (**Fig. S41**).

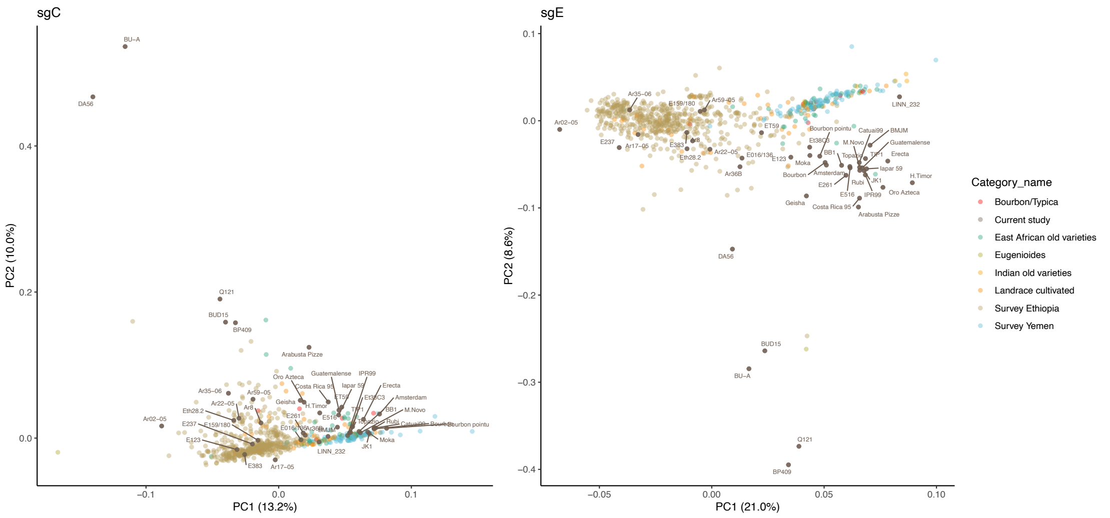


***Figure S41.*** *Scatterplots of the first (PC1) vs the second (PC2) principal component from the PCA based on the genotype calls from 235 SNP positions in subCC (left) and 278 SNP positions in subEE (right) in our 39 whole genome resequencing samples and the 736 GBS samples from Scalabrin et al. (2020). Labels indicate*

*the 39 whole genome resequencing samples, whereas colored dots mark the GBS samples. The GBS samples were partitioned into seven categories: Bourbon/Typica, East African old varieties, Eugenioides, Indian old varieties, Landrace cultivated, Survey Ethiopia, and Survey Yemen.*

#### 6.3 SNP analyses

Genome-wide nucleotide diversity was calculated with VCFtools v.0.1.17, by calculating the mean of π values from sliding windows of 100 kb with 10 kb step size. Similarly, genome-wide Tajima's D was calculated from the mean of Tajima's D values with window size of 10 kb (**Table S10)**. The inbreeding coefficient was calculated with VCFtools (**Fig. S42**). The Linnaean sample stood out with an especially high level of homozygosity, but this was likely due to low sequencing coverage obtained from the centuries-old herbarium sample (~1.5x).

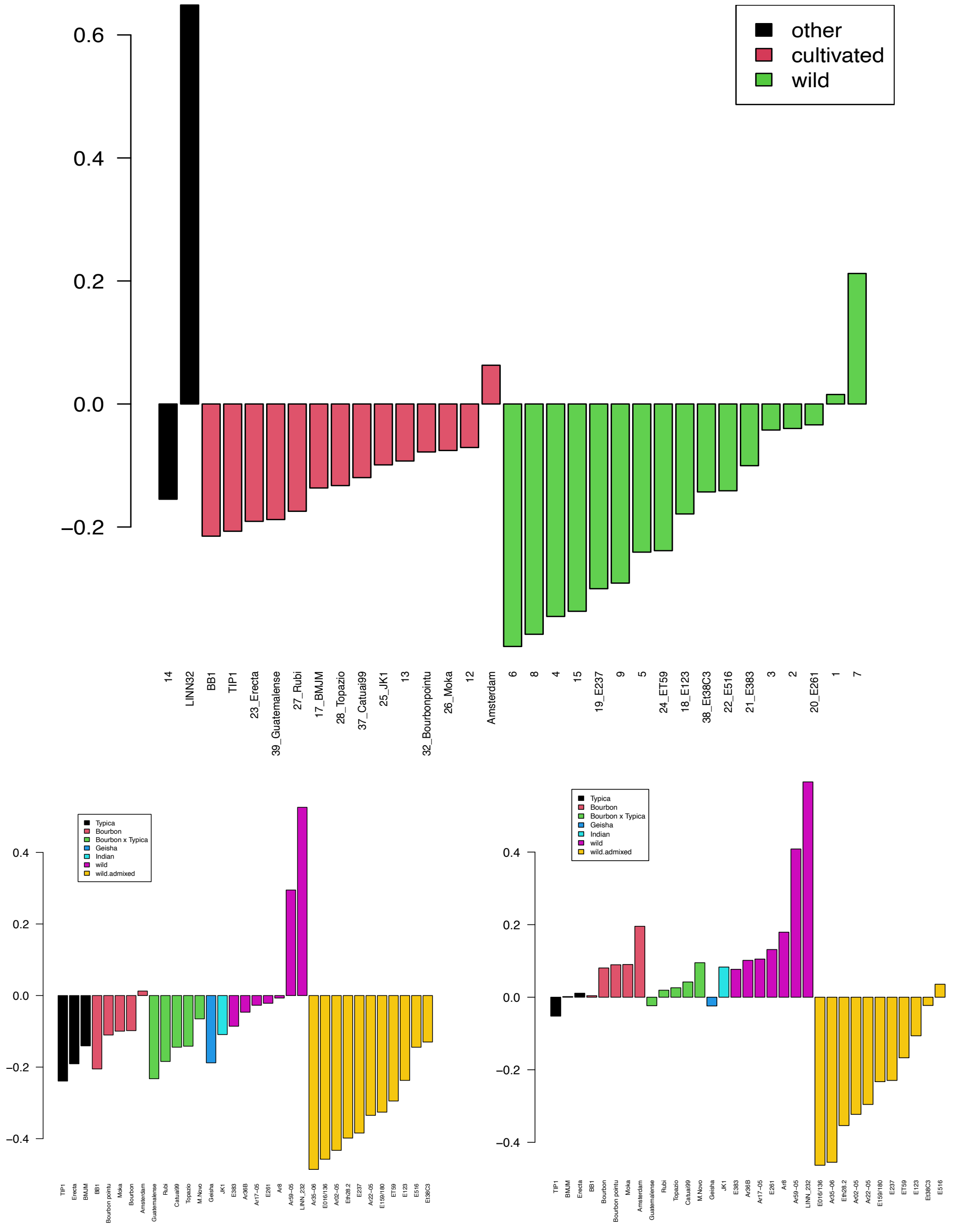


***Fig. S42.*** *F inbreeding coefficient for wild and cultivated lines.*

Nonsynonymous nucleotide diversity, $\pi_0$, and neutral, integenic $\pi_s$, were calculated using the PiNSiR R package (https://github.com/jsalojar/PiNSiR) and ANGSD v.0.933 (Korneliussen et al., 2014). First, ANGSD was used to calculate site-wise diversities following the manual at http://www.popgen.dk/angsd/index.php/Thetas,Tajima,Neutrality_tests. Then the PiNSiR R package was used to identify neutral and deleterious positions. Briefly, the software first identifies high quality gene models, by searching for gene models with a methionine as start codon and a coding sequence length divisible by three. From these high-quality models, it identifies the positions with amino acid changing mutations, the first and second codon positions, and estimates the nonsynonymous nucleotide diversity as mean of site-wise estimates calculated using ANGSD in a parallel manner.

Neutral diversity is estimated from intergenic positions using the same package by excluding all regions with gene predictions.

A domestication bottleneck usually results in highly reduced genetic diversity in cultivars compared to wild individuals. Accordingly, in Arabica, the mean nucleotide diversity in neutral sites ($\pi_s$) was lower among cultivars and older cultivars with fewer breeding cycles, while Geisha demonstrated higher diversity (**Fig. S43**). From the major cultivar groups, Typica had higher diversity than Bourbon, whereas modern cultivar crosses between the two lines showed intermediate values. The differences in $\pi_s$ could result from the known single-individual bottleneck in Bourbon. Overall, subEE showed higher neutral diversity, suggesting higher original effective population sizes in the CE diploid progenitor. Long-continued inbreeding eventually leads to purging of deleterious alleles, and this can be measured, for example, by the ratio of non-synonymous diversity to synonymous diversity. This ratio was exceptionally high in the Linnaean sample, possibly due to a high level of inbreeding.

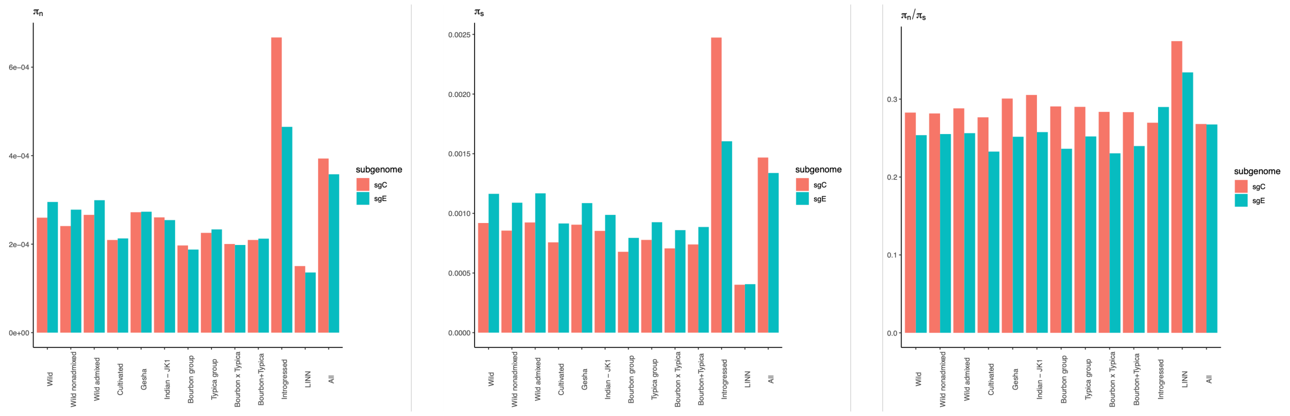


**Fig S43**. *Non-synonymous ($\pi_n$; left) and synonymous ($\pi_s$; middle) diversity between the different population groupings in* Coffea arabica*. The ratio of synonymous vs non-synonymous diversity ($\pi_n/\pi_s$) is illustrated on the right.*

##### 6.3.1 Population structure

Principal component analysis was run using Plink v1.90 (Chang et al., 2015), separately for both subgenomes (**Fig. S44**)

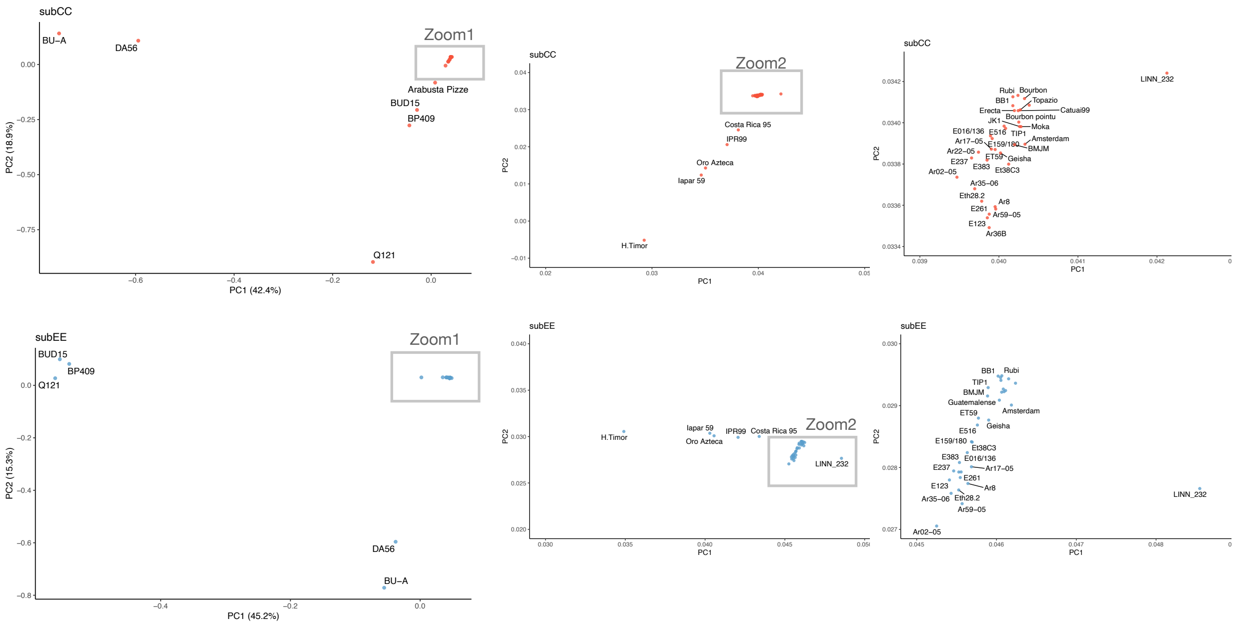


***Fig. S44.*** *PCA plot of the SNPs called in subCC (top) and subEE (bottom) subgenomes. The rectangles highlight the region zoomed in at the plots towards the right of the initial plots.*

ADMIXTURE software (Alexander & Lange, 2011) was run for SNP data where the variants in repeat regions were filtered out and the outgroup species (=diploid *Coffea* species) were excluded. The sites were filtered for linkage disequilibrium in plink v1.90 according to the recommendation in the ADMIXTURE manual with (--indep-pairwise 50 10 0.1), while allowing max. 10% missing values (--geno 0.1). ADMIXTURE analysis was run using 10-fold cross-validation, and software version 1.3.0. The solution giving lowest cross-validation score was selected as the best solution.

##### 6.3.2 SNP trees

The SNPs were filtered for repetitive regions using BEDtools, followed by filtering for LD > 0.4 and loci with > 40% missing values, as well as minor allele prevalence <10%. The fasta file obtained from the selected sites was then input into RAxML (Stamatakis, 2014) with -T 30 -m GTRGAMMA model, using 30 starting trees and 1000 bootstrap samples. Trees were calibrated using the chronos function in the R package ape, using a discrete clock model and by setting the divergence between *C. canephora* and *C. eugenioides* to vary between 4-7 Mya (**Fig. S45**).

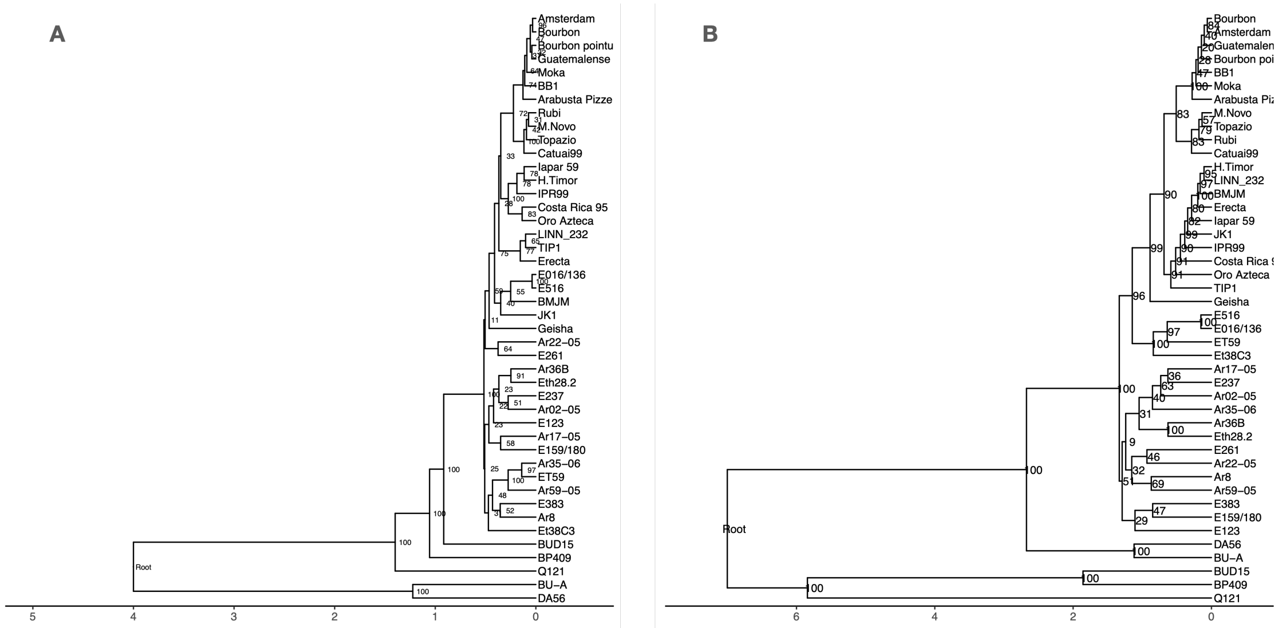


***Fig. S45.*** *Divergence time estimates for the* C. canephora*,* C. eugenioides *and* C. arabica *accessions when analyzed using SNPs from subCC (A) and subEE (B). X-axis is in millions of years.*

#### 6.4 Demography

Since our analysis did not identify pure wild representatives among the modern cultivated population, the precise place of origin for the latter remains unknown. However, an extended period of migration between the two populations is most parsimonious if they were separated only by a relatively small geographic distance, such as along the two sides of the African Great Rift Valley (**Fig. 3C**). It is possible that the second ancestral population could have extended as far as Yemen, ~1,000 km away, and in that case, the end of migration between the two populations may have coincided with the end of the AHP and widening of the Bab al-Mandab strait (separating Yemen and Africa) to tens of kilometers due to rising sea levels (Lambeck et al., 2011). A wild native *C. arabica* population has been identified in Yemen (Montagnon et al., 2021), which could support this hypothesis. The Linnaean sample, together with the Typica and Bourbon cultivars, originate from this second population that was also used to establish cultivation in Yemen, as suggested by the ADMIXTURE, PCA, and SNP analyses (**Fig. 3B**, **S44, S45**). More recently, when inter-lineage migration ended, both wild and cultivar populations underwent strong independent bottlenecks at ~1 kya; this was observed when analyzing the SMC++ curves for subgroups of wild, wild admixed, Typica and Bourbon individuals (**Figs. S48-S50**). All trajectories showed diverging behaviors between the subgenomes at ~8 kya, roughly the time at which migration was modeled to end. These differences may be due to a limited number of migrating individuals with different histories; in contrast, the wild, non-admixed individuals showed a strong bottleneck and overall low effective population sizes for both subgenomes.

##### 6.4.1 Pairwise sequentially Markovian coalescent (PSMC)

For each individual, the reads were mapped against the full reference assembly. Then the mappings were filtered for indels using BCFtools (Danecek et al., 2021) and regions with <8x or >100x coverage. After filtering, the obtained fastq file was split into subEE and subCC specific parts, and PSMC (Li & Durbin, 2011) demography was estimated using standard parameter settings (-N25 -t15 -r5). The inferred history was then visualized using R and the ggplot2 package (**Fig. S46**).

We have recently demonstrated that widespread selfing within a population biases demographic trajectories estimated from sequencing data (Hu et al., 2022). Theory predicts (Nordborg & Donnelly, 1997) that the loss of heterozygosity results in faster coalescence, and this would be visible from population trajectories by a "shift" towards modern times. In the case of CA, the pure wild individuals showed higher levels of inbreeding than the admixed wild individuals. This effect was clearly visible in PSMC plots of the wild individuals, where the more heterozygous individuals demonstrated a shift towards more ancestral coalescence (**Fig. S47**). The higher level of heterozygosity in coffee cultivars is due to selective breeding, where lineages are preferentially propagated by crossing of two parents, whereas in the wild, CA preferentially propagates by selfing.

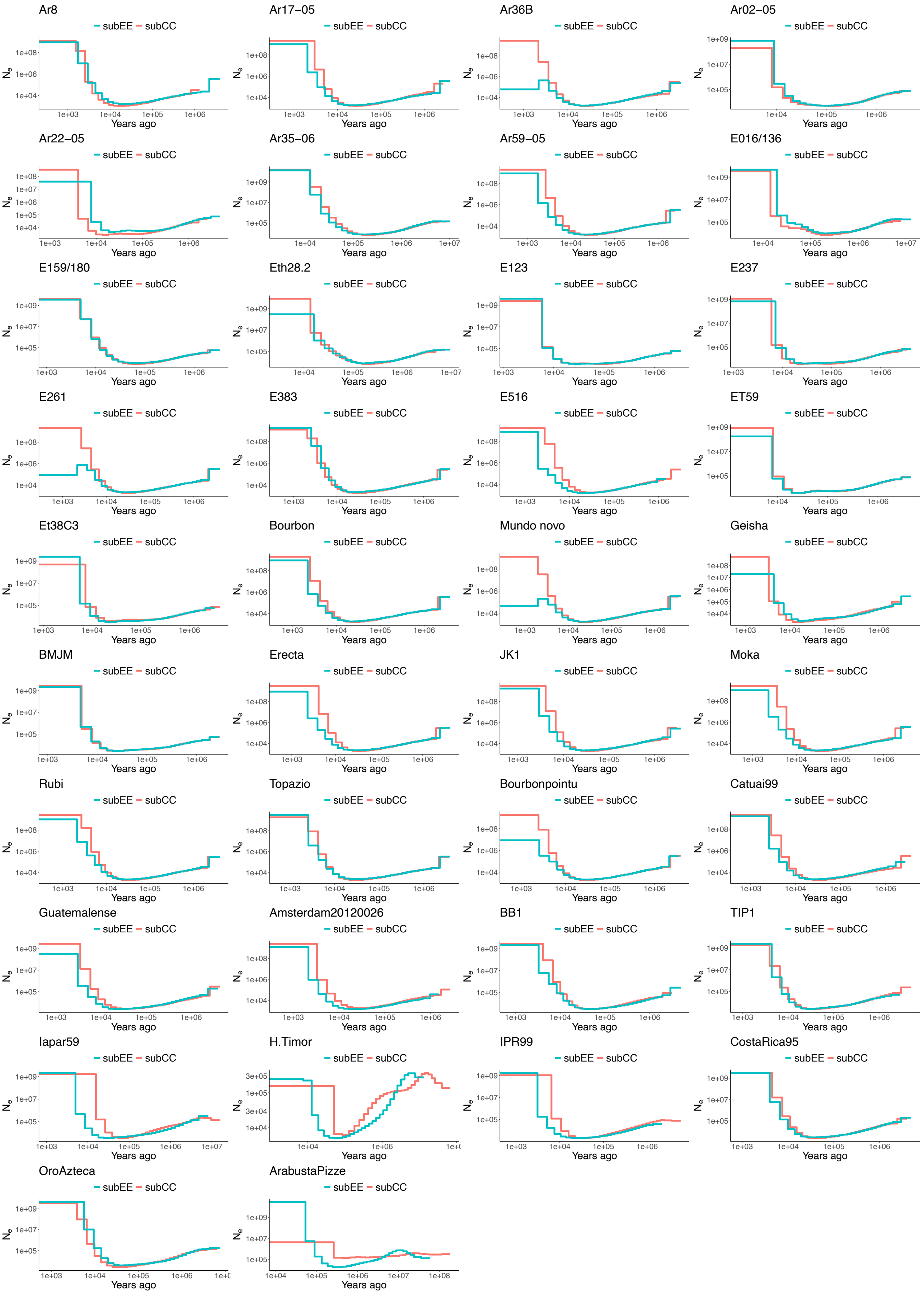
Ar8
Ar17–05
Ar36B
Ar02–05
Ar22–05
Ar35–06
Ar59–05
E016/136
E159/180
Eth28.2
E123
E237
E261
E383
E516
ET59
Et38C3
Bourbon
Mundo novo
Geisha
BMJM
Erecta
JK1
Moka
Rubi
Topazio
Bourbonpointu
Catuai99
Guatemalense
Amsterdam20120026
BB1
TIP1
Iapar59
H.Timor
IPR99
CostaRica95
OroAzteca
ArabustaPizze
subEE subCC
$N_e$
Years ago

***Fig. S46.*** *Pairwise sequentially Markovian coalescent (PSMC) plots of population demography for each of the Arabica accessions. Red illustrates population history inferred from the subCC genome, cyan color the demography estimated from subEE.*

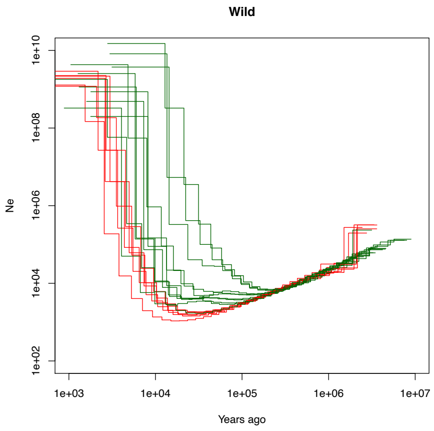


***Fig. S47.*** *Population histories estimated with wild, non-admixed individuals (red) vs. wild, admixed individuals (green). Admixture was determined with ADMIXTURE (see Fig 4), and it shows a shift towards more ancestral coalescence when PSMC was estimated for the subCC genome.*

##### 6.4.2 Ancestral state estimation

The ancestral state was inferred from reads of two representatives of each of the diploid *Coffea* species *C. canephora* (BUD15, Q121) and *C. eugenioides* (BU-A, DA56) mapped against each of the subgenomes and their unassigned contigs. Subsequently, a majority vote was carried out to infer the ancestral allele using ANGSD v.0.933 with options -doFasta 2 and -doCounts 1. The SNP calls in the VCF file were then flipped to the ancestral states using BCFtools +fixref.

##### 6.4.3 SMC++

To identify the demographic events that shaped overall Arabica evolution, we modeled the population history of *C. arabica* accessions using the SMC++ (Terhorst et al., 2017) and pairwise sequentially Markovian coalescent (Li & Durbin, 2011) models (**Fig. 3D**; **Figs. S46-S50**). To avoid possible confounding effects of admixture, the analyses initially focused on non-admixed wild individuals. Both subgenomes concordantly showed two bottlenecks during their paleohistories (**Fig. 3D**). Using a mutation rate of $7.77 \times 10^{-9}$ / (bp*generation)(Xie et al., 2016) and a generation time of 21 years (Moat et al., 2019), the most recent bottleneck initiated ~5,000 years ago (5 kya), and an additional, longer period of lower population size was modeled between 20-100 kya (**Fig. 3D**). Because a large part of modern human evolution occurred in East Africa, its past geoclimatic history is well known, with evidence for an extended drought and cooler climate at 40-70 kya, coinciding with human migration out from Africa (Lambeck et al., 2011) (**Fig. 3E**). During the African humid period (AHP), around 6-15 kya (Kuper & Kröpelin, 2006), growth conditions were likely more beneficial for CA, and this period may coincide with the population increases modelled in SMC++ for both wild and cultivar populations (**Fig. 3D**). While modeling can provide accurate estimates of demographic changes, their timings should be treated with caution since considerable uncertainty exists in generation time(Salojärvi et al., 2019) and mutation rate estimates in plants, including factors contributing to it(Bashir et al., 2014). CA generation time has been estimated (Moat et al., 2019), but the precise mutation rate is not known.

The input data for SMC++ (Terhorst et al., 2017) comprised of the VCF file where the ancestral state was used as reference (see above), and the SNPs in repeat regions were filtered out. For the cultivar population, the representatives of Bourbon and Typica lineages were included (TIP1, Bourbon, Mundo Novo, BMJM, Moka, Rubi, Topazio, Bourbonpointu, Catuai99, BB1, Erecta, JK1, Guatemalense, Amsterdam); Geisha was removed from the analysis because of its unknown pedigree. SMC++ parameter selection was carried out using 3-fold cross-validation (smc++ cv).

In SMC++, the pure, wild, non-admixed individuals showed a strong bottleneck from ~5kya, whereas in admixed individuals the drop was less severe, as expected in admixed individuals with higher overall heterozygosity. Interestingly, in the wild admixed samples the trajectories from subEE and subCC data differed from ~10kya towards modern times (**Figs. S48-S49**). This could result for example from drift, for example in a case where a very small founder population, such as new migrants, hybridizes with a larger wild population.

Interestingly, the SMC++ analysis for introgressed cultivars did not show similar coalescence but rather a steady decline in ancestral population size at this time horizon (**Fig. S48**). Widespread inbreeding is known to accelerate coalescence(Nordborg & Donnelly, 1997), and we have recently shown this to affect demographic modeling as well(Hu et al., 2022). Admixture and introgression, in contrast, introduce intra- and interspecific polymorphisms into genomes. Reflecting this, the population histories of admixed Arabica individuals demonstrate shifts towards more ancient coalescence times (**Fig. S47**), and in the case of Timor hybrids, introgression may have resulted in ancestral polymorphisms introduced back into CA through the hybridization event, resulting in alleles with deep coalescence.

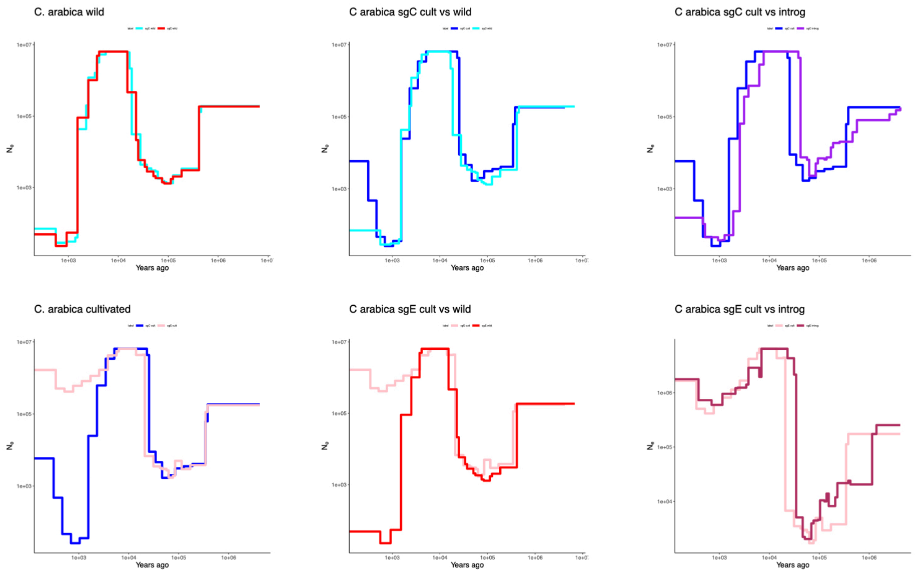


***Fig. S48.*** *Population history of* Coffea arabica *estimated with SMC++. Left: population history from wild and cultivated accessions. Middle: comparison of population histories in wild vs cultivated individuals. Right: Comparison of population trajectories from introgressed individuals vs. cultivars. The different colors in the plots illustrate demographies estimated from different subgenomes (subCC vs subEE).*

**A** **B**

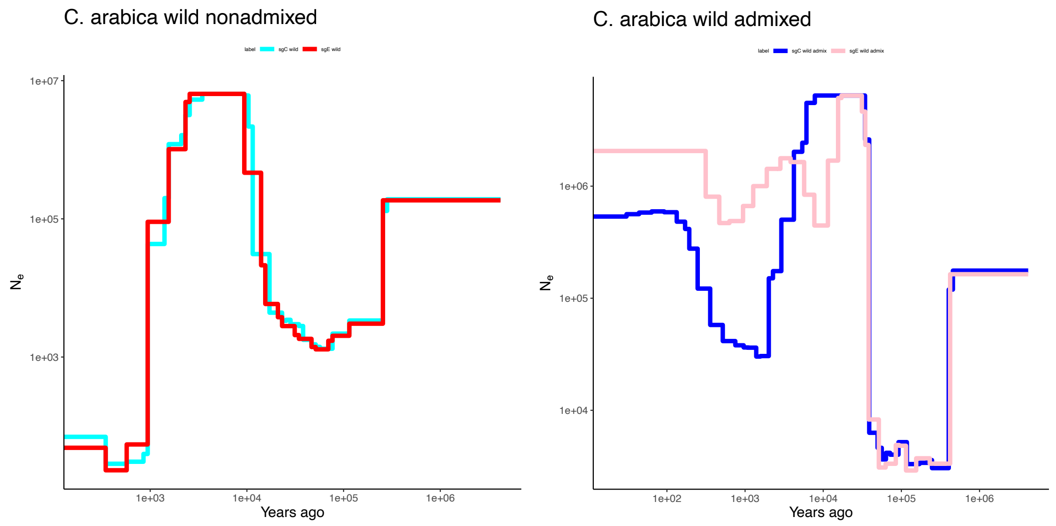


***Fig. S49.*** *Population histories estimated for* Coffea arabica *using SMC++, for non-admixed wild individuals (left) and admixed wild individuals (right). Admixture was determined with ADMIXTURE (see Fig 3). The two trajectories show the histories estimated from subCC or subEE.*

Strikingly, the cultivar groups did not show a strong population decline in subCC (**Fig. S50**). This effect may be due to crossing of genetically diverged populations for breeding purposes.

Interestingly, SMC++ for introgressed cultivars showed ancestral population size changes beyond 370kya; this could be due to reintroduction of ancestral polymorphisms into Arabica through the hybridization event.

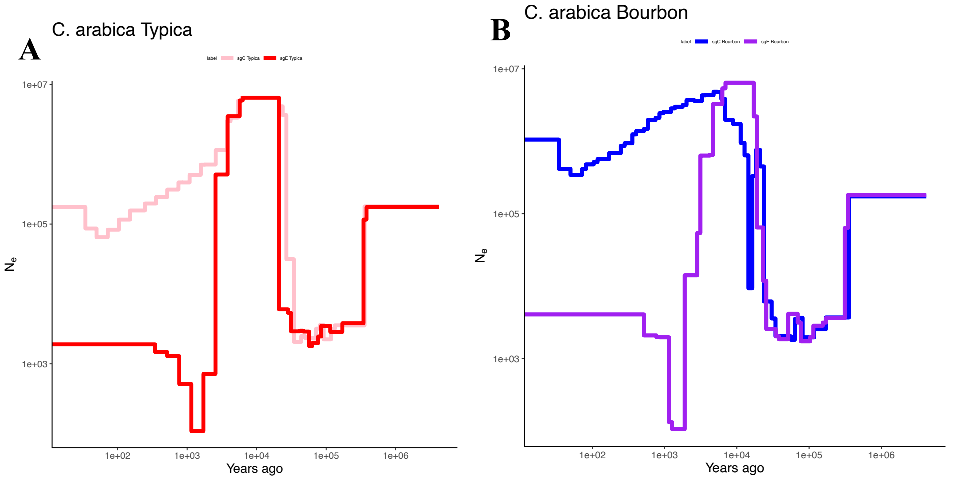


***Figure S50.*** *Population histories estimated using SMC++.* ***A****: Typica group* ***B****: Bourbon cultivars.*

##### 6.4.4 Kinship analysis

Prior to the analysis, the diploid species were removed from the SNP file, and kinship was estimated using KING software v2.2.5. (Manichaikul et al., 2010) with the --kinship option. The results were visualized using Keynote, for each subgenome separately (**Fig S51 – Fig. 4**).

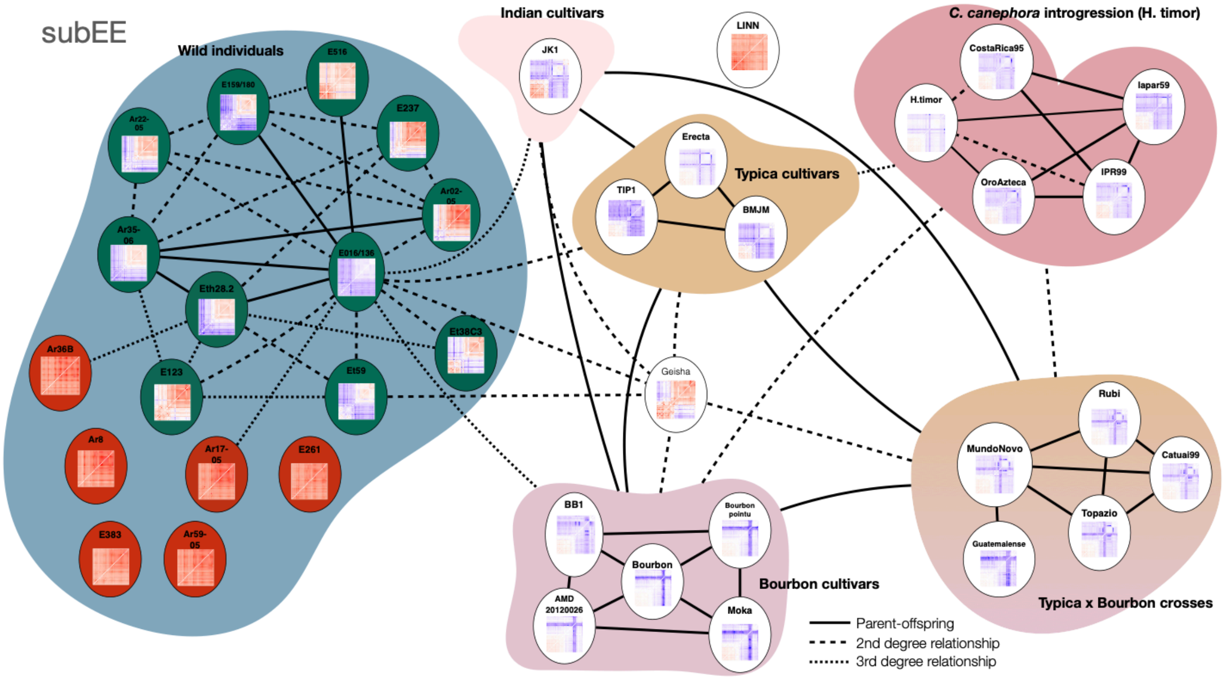


***Figure S51.*** *Kinship analysis on subEE. The degree of relatedness was estimated using Kinship-based INference for GWAS (KING); Thumbnail images show false discovery rate corrected F3 tests of introgression Z-statistics for each of the target individuals; purple-blue color illustrates significant gene flow from the two sources (horizontal and vertical) to the target, or close infrapopulational relationship between sources and target. The green background in the wild accessions highlights the admixed individuals (Figure 3B); the non-admixed individuals are highlighted with red. The corresponding analysis on subCC is shown in* ***Fig. 4.***

##### 6.4.5 F3 statistics

The analysis of introgression was carried out using the SNP data filtered for linkage disequilibrium and repeat regions. The Admixtools package (Patterson et al., 2012) (https://github.com/DReichLab/AdmixTools) was used to calculate the F3 statistics, and the obtained *p*-values were subjected to false discovery rate (FDR) correction using the procedure developed in Salojarvi et al., 2017 (Salojärvi et al., 2017), where the Z-scores were converted into *p*-values, subjected to FDR correction using the Benjamini-Hochberg method, and then converted back to Z-scores. (**Figs. S52-S53**)

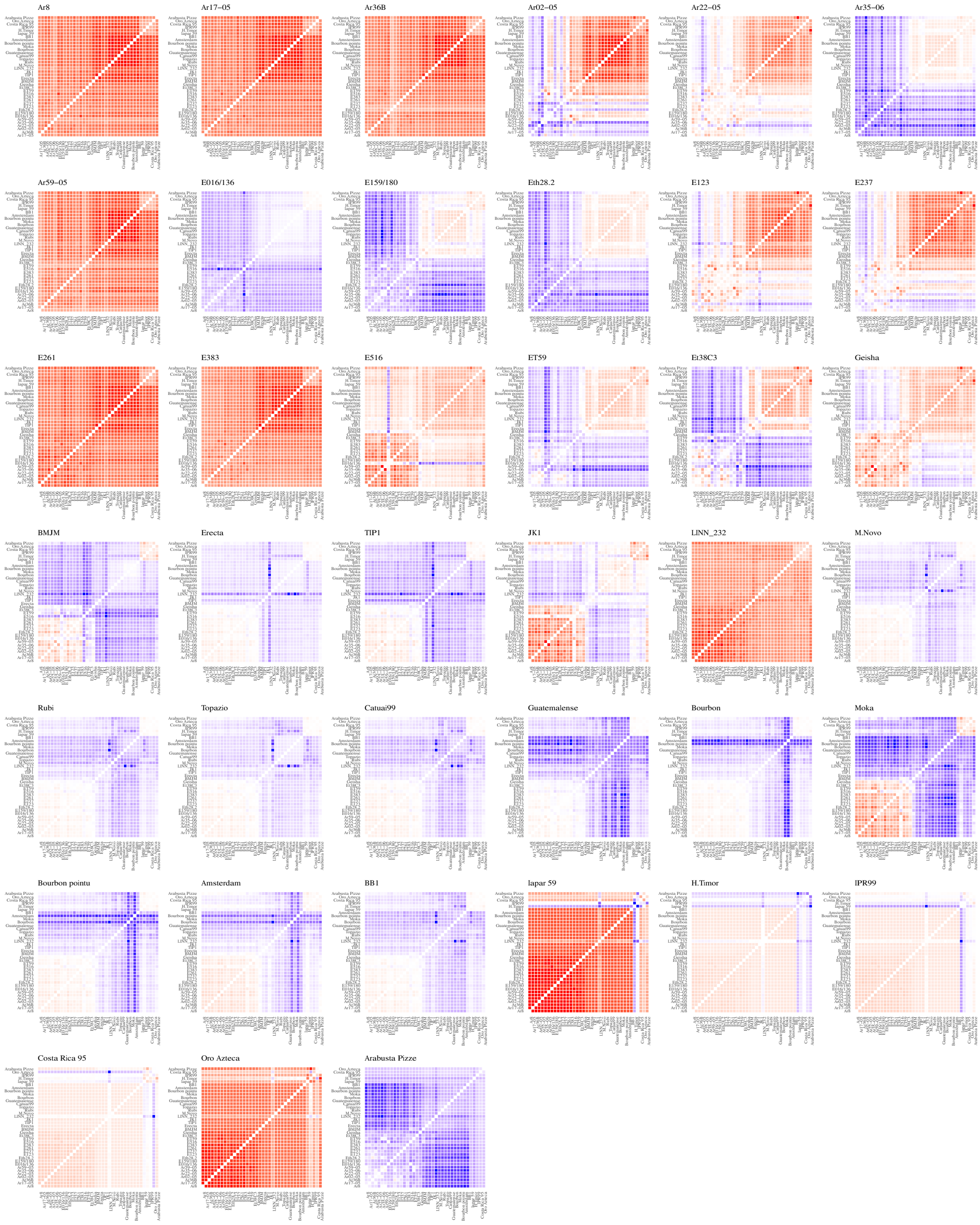


***Fig. S52.*** *subCC F3 statistics calculated for a target (the target name shown in top left of the matrix; possible sources are shown in horizontal and vertical axes). The result is illustrated with the color in the matrix cell; purple-blue color indicates a significant introgression test result (or close infrapopulational relationship), red a positive value, whereas white is non-significant.*

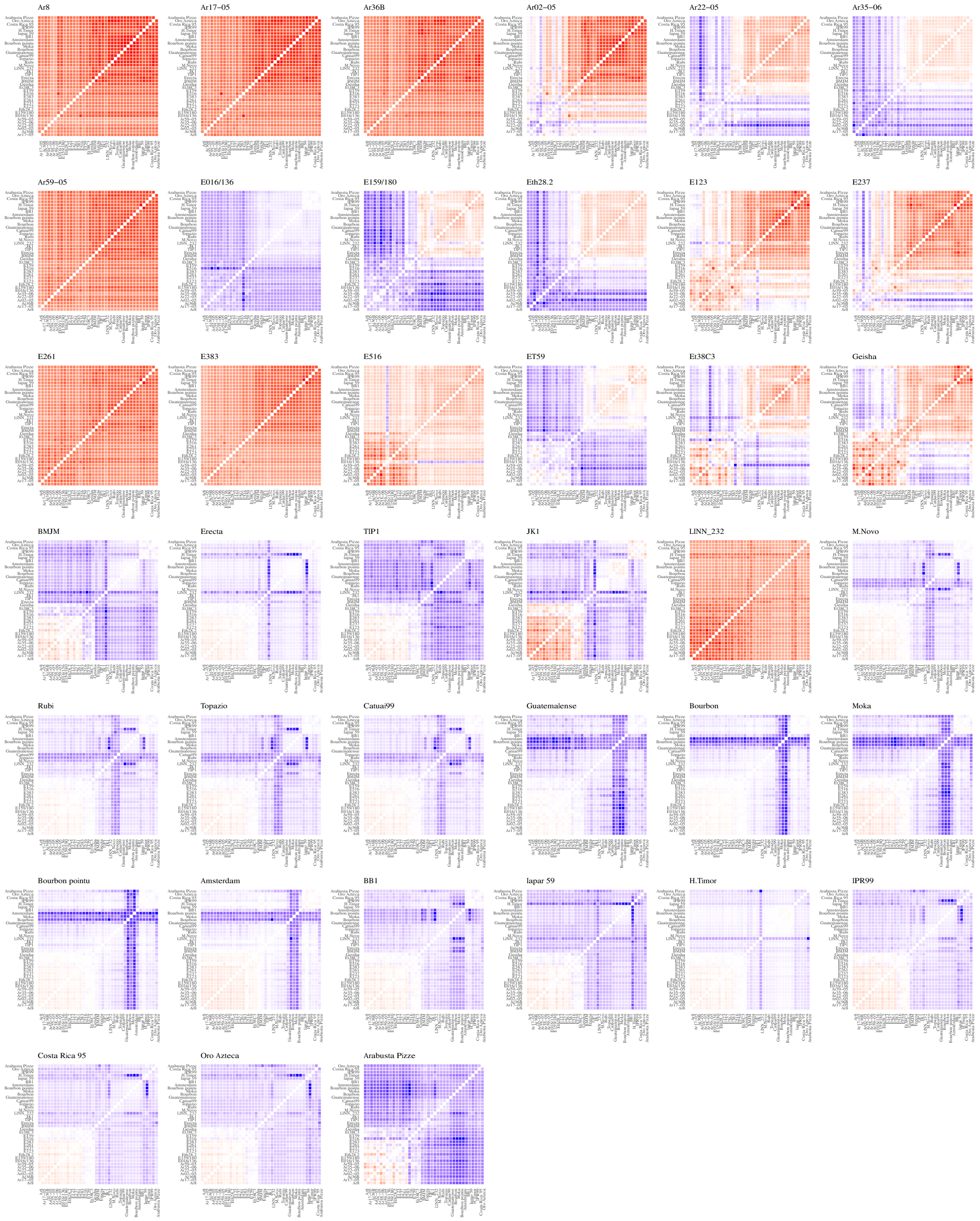


***Figure S53.*** *subEE F3 statistics calculated for a target (the target name shown in top left of the matrix; possible sources are shown in horizontal and vertical axes). The result is illustrated with the color in the matrix cell; purple-blue color indicates a significant introgression test result (or close infrapopulational relationship), red a positive value, whereas white is non-significant.*

##### 6.4.6 Introgression analyses

Orientagraph v1.0 (Molloy et al., 2021) was run for each of the subgenomes separately according to the developer recommendations by carrying out filtering for linkage as recommended for TreeMix (Pickrell & Pritchard, 2012).

The PopGenome R package (Pfeifer et al., 2014) was used to calculate $d_f$ statistics (Pfeifer & Kapan, 2019). For the analyses to detect possible admixture from cultivars into the wild admixed population, the analyses were carried out for the subEE genome. *Coffea eugenioides* accession DA56 was used as the outgroup in the analyses, and Typica accessions BMJM, Erecta and TIP1 were used as possible sources of introgression. Each individual from the admixed wild population was used as a putative target of introgression separately, and the non-admixed wild individuals were used as the non-admixed population. The results were visualized using scripts in R (**Fig. S54**).

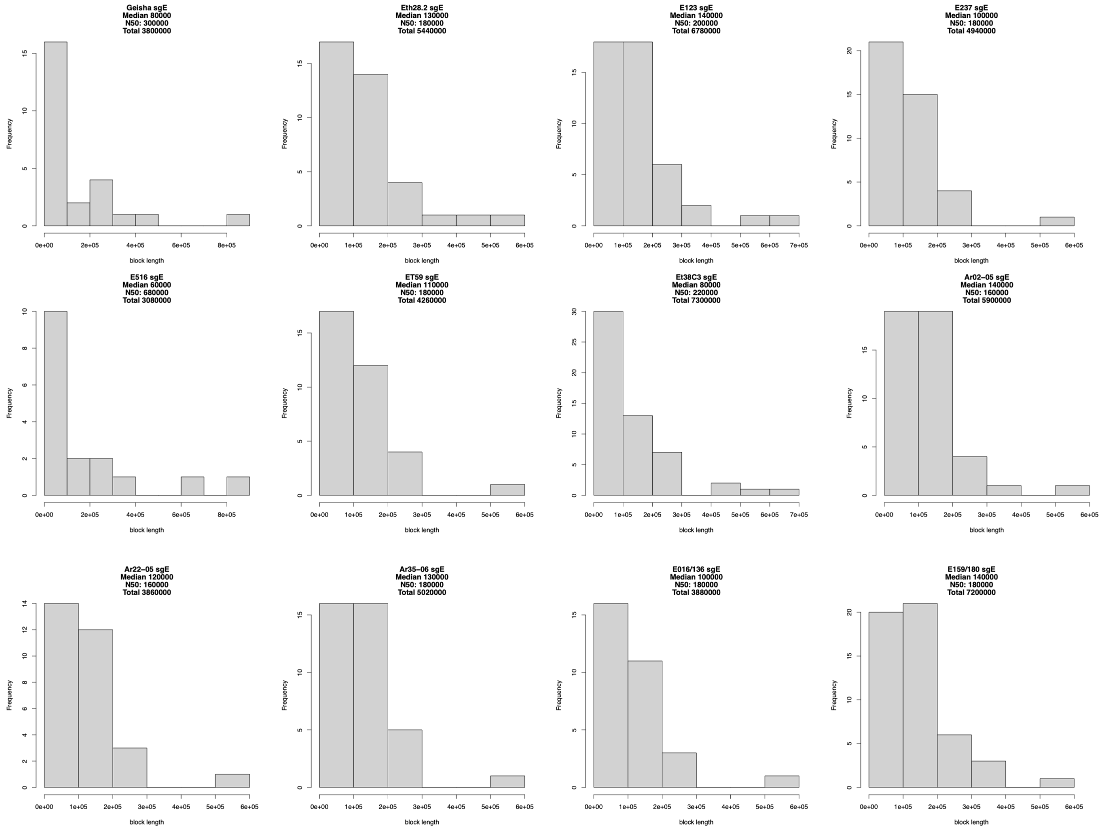


***Fig. S54.*** *The lengths of Typica-introgressed blocks in wild Coffea accessions.*

For the analyses to detect introgression from *C. canephora* in Timor hybrids, for subEE, *C. canephora* accession BUD15 was used as outgroup, *C. eugenioides* accession DA56 as the source of introgression, and E383 as a non-admixed wild representative. For subCC, DA56 was used as outgroup and BUD15 as the source of introgression. The statistic was calculated in 20 kb non-overlapping windows using a weighted jackknife to assess the significance of introgression. The results were visualized using R (**Fig. S55**).

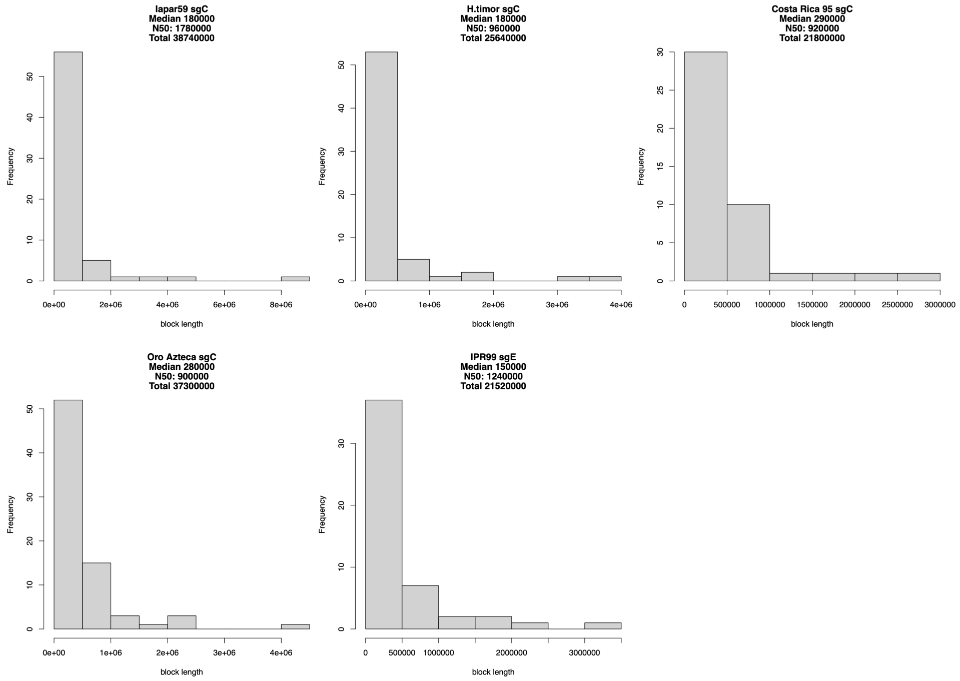


***Fig. S55.*** *The lengths of* Coffea canephora *introgressed blocks in Timor hybrid accessions.*

##### 6.4.7 Simulation

To gain more insight into population splits, we modeled Arabica population history with approximate Bayesian computation using FastSimcoal2 v2.6 (Excoffier et al., 2021). A site frequency spectrum (SFS) was calculated using angsd with the VCF file containing wild individuals and repetitive regions filtered out. For two populations, easySFS was used to obtain two-dimensional SFS. The ancestral states were estimated as described above. The initialization parameter file for the FSC was set to follow the overall trajectory observed for SMC++. Three different models were considered: (i) a model for only the non-admixed wild individuals (**Fig S56**), (ii) a model with wild individuals and cultivars and a population split, and (iii) a model with wild accessions and cultivars, with migration between the two populations after the split (**Figure 3E**). Identical models were run for the two subgenomic data sets.

For each of the models, 100 parameter files were simulated. For each parameter file 1,000,000 simulations were run; monomorphic sites were not used. Maximum composite likelihood estimation of parameters was carried out with 40 expectation-conditional maximization iterations. The model with the best likelihood was selected as the representative of the scenario.

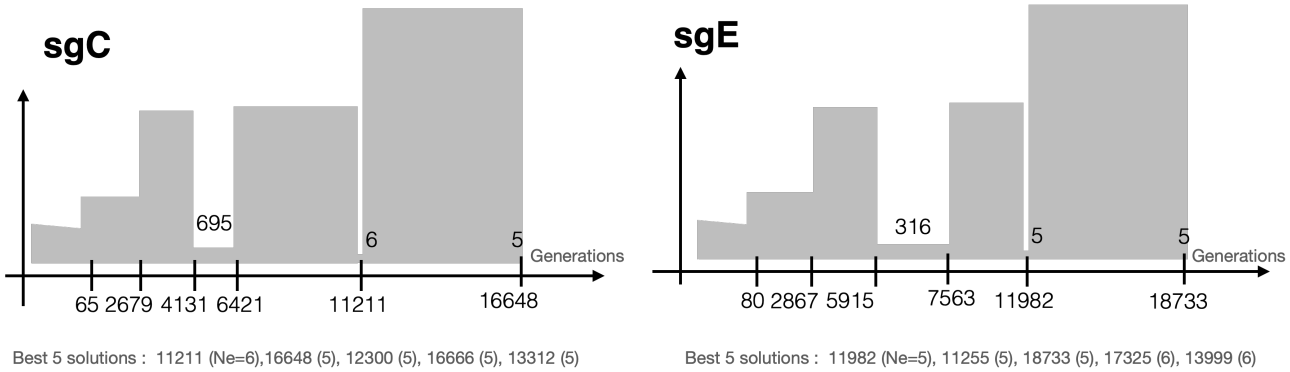


***Fig. S56****. Best models according to FatSimCoal for subCC and subEE and the wild, non-admixed population.*

In the best-fitting model (**Fig. 3E**), the wild population, including the admixed wild individuals, was predicted to split from the cultivar founding population at 1,450 generations ago (~30 kya), while the two populations maintained some gene flow (in terms of migration) until ~8-9 kya; this may have contributed to the modeled increase in effective population size between 8-20 kya (**Fig. 3D-E**). When compared against major climatic events, the wild vs. cultivated population split was predicted to occur before the latest glacial maximum (20-27 kya), with migration maintained until the end of the AHP.

To look for evidence of the original Arabica speciation event we also modeled older population bottlenecks. The wild population displayed strong coalescence at three different time points, with the first two at 11,200 and 17,700 generations ago. The oldest bottleneck, identified independently in both subgenomes, suggested coalescence to an extremely small population around 16,648-18,733 generations ago (350-393 kya); this also corresponds with the "flatlining" of all the SMC++ results (indicating full coalescence). Since subEE has experienced homoeologous exchange events which might inflate the time estimates, we suggest 350 kya to be a more reliable estimate.

Altogether, 610 kya is close to some previous estimates for the allopolyploidy event in recent literature (Bawin et al., 2020; Yu et al., 2011), the crown age of the *C. arabica* accessions in the subCC SNP tree (**Fig. S45**), as well as divergence estimates based on gene fractionation and the distribution of non-synonymous mutations (**Fig. S31**).

##### 6.4.8 Fixation index

Site-wise $F_{ST}$ values between wild and cultivated individuals were calculated for each gene annotation and 2kb flanking regions using VCFtools. Then, mean $F_{ST}$ values were calculated for each gene model using R.

### 7 Transposon Insertion polymorphisms

The *Coffea* genus has a narrow diversity (genetic pool), probably due to the bottleneck of the allotetraploid origin of this species. Our analyses revealed that the majority of the *C. arabica*, *C. canephora* and *C. eugenioides* genomes is composed of transposable elements (57%, 61% and 56% respectively). The vast majority of annotated transposable elements (TEs) falls into the Long Terminal Repeat (LTR) retrotransposon order (Copia and Gypsy, Supplementary data section 2.2). Among them, the Gypsy superfamily and more particularly the Del/Tekay family are the more abundant TEs. These elements have the potential to alter the expression of genes via insertion near or within them. So far, the contributions of transposable elements to *C. arabica* genetic and phenotypic diversity have not been investigated. Here, via the analysis of 39 *C. arabica* re-sequenced accessions from wild, cultivated and introgressed individuals, we investigated the impact of LTR retrotransposon insertions on *C. arabica* diversity.

We studied LTR retrotransposon insertions via analysis of short read whole genome resequencing data using TIP_finder (Orozco-Arias et al., 2020), using the discordant mapping pair approach. Briefly, Illumina read pairs were first mapped against the *C. arabica* reference LTR retrotransposon database annotated from the sequenced genome (ET39 wild individual). Discordant mapping pairs (one unique read that mapped to the reference TEs) were further processed for alignments against the reference *C. arabica* genome sequence. The uniquely mapped reads were placed in windows of 10 kb, which were then considered as the TE insertion location (with a minimum 10 mapped reads per insertion). From the 37 resequenced accessions, we selected a subset of 27 accessions for which classification as wild, cultivated or introgressed was unambiguous based on STRUCTURE data. Groups were constructed as follows: Wild: Ar8, CCCA33, Ar36B, FAO-0205, FAO-2205, FAO-3506, FAO-5905, E016/136, E159/180, Geisha, Eth28.2, E383, E51, TIP1; cultivated: M. Novo, Bourbon, Rubi, Topazio, Bourbon pointu, Catuaï99, Guatemalense, BB1; Introgressed group 1: Iapar 59, H. Timor, IPR99; and Introgressed group 2: Costa Rica 95, Oro Azteca. Results of insertions (a matrix of presence / absence per location) were processed with R v.3.6.3 (R_Core_Team, 2021) in order to run a discriminant analysis of principal components (DAPC) with the Adegenet package v. 2.1.3 (Jombart et al., 2010). The remaining ten additional accessions, not clearly assigned into the above genetic groups, were also analyzed by projecting them onto the discriminant axis found with DAPC, using the predict function from Adegenet. Circular visualizations of retrotransposon insertion locations were produced with ShinyCircos (Yu et al., 2018).

Insertion polymorphisms were mapped along the *C. arabica* pseudomolecules using TIP_finder (Orozco-Arias et al., 2020). In total 15,390 TIPs were identified, composed of 4,073 Copia TIPs and 11,317 Gypsy TIPs. The insertion frequencies of *Copia* and *Gypsy* LTR retrotransposon superfamilies was higher for wild accessions and introgressed (group 1) accessions, and lower for cultivated accessions and Introgressed group 2 (**Fig. S57**). Similar observations were made for maize accessions (Zhang & Qi, 2019). A large number of the TIPs overlap with gene regions (**Fig. S58**). TIPs were distributed along pseudochromosomes of *C. arabica* and follow the distribution of LTR retrotransposons (**Fig. S59**).

Wild, introgressed, and cultivated groups distinguished above using SNPs were also separated by a discriminant analysis of principal components - DAPC analysis (Jombart et al., 2010), based on the presence/absence LTR retrotransposons insertion polymorphism matrices for each sub-genome and superfamilies analyzed. Independently of the analyzed sub-genome or superfamily, the first discriminant axis (DA) distinguished clearly introgressed group 1 accessions from others while the second axis distinguished cultivated versus wild accessions (**Figs. S60-S63**).

Copia subCC Copia subEE Gypsy subCC Gypsy subEE

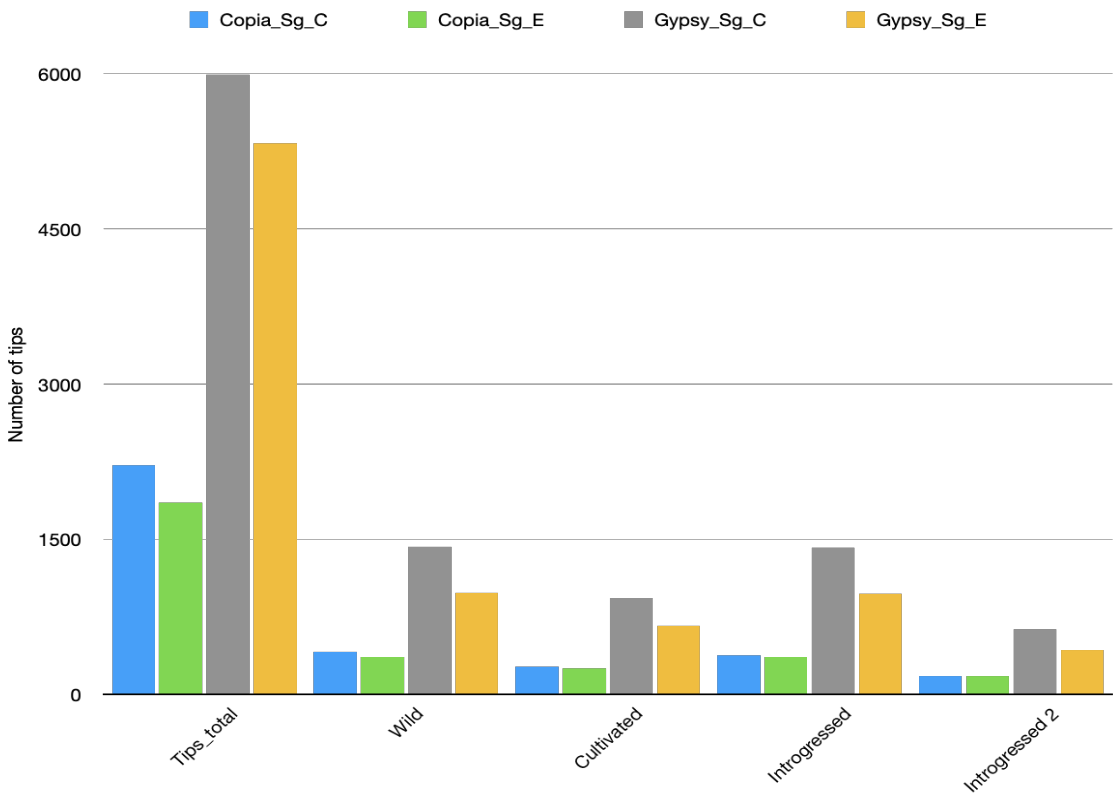


***Fig. S57.*** *Number of TIPs along* Coffea arabica *subCC and subEE per selected groups of accessions and per order of LTR retrotransposon (Copia, Gypsy): Total, Wild, Cultivated, Introgressed groups 1 and Introgressed groups 2.*

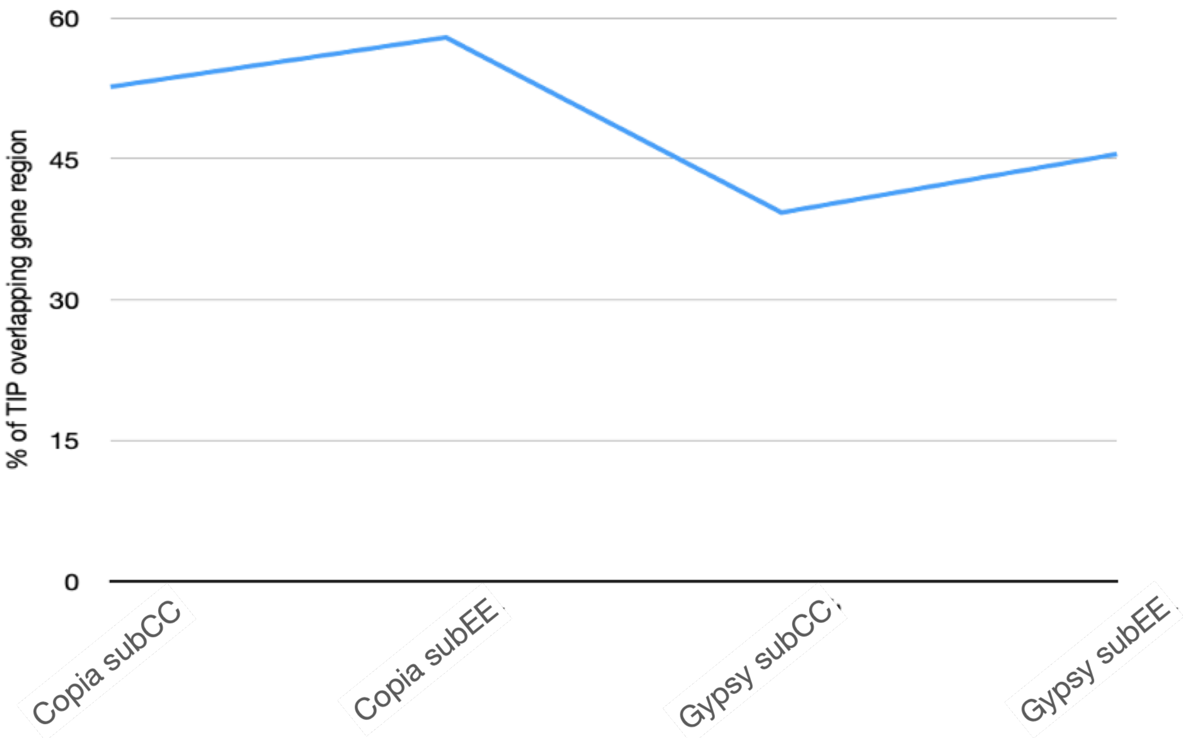


***Fig. S58.*** *Percentage of TIPS (window of 10 kb) overlapping gene annotations in* Coffea arabica *subCC) and subEE.*

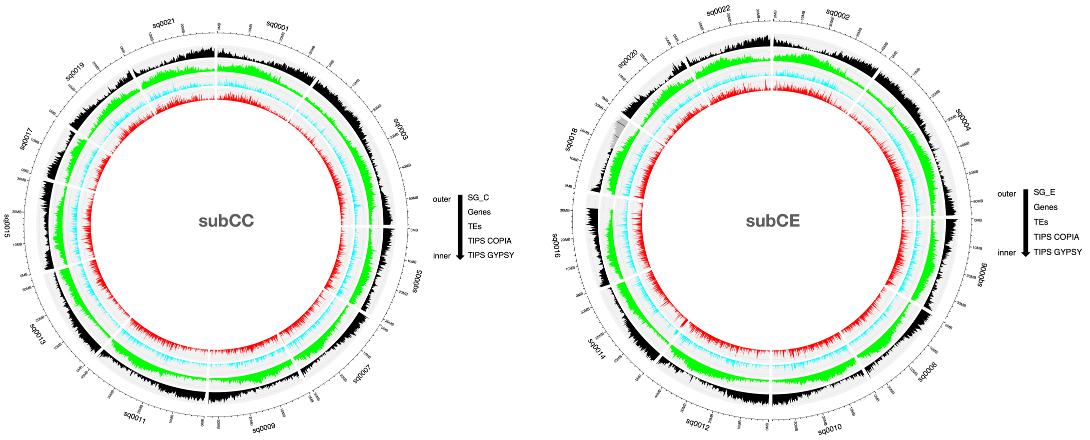


***Figure S59.*** *Landscape of TIPS along* Coffea arabica *subCC and subEE. Circos plots show the distribution of genes (black), Transposable elements (green), TIPs of LTR retrotransposons Copia (blue) and TIPS of LTR retrotransposons Gypsy (red).*

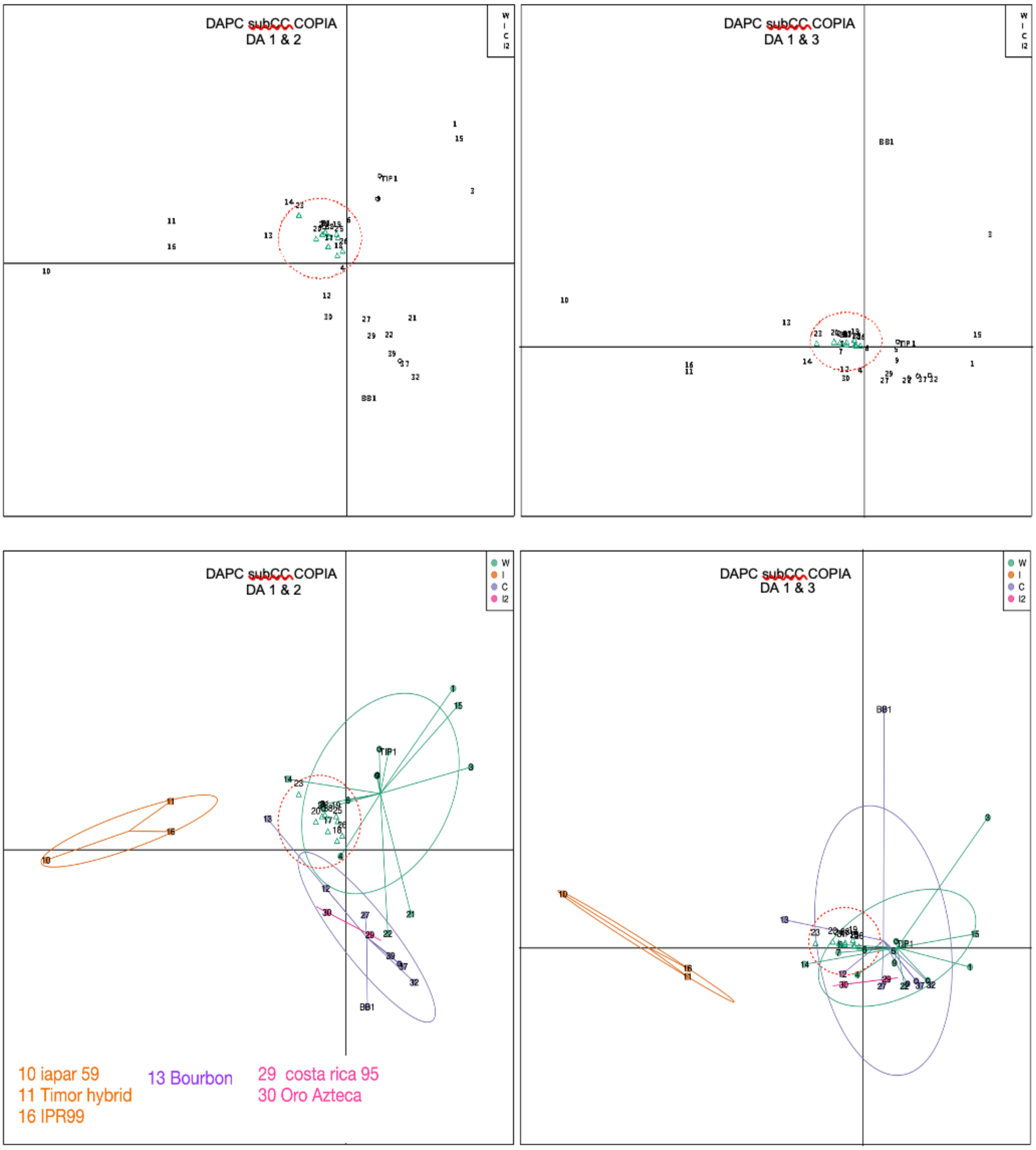


***Fig. S60.*** *DAPC (Axes 1 & 2, and axes 1 & 3) of Copia insertion polymorphisms on* Coffea arabica *subCC. The groups shown are wild (green), Introgressed group 1 (orange), cultivated (violet) and Introgressed group 2 (purple). Non-assigned accessions are presented and grouped by a red circle. 13 and BB1: Bourbon, 14: Geisha, TIP1 : Typica.*

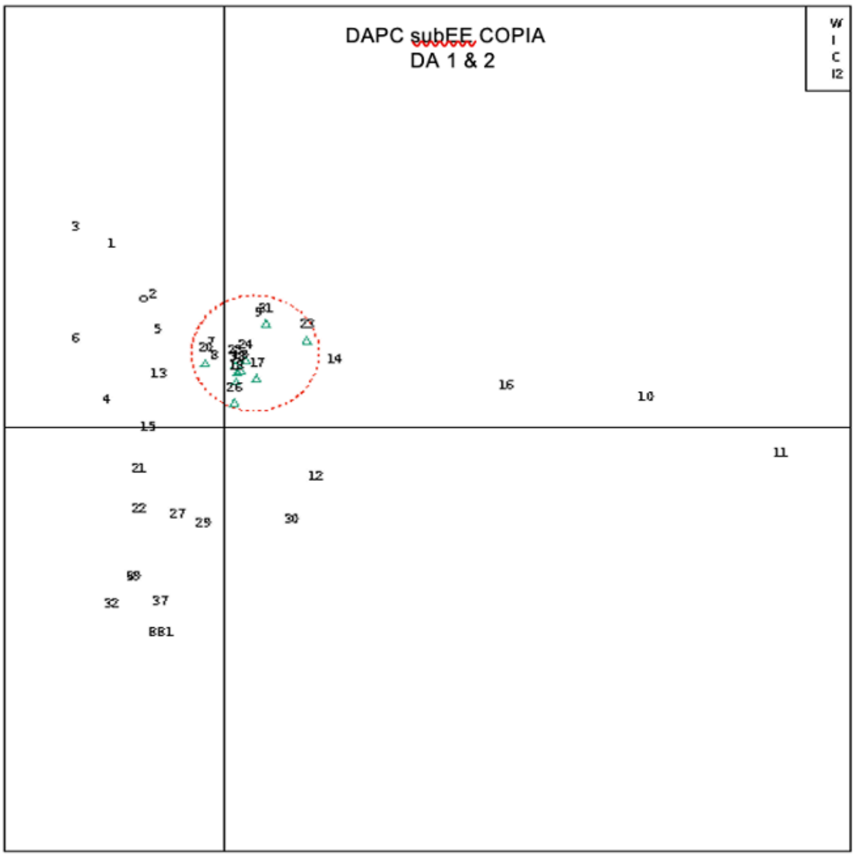
DAPC subEE COPIA
DA 1 & 2
W
I
C
I2

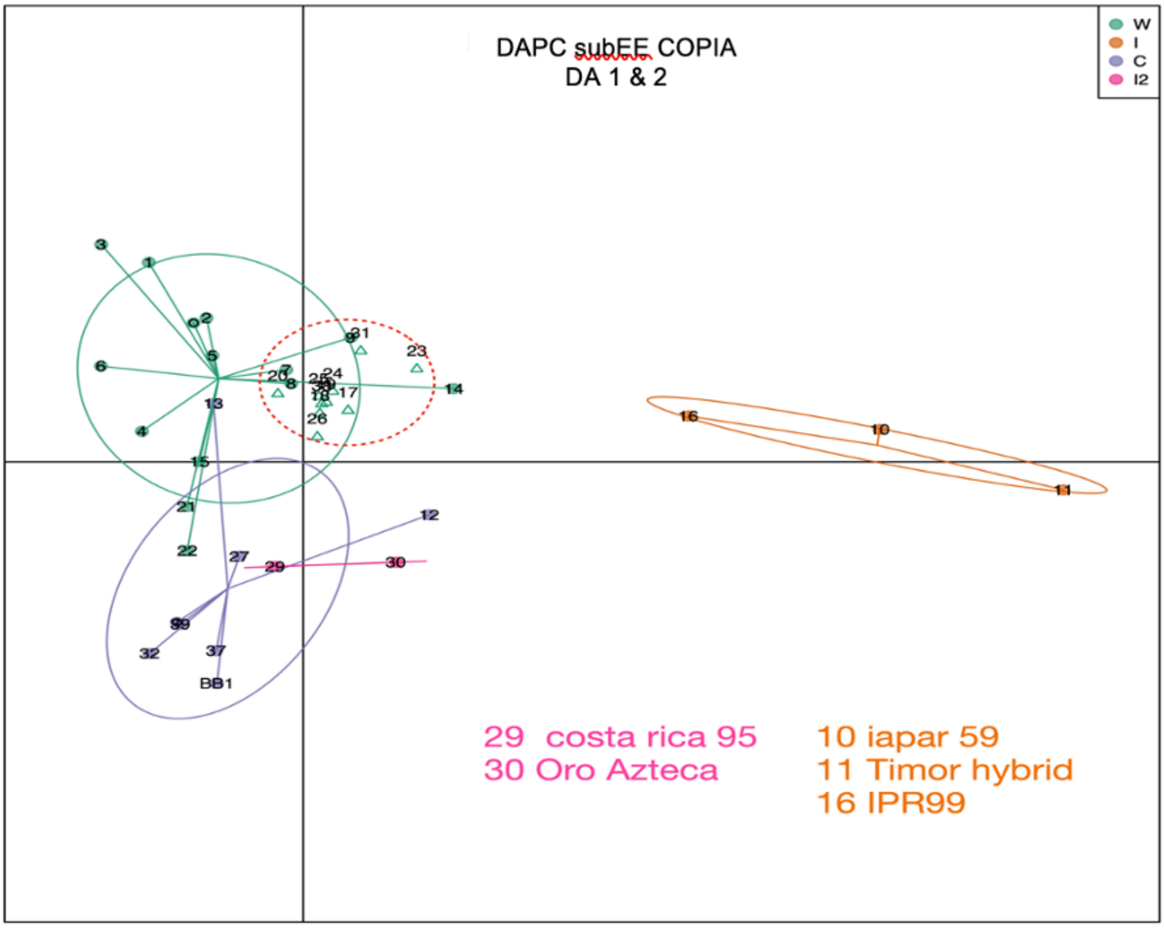
DAPC subEE COPIA
DA 1 & 2
W
I
C
I2
29 costa rica 95
30 Oro Azteca
10 iapar 59
11 Timor hybrid
16 IPR99

*Fig. S61. DAPC (Axis 1 and 2) of Copia insertion polymorphism on* Coffea arabica *subEE. The groups shown are wild (green), Introgressed group 1 (orange), cultivated (violet) and Introgressed group 2 (purple). Non-assigned accessions are presented and grouped by a red circle.*

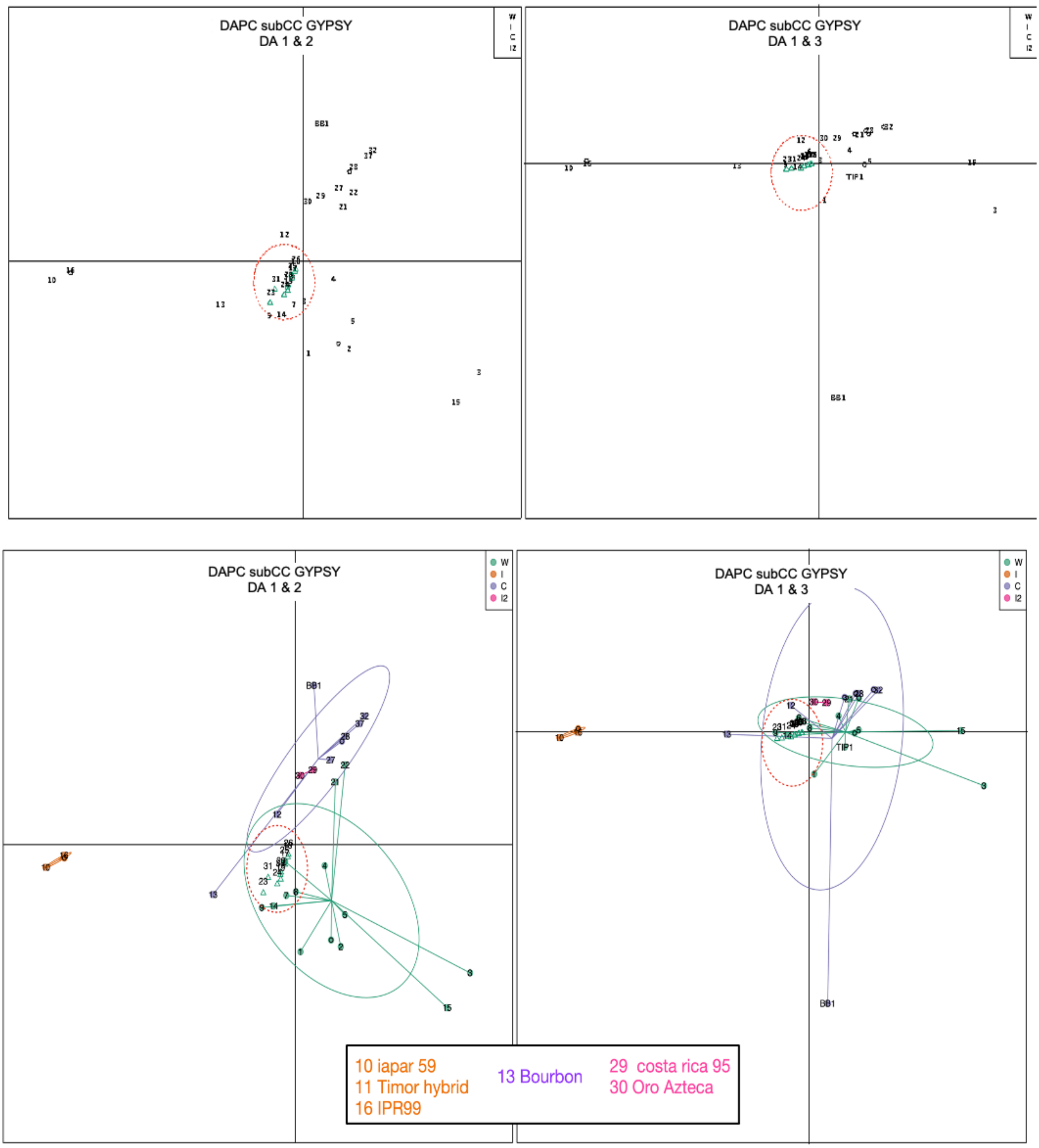


***Fig. S62**. DAPC (Axes 1 and 2, and axes 1 and 3) of Gypsy insertion polymorphisms on* Coffea arabica *subCC. The groups shown are wild (green), Introgressed group 1 (orange), cultivated (violet) and Introgressed group 2 (purple). Non-assigned accessions are presented and grouped by a red circle.*

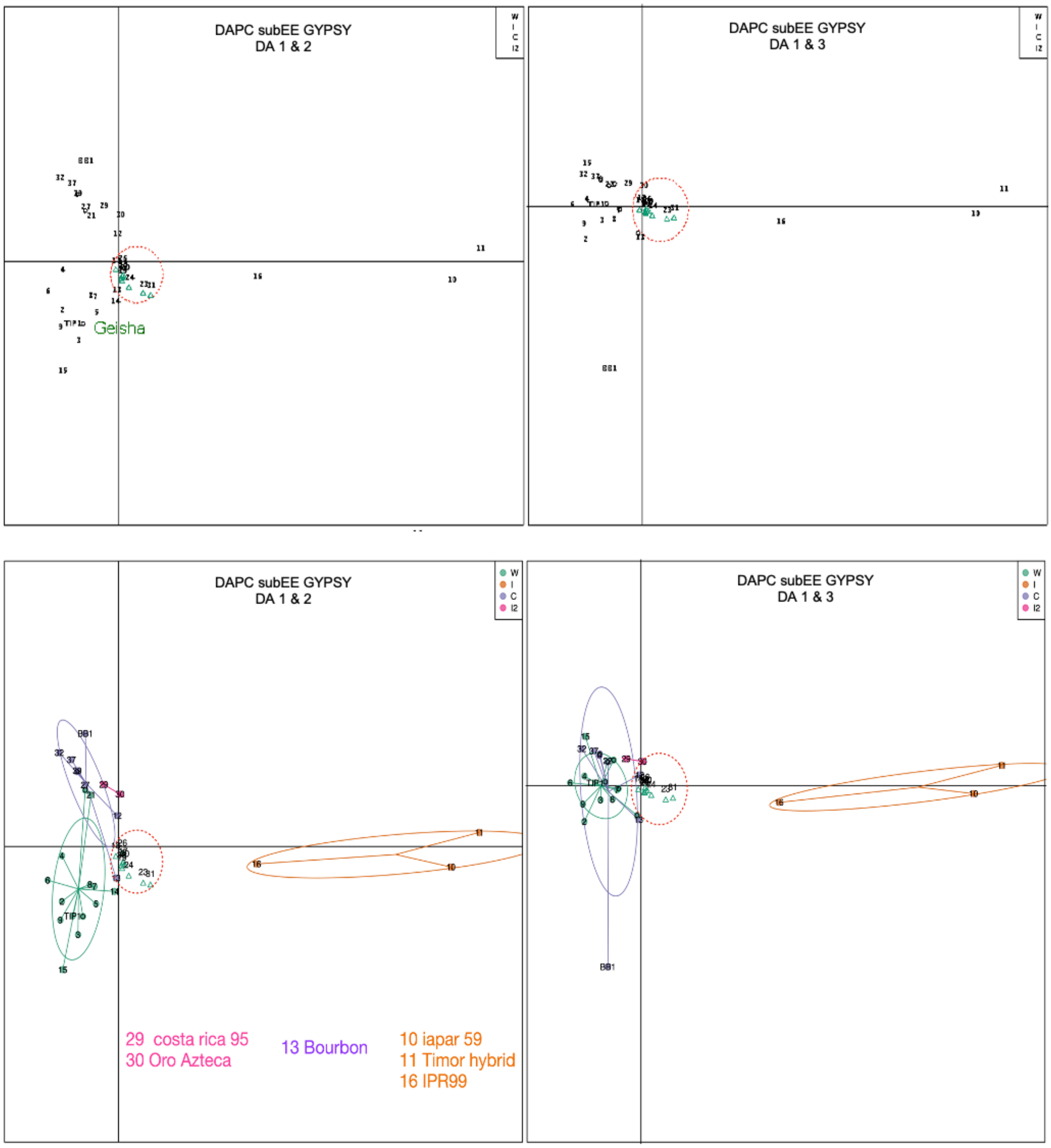


***Fig. S63.*** *DAPC (Axes 1 and 2, and axes 1 and 3) of Gypsy insertion polymorphism on* Coffea arabica *subEE. The groups shown are wild (green), Introgressed group 1 (orange), cultivated (violet) and Introgressed group 2 (purple). Non-assigned accessions are presented and grouped by a red circle.*

Retroelement insertions have been shown to play a role in gene regulation and thereby plant diversity, and for example in tomato, they have been found to be associated with variation of agronomic traits(Domínguez et al., 2020). The five Timor hybrid individuals permitted quantification of the distribution of Transposon Insertion Polymorphisms (TIPs) in individuals with recent introgression. Overall, we identified 157 Gypsy and 63 Copia insertions unique to the Timor hybrids; only 13 Gypsy and 4 Copia elements were shared by at least two accessions, testifying to their recent origins and dynamic nature. The TIPs showed a significant overlap with regions that were targets of introgression in different hybrids (Gypsy p=0.0002107; Copia p=0.03527; **Fig. 4B**). Similar to all insertions, the Timor hybrid-specific TIPs significantly overlapped tandemly duplicated gene regions (Gypsy $p$<2.2e-16; Copia $p$=3.99e-06).

#### 7.1 Biosynthetic gene clusters.

Biosynthetic gene clusters were identified with the Plantismash web server (http://plantismash.secondarymetabolites.org/). The tool identifies biosynthetic gene clusters (BGCs) by first finding genes that encode enzymes along biosynthetic pathways; a BCG is then defined as a locus where enzymes contributing to at least three different types of reactions are encoded. To find BGCs with three different types of reactions, and to separate them from tandem duplications, the genes at a putative locus are clustered to groups with >50% amino acid similarity; if three or more distinct clusters remain, the locus is a BGC (Kautsar et al., 2017).

The tool identified 95 BGCs in the *C. arabica* genome assembly (**Table S33**), including 7 alkaloid, 17 terpene and 27 saccharide gene clusters. Their functions are yet to be determined, but overall, BCGs have been found to play important roles in plant specialized metabolism (Polturak & Osbourn, 2021).

Many plant secondary metabolites are synthesized by genes arranged in biosynthetic gene clusters, a set of co-located genes contributing to the same pathway(Polturak et al., 2022; Polturak & Osbourn, 2021). Within the 39 accessions, the TIPs significantly also overlapped genes residing in biosynthetic gene clusters ($p$<2.2e-16; Fisher exact test). We similarly found TIPs in the Timor hybrid-derived material near genes of the *FAD2* and *TPS* families; it remains to be ascertained, however, whether these TIPs affect sensory characteristics in these cultivars.
