## Extended data figures for "The genome and population genomics of allopolyploid *Coffea arabica* reveal the diversification history of modern coffee cultivars"

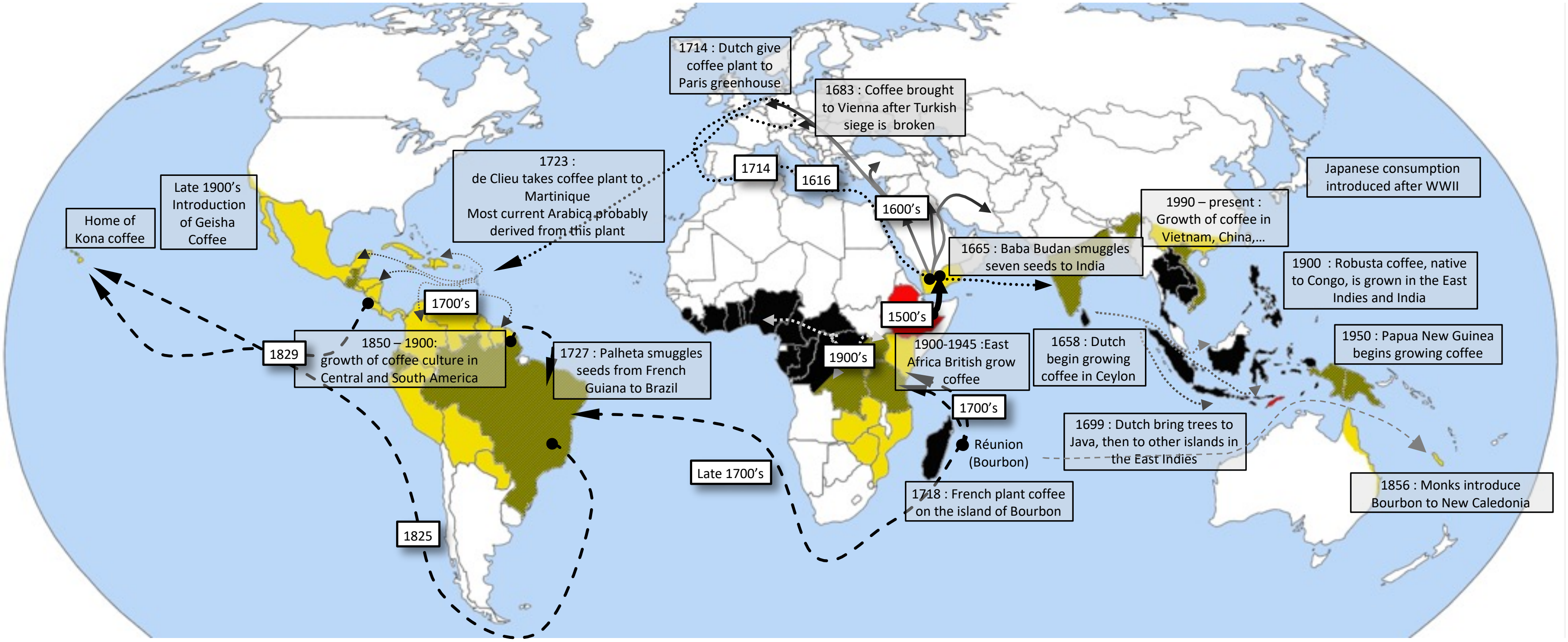


**Ext. data Fig. 1.** *Coffee dissemination routes. Yellow: current* Coffea arabica *cultivation; black: current* C. canephora *cultivation; ochre: current cultivation, both species. Solid lines: early spread of coffee consumption; dashed lines: main Bourbon routes; dotted lines: main Typica routes; Ethiopia (the center of origin of Arabica) and Timor Island (the origin of the Timor hybrid) are colored in red.*

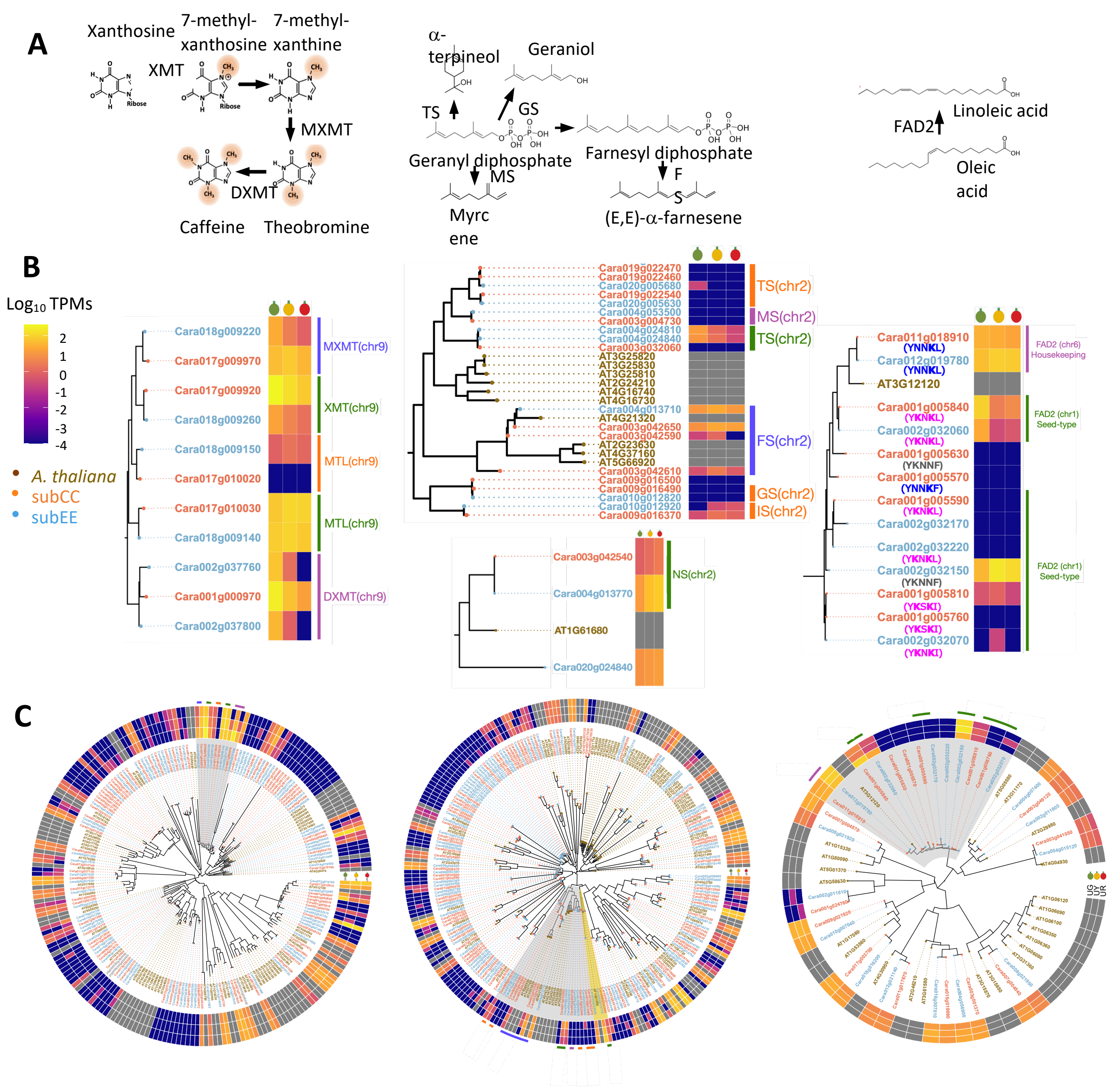


**Ext. data Fig. 2.** *Composition and expression of* exemplar Arabica *gene families contributing to bean quality traits.* ***A****. Schematic biosynthesis of caffeine (left), terpenoids (middle), and unsaturated fatty acids (right)* ***B.*** *Phylogenies and expression during fruit development of* CA *genes for N-methyltransferases (NMTs) mediating caffeine biosynthesis (left), terpene synthases (TPS) (middle), and fatty acid dehydrogenase 2 (FAD2) (right). RNA sequencing was carried out for three biological replicates from three different fruit maturation stages (green, yellow, and red) of the K7 cultivar.* ***C.*** *Genome-wide NMT (left), TPS (middle), and FAD2 (right) phylogenies and expression patterns during fruit development. Genes located in the two subgenomes are indicated by font color; subCC (red) and subEE (blue). Grey areas highlight the parts of phylogenies shown in B. XMT: xanthosine methyltransferase; MXMT: 7-methylxanthine methyltransferase; DXMT: 1,7-dimethylxanthine methyltransferase; MTL: N-methyltransferase-like; FS: (E,E)-a-farnesene synthase; GS: Geraniol synthase; IS: Isoprene synthase; MS: myrcene synthase; TS: (-)-a-terpineol synthase; FAD2: Fatty acid desaturase 2.*

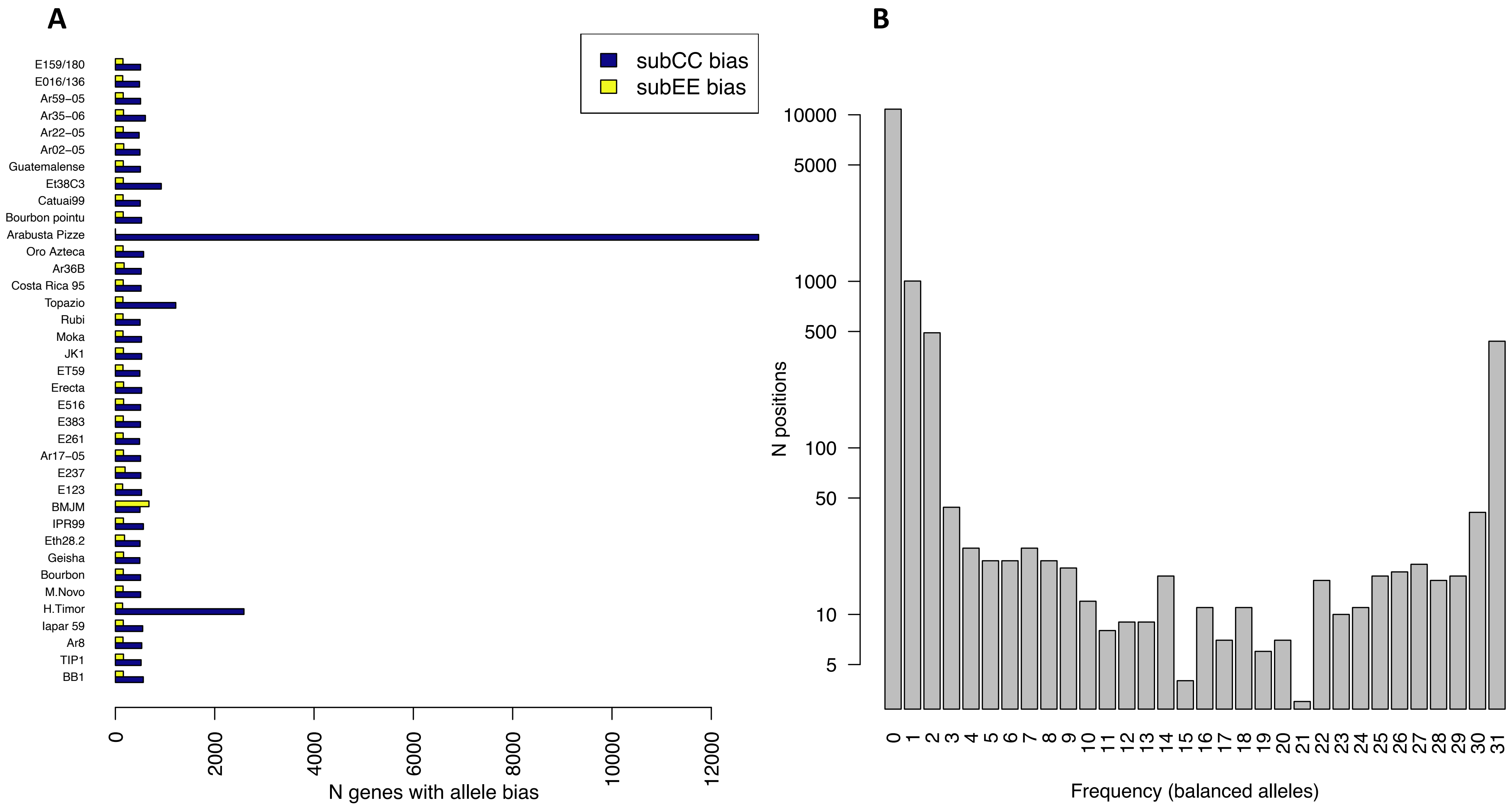


**Ext. data Fig. 3. A.** Summary of homoeologous exchange between subgenomes. Blue bars indicate genes with 3:1 allele bias towards subCC, whereas yellow bars indicate genes with allele bias (1:3 or 0:4) towards subEE. **B.** Frequency spectrum of shared homoeologous exchange from subCC to subEE.

**A** **B**

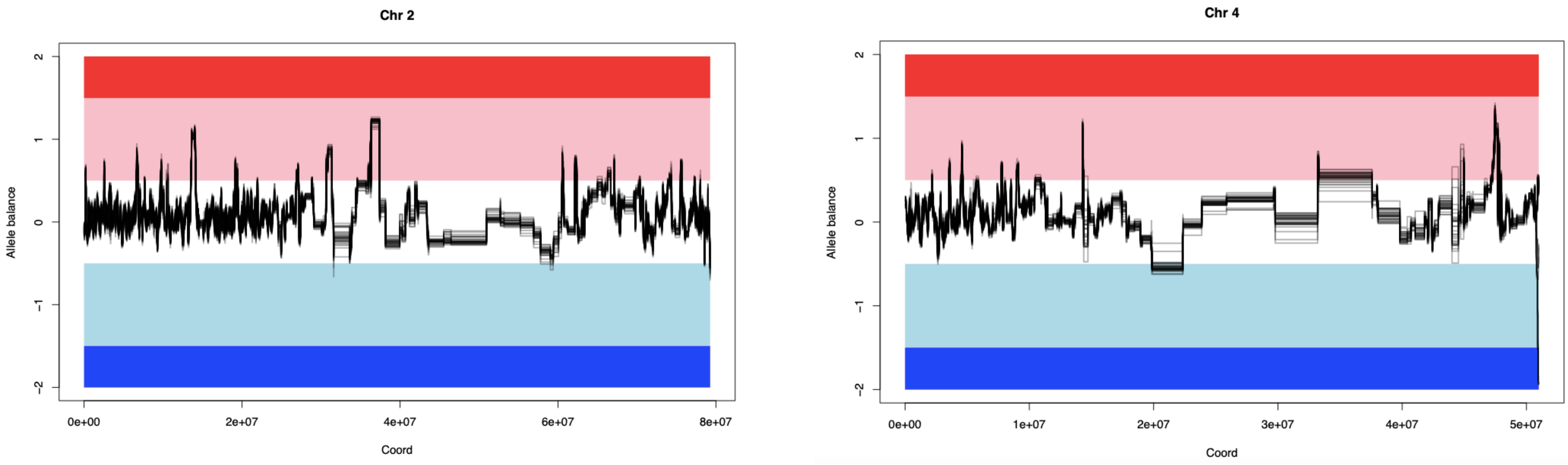


**Ext. data Fig. 4.** *Homoeologous exchange plots of chromosomes 2 (left) and 4 (right) overlaying all* Coffea arabica *accessions in this study. The dark red region indicates 4:0 allele balance in favor of subCC, while the pink region illustrates=3:1, white 2:2, light blue 1:3 and dark blue 0:4 balances, respectively. The gray lines indicate the observed allele balances in syntelog gene pairs for the different Arabica accessions. For a view of all chromosomes and of the genes involved, see* ***Fig. S38*** *and* ***Table S15.***

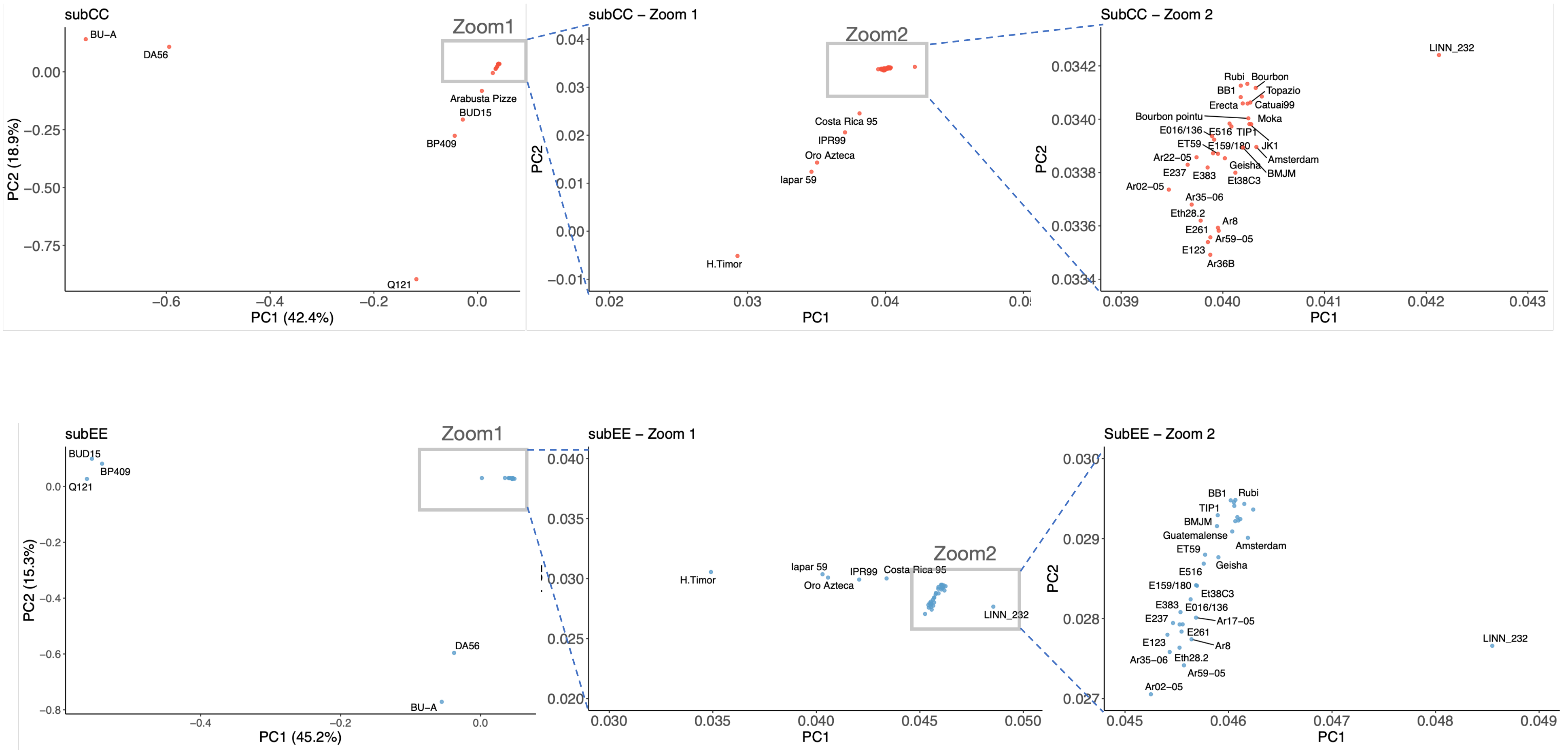


**Ext. data Fig. 5.** *Left: PCA plot based on the SNPs called in subCC (top) and subEE (bottom) subgenomes. The rectangles highlight the regions zoomed in the following plots.*

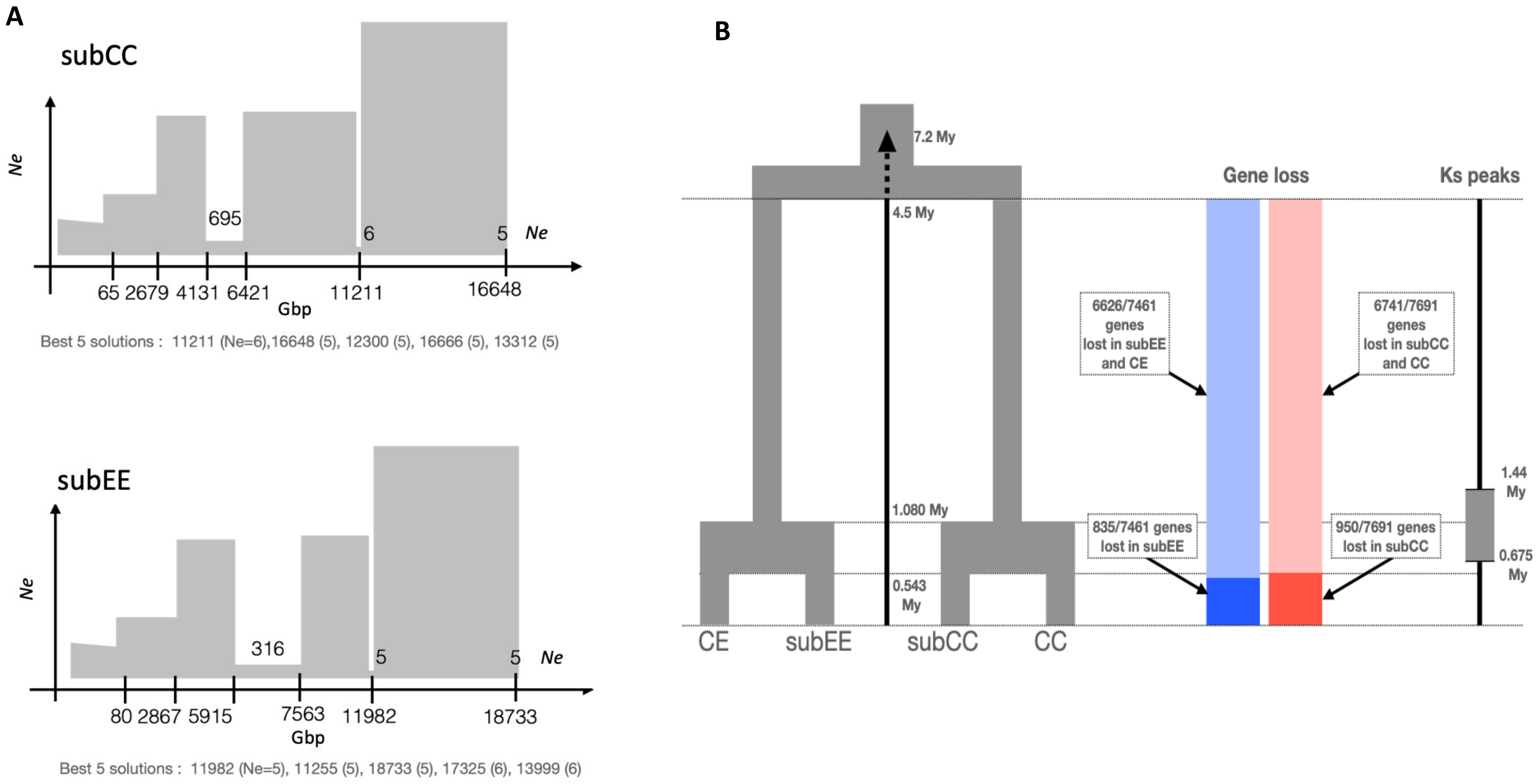


**Ext. data Fig. 6**. ***A.*** *Summary of FastsimCoal2 models for historical effective population sizes (Ne). The effective population size (y-axis) is plotted against the number of generations before present (Gbp, x-axis). The bottlenecks were identified with 100 FastSimCoal runs with 10^6 simulations. Maximum composite likelihood estimation of parameters was carried out with 40 expectation-conditional maximization iterations. The plots summarize the best models for subgenomes CC and EE in the wild, non-admixed population. To convert the generations to years, an estimate of 21 years/generation was used (Moat el al. 2019).* ***B.*** *Summary of the genome fractionation rate and divergence of syntenic gene models. The timing of the splits in the phylogeny (left) reflects the most recent estimates from (Bawin et al., 2020). The rate of gene loss (barplot) is presented as the percent of syntenic genes lost in the Eugenioides/subEE common ancestor (light blue) or only in subEE (blue). A similar analysis was carried out for Robusta-derived genomes, where the percent of genes lost in Robusta/subCC is shown in light red and genes lost only in subCC with dark red. The Ks peaks method (right) scales the divergence time between the subgenomes, estimated from numbers of synonymous mutations between syntenic genes to the timing of the speciation event.*

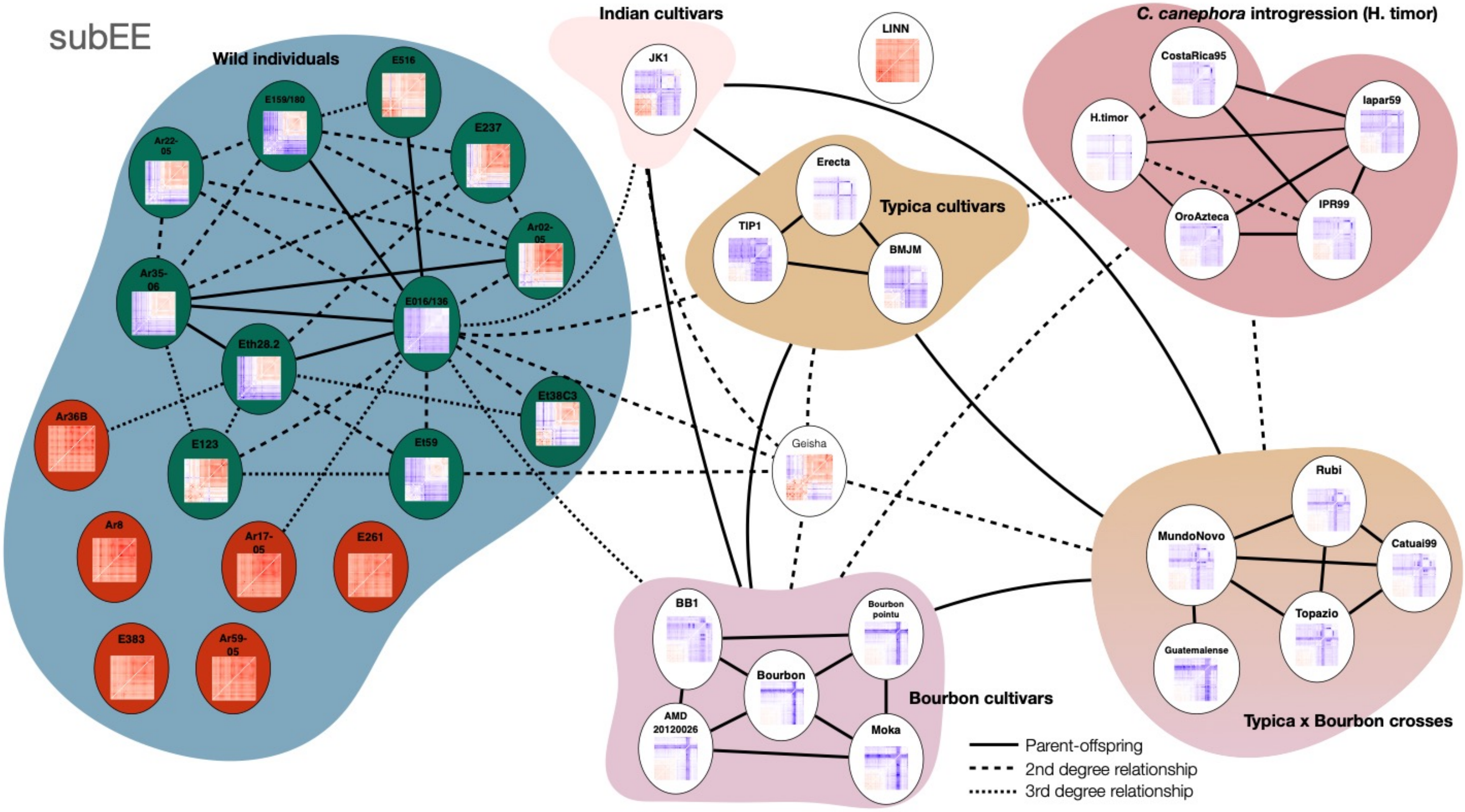


**Ext. data Fig. 7**. *Kinship analysis on subEE. The degree of relatedness was estimated using Kinship-based INference for GWAS (KING); Thumbnail images show false discovery rate corrected F3 tests of introgression Z-statistics for each of the target individuals; purple-blue color illustrates significant gene flow from the two sources (horizontal and vertical) to the target, or close infrapopulational relationship between sources and target. The green background in the wild accessions highlights the admixed individuals (Figure 3B); the non-admixed individuals are highlighted with red. The corresponding analysis on subCC is shown in* ***Fig. 4.***

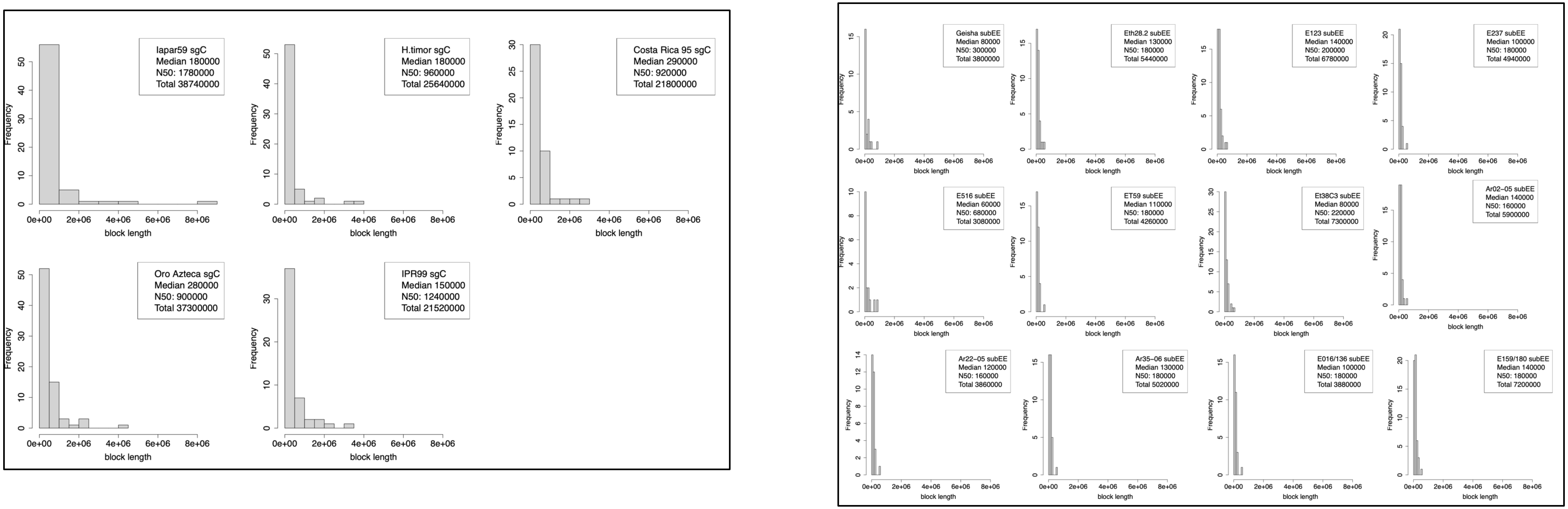


**Ext. data Fig. 8.** *Lengths of Robusta introgressed blocks in Timor hybrid accessions (left) and, as a control, of Typica introgressed blocks in wild Arabica accessions (right) .*

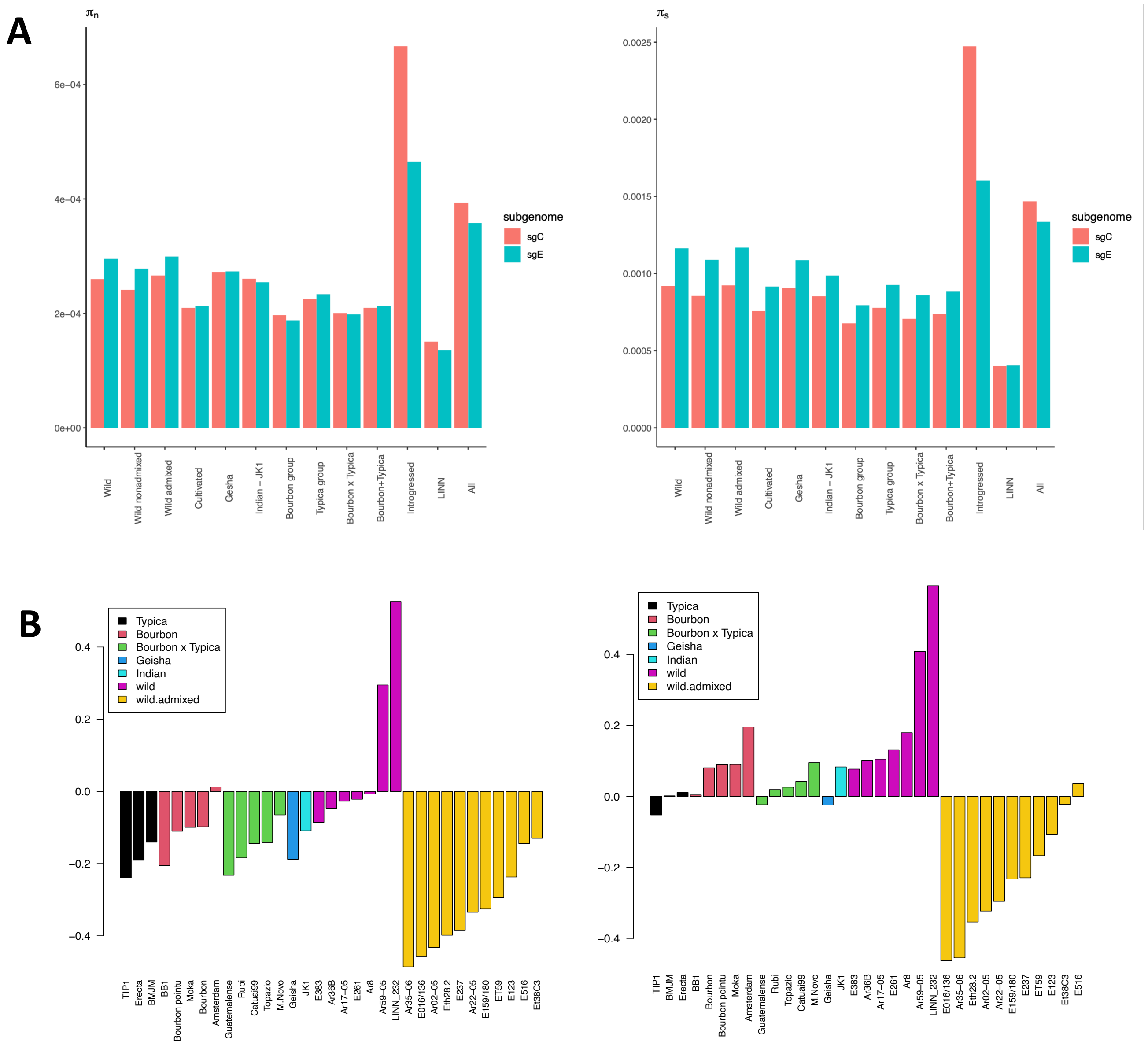


**Ext. data Fig. 9. A.** *Non-synonymous ($\pi_n$; left) and synonymous ($\pi_s$; right) diversity in the different populations of C. arabica.* ***B.*** *F inbreeding coefficients for wild and cultivated lines, shown separately for subCC (left) and subEE (right).*
